# Simple Causal Relationships in Gene Expression Discovered through Deep Learned Collective Variables

**DOI:** 10.1101/2023.01.18.524617

**Authors:** Ching-Hao Wang, Kalin Vetsigian, Chris Lin, Finnian Firth, Glyn Bradley, Lena Granovsky, Jeremy L. England

## Abstract

Developments in high-content phenotypic screening with single-cell read-out hold the promise of revealing interactions and functional relationships between genes at the genomic scale scale. However, the high-dimensionality and noisiness of gene expression makes this endeavor highly challenging when treated as a conventional problem in causal machine learning, both because of the statistical power required and because of the limits on computational tractability. Here we take different tack, and propose a deep-learning approach that finds low-dimensional representations of gene expression in which the response to genetic perturbation is highly predictable. We demonstrate that the interactions between genes that are cooperative in these representations are highly consistent with known ground-truth in terms of causal ordering, functional relatedness, and synergistic impact on cell growth and death. Our novel, statistical physics-inspired approach provides a tractable means through which to examine the response the living cell to perturbation, employing coarse graining that reduces data requirements and focuses on identifying simple relationships between groups of genes.

**Author summary:** Understanding the causal relationships between genes and the functions of a cell’s molecular components has long been a challenge in biology and biomedicine. With recent advancements in technologies that manipulate and measure the activity of thousands of genes at once at the single-cell level, scientists are now afforded with the opportunity to interrogate such relationships at scale. However, extracting useful information from the vast readouts of these technologies is non-trivial, in part due to their many-dimensional and noisy nature. Here we develop a machine learning model that allows for the interpretation of complex genetic perturbations in terms of a simple set of causal relations. By analyzing cooperative groups of genes identified by our model, we demonstrate the model can group genes accurately based on their biological function, their relative ordering up- or downstream in the flow of causation, and how their activities combine to affect cell growth and death. Our approach complements existing machine learning methods in providing a simple way to interpret causal mechanism governing genetic interactions and functional states of cells.

## Introduction

The development of high-throughput-omics technologies has opened up a broad frontier of new possibilities for characterizing and controlling biological systems at the molecular level. Recent advances in both the measurement and manipulation of gene expression through technologies such as scRNA-seq and CRISPR now provide the opportunity to treat the living cell as a high-dimensional dynamical system, which not only can be described in detailed quantitative terms, but also can be steered with controlled perturbations. This potent combination of capabilities hold significant promise both for novel therapeutic approaches and for a different way forward in understanding basic biology. Single-cell platforms such as Perturb-seq [1–7], which enables simultaneous modulation of multiple genes with a full transcriptomic read-out after perturbation, have already offered many new insights into the functional relationships of different proteins. Viewed as a predictive problem of mapping input to output, however, the Perturb-seq scenario presents significant challenges. The combinatorial vastness of the input space (with combined activations and repressions of tens of thousands of genes to choose from) ensures that most experiments that are possible in principle cannot be carried out. Meanwhile, the high-dimensionality, discreteness, and stochasticity of the measured quantities of transcripts for each gene may hide biologically interesting patterns in subtle, many-body correlations that are covered in a blanket of experimental and molecular noise.

Machine learning has already been applied successfully to address some of these problems [8]. The use of autoencoders [9], UMAP [10,11], and other methods of dimensionality reduction [12,13] have demonstrated that it is possible to discover relationships in statistical covariation of many genes that can be used, for example, to distinguish different cell types and lineages [14], to predict mean gene expression under perturbation [15,16], to predict the interactions between genes under combinatorial perturbations [17], and to generate counterfactual predictions on single-cell response under new drug treatments [18] or novel perturbations [19]. However, the greatest successes in the analysis and modeling of single-cell transcriptomes have principally been restricted to mapping the manifold where the observed data distribution tends to fall, and have done less to show how to use perturbations to navigate across it. Though it is possible in principle to learn causal relationships between individual genes from data, most approaches to this task fail, whether for lack of sufficient samples or due to computational slow-down, when trying to handle thousands of genes at once. Additionally, it remains an open question how easily system behavior can be predicted from even an accurate map of so many complex relationships, particularly when strong perturbations lead to non-linear, synergetic response. Thus, attempts at predicting the simultaneous changes in expression of thousands of genes following new combinations of cellular perturbations have so far met with difficulty.

In one sense, however, this difficulty is not a new one to quantitative science. More than a century ago, it was appreciated that the task of exactly predicting the behaviors of all the molecules in a gas or liquid was hopeless, but that there nonetheless were highly accurate predictive laws that did govern the collective, macroscopic properties of these substances. Thus, while it was not (and still is not!) feasible to keep track of what every molecule in a fluid is doing, many-to-one functions of molecular coordinates such as temperature and pressure have simple and reproducible relationships that grant tremendous predictive power. Taking inspiration from this ‘statistical mechanical’ framework, we now ask a similar question about where to find predictability in the cellular transcriptome. If individual genes are highly variable and extremely numerous, could there perhaps be a set of quantities computed from thousands of gene expression levels at once that comprise a low-dimensional space of simple, predictable response to perturbation?

Specific examples of such relationships are already well-known to biology through the concept of biochemical pathways. Whether in metabolism, the cell cycle, or regulation of programmed death, the activation or inhibition of processes involving many proteins working in concert is often a precisely regulated event, in which particular perturbations can trigger large collective changes in gene expression. Historically, pathways have been discovered by building outward from initial biochemical discoveries, until a web of functional relationships is uncovered. Alternatively, genes have long been clustered and grouped according to observed correlations in their expression or other measures of relatedness. While the first of these approaches is painstaking and incomplete, the second often does not imply a clear prediction about how a cell is expected to respond when a gene or group of genes is targeted. Yet, the past successes of pathway identification through biochemical experimentation and clustering suggest an opportunity to capture what is predictable about cellular response to perturbation in a unified framework that can be learned from panoramic data.

## Results

We develop Lowdeepredict, a deep learning model for single-cell gene expression under perturbation. It maps perturb-seq single-cell RNA sequencing data to a low-dimensional representation in which the response to single and multiple genetic perturbations is highly predictable. Since this is a neural network-based model, we devise an efficient procedure to impute causal relationship between genes (through a forward pass) and use that to generate hypotheses regarding genetic interactions and functional relatedness. We also demonstrate a Monte Carlo approach to identify novel gene sets with high synergistic interactions. We further show that the causal interactions between genes in these sets are preserved through a benchmarking scheme.

### Lowdeepredict learns a low-dimensional representation in which causal relationships between genes can be robustly detected

Uncovering the causal relationships between genes from single-cell RNA sequencing data from perturbation screens is a daunting task, due to long-standing challenges in causal inference for high-dimensional data [20–24]. Deep learning approaches present a possible way forward because they excel at projecting such data into low-dimensional spaces where relationships between the projected variables are easier to uncover and sometimes more interpretable [25]. We propose a novel deep learning model that identifies such low-dimensional representations in a perturbation-response setting. Unlike existing methods that focus on, for example, predicting the expression patterns for unseen perturbations [15–19] (see [8] for a summary of these methods), our approach *stipulates* a low-dimensional set of causal relationships and then *requires* neural networks to find new representations of gene expression that respect these relationships. Doing so allows us to robustly detect and predict changes of state in this low-dimensional space that result from perturbations.

In Fig. 1, we illustrate the ideas behind this scheme. Consider the expression of three disjoint gene groups {*b*_1_, *b*_2_,…,*b_N_b__*} (blue), {*g*_1_, *g*_2_,…, *g_N_g__*} (green) {*r*^1^, *r*_2_,… *r_N_r__*} (red) and neural networks *F_b_*, *F_g_*, and *F_r_* that map them to real numbers *z_b_*, *z_g_*, and *z_r_*, respectively. Different transcriptional states define points on the level hypersurfaces described by *z_b_* = *F_b_*(*b*_1_,*b*_2_,…,*b_N_b__*) in 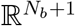, by *z_g_* = *F_g_*(*g*_1_,*g*_2_,…,*g_N_g__*) in 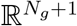, and by *z_r_* = *F_r_* (*r*_1_,*r*_2_,…,*r_N_r__*) in 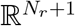. In Fig. 1, we assume that without any perturbation, the state of a cell is given by (*z_b_,z_g_,z_r_*) = (0, 0, 0) and that the *imposed* causal order is such that *z_b_* causes *z_g_* and *z_g_* causes *z_r_*. When gene *b_k_* is perturbed, e.g., being CRISPR activated to the expression level of 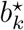 (i.e. expression is *quenched* to the value 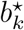), the cell state moves along the blue surface to where *z_b_* takes value 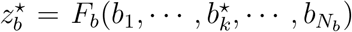 (i.e. the “height” of the surface is 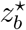, see Fig. 1**b**). Since *z_b_* is upstream of *z_g_* and *z_r_*, this response to perturbation is sequentially *propagated* to 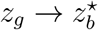 then 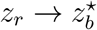, thereby causing the cell state to move on the green and red surface to where the “height” is 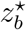. Instead, if gene *r_k_* is perturbed with its value quenched to 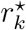, the cell state moves on the red surface to where the “height” is 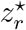 (Fig. 1**c**). Since *z_r_* is the most downstream of all, this response is *not propagated* in the low-dimensional space. However, in the expression space quenching *r_k_* very likely incurs changes in the expression of genes in the blue and green group, but since their low-dimensional representations *z_b_* and *z_g_* remain the same (at 0), the initial cell state traverses a *constant-valued contour* on the blue and green surfaces. Once such a representation is learned, the perturbation-response behavior of gene expression has been reduced to a small set of effective variables that respect the requirements of a specific causal graph.

**Figure 1:**
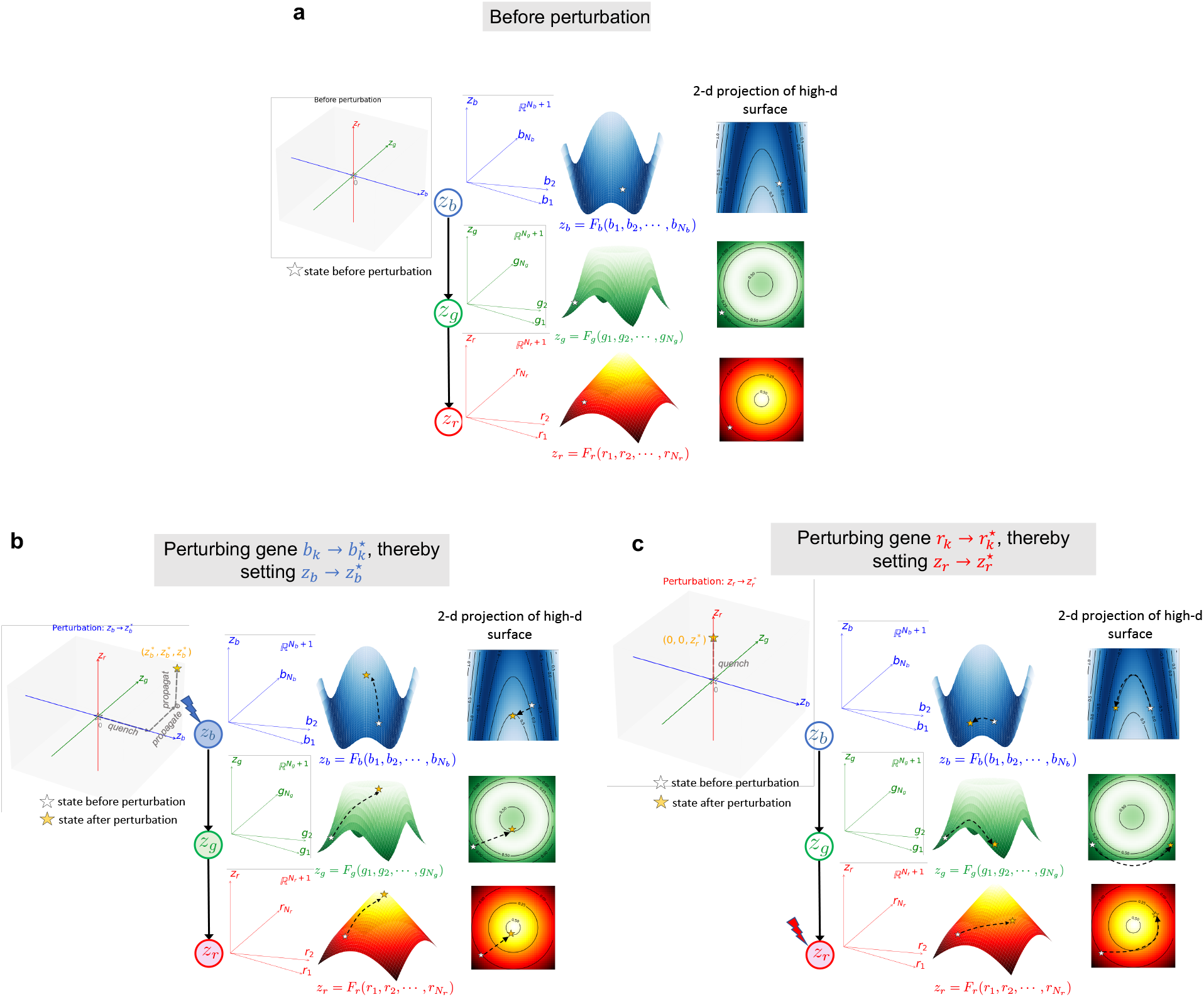
Learning a low-dimensional representation of gene expression that enables accurate prediction of perturbation response. Shown here is an example where the expression of three disjoint (high-dimensional) gene groups {*b*_1_, *b*_2_,…,*b_N_b__*} (blue), {*g*_1_,*g*_2_,…,*g_N_g__*} (green), and {*r*_1_,*r*_2_,…,*r_N_r__*} (red) are mapped to three real numbers, *z_b_*,*z_g_* and *z_r_*, via neural networks *F_b_*, *F_g_* and *F_r_*, respectively. These expressions define three surfaces in 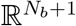 (blue), 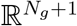 (green), and 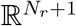 (red) through *z_b_* = *F_b_* (*b*_1_, *b*_2_,…,*b_N_b__*),*z_g_* = *F_g_* (*g*_1_,*g*_2_,…,*g_N_g__*) and *z_r_* = *F_r_* (*r*_1_, *r*_2_,…,*r_N_r__*). Here we stipulate a causal order between *z_b_*, *z_g_* and *z_r_* by imposing a causal graph: *z_b_* causes *z_g_* and *z_g_* causes *z_r_*. **a**: Prior to perturbations, the transcriptional state of a cell defines the locations on these surfaces (indicated by the open star). We assume that neural networks map this state to (*z_b_*, *z_g_*, *z_r_*) = (0, 0, 0) for simplicity. **b**: Perturbing the *k*-th gene in the blue group by setting its expression 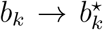 changes the value of *z_b_* to 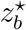 (i.e., the value of *z_b_* is *quenched* to 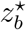). Due to the stipulated causal order, this effect is *propagated* sequentially to *z_g_* then *z_r_*. Assuming no attenuation in the propagated effect, we have that 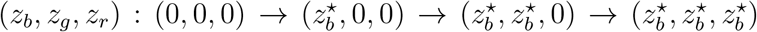. In the expression space, such perturbation brings a cell to a new state defined by new locations on all three surfaces (indicated by the yellow star). **c**: When the *k*-th gene in the red group is perturbed 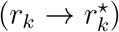, the state of *z_r_* is *quenched* to 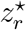. Since *z_r_* is the terminal node in the causal graph, such effect is not propagated, i.e., 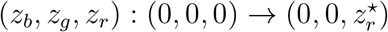. In the expression space, the new cell state differs from prior to perturbation only in terms of its location on the red surface.

### Neural networks that distinguish between different stages of perturbation enables a learning scheme consistent with an imposed causal graph

Training Lowdeepredict begins by randomly partitioning all genes into several large groups of hundreds or thousands of genes, each one of which will correspond to a single node in a causal graph. The implicit assumption in doing this is that as long as each node has a sufficient number and diversity of genes associated with it, the genes in the node will have the potential to capture whatever biological relationships the model fixates on. For example, the dataset we use in this work consists of single-cell RNA-seq expression matrix of roughly 84k cells × 14k genes [5]. A random partitioning into 3 groups (Fig. 2**b**) yields ~ 4.6k genes per group which well-samples a diverse set of biological pathways (e.g. KEGG [26]). We then associate each group *α* with a neural network *F_α_* that maps the expression pattern to a low-dimensional space. To capture the complex dynamics of expression throughout the course of perturbations, *F_α_* takes in a 3-dimensional input for every gene in group *α* which we explain below.

**Figure 2:**
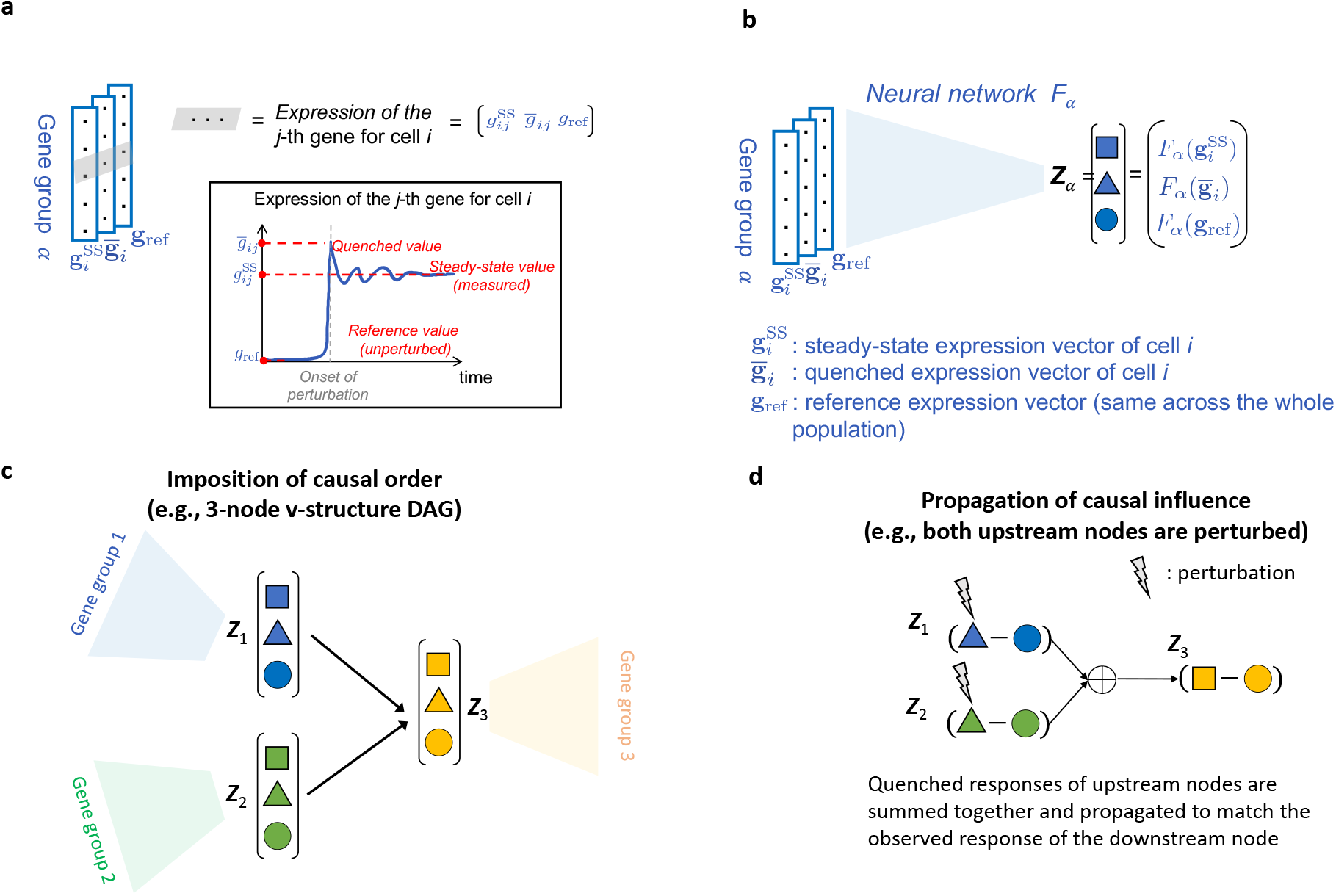
Causal order is imposed in the low-dimensional space learned by neural networks that distinguish between different stages of perturbations. **a**: Our model distinguishes between three expression values: **g**_ref_ is the expression level prior to perturbation, 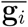 is the value immediately after perturbation (i.e. the value of expression is “quenched” to the level of perturbation), and 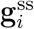 is the steady-state value. For the CRISPRa Perturb-seq data in [5], the steady-state value is processed so that the expression level of perturbed cells is *z*-normalized with respect to the expression of unperturbed cells (which effectively sets the reference expression to zero for all cell indexed by *i*.) The quenched state is to be learned by our model. **b**: Shown here is an example where three disjoint gene groups *α* = 1, 2, 3 are mapped to their corresponding low-dimensional representations *z_α_*’s via neural networks *F_α_*’s. *F_α_* maps the steady-state expression 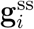, quenched expression 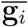, and reference expression **g**_ref_ of genes in group *a* to three real numbers (i.e. low-dimensional embeddings): 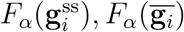, and *F_α_*(**g**_ref_), respectively. We use markers to distinguish these values: 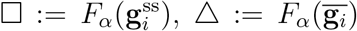 and ○:= *F_α_*(**g**_ref_). **c**: A 3-node v-structure DAG is imposed: both group 1 and group 2 cause group 3. We use colors to distinguish between groups: blue for *α* = 1, green for *α* = 2, and yellow for *α* = 3. Markers are as defined in **b**. **d**: When genes in groups *α* = 1,2 are perturbed, their respective *quenched response*, defined as 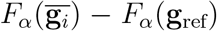 (i.e. blue/green Δ– blue/green ○) are summed together and propagated downstream to match the *observed response* of *z*_3_ (i.e. 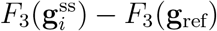, yellow □ – yellow ○). In Fig. S2, we enumerate all possible cases of perturbation to illustrate the mechanism behind the imposition of causal order. Definitions of all quantities are given in Table S2. The mathematical expression of the propagation of causal influence in given in the main text.

Before any perturbation is applied, cells are at the basal state characterized by their *reference expression* which we denote by *g*_ref_. Note that we follow the data processing procedure in [5] so that all cells have the same reference expression, which explains the missing cell and gene index in *g*_ref_. Now consider a perturbation on gene *j* (in group *α*) that “quenches” its value to 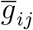, where *i* is the cell index. We call this the *quenched expression* as this is what we would have measured at the onset of perturbations. Following this, the perturbation effect is propagated throughout the unknown genetic network that involves *d* » 1 genes, after which the expression of gene *j* relaxes to a *steady-state* 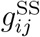 (which we call the *steady-state expression*, see Fig. 2**a**). There are a few things to note before we describe how our proposed neural network utilizes these data to impose causal order. First, in addition to gene expression, every cell *i* in group *α* (with size *d_α_*) is associated with a one-hot encoding vector ***μ**_i_* ∈ {0,1}^*d_α_*^ that indicates which genes in *α* are perturbed in cell *i*. For the dataset in [5], these are the barcodes that cells carry about which gene or pair of genes are perturbed in the CRISPRa experiment (see Fig. S1 for the perturbation coverage). Second, although quenched expression is never measured, it can nonetheless be computed as 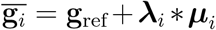, where 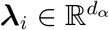 is the *perturbation weight vector* (to be learned by the model) that represents the strength of the applied perturbation and 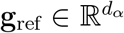 is the reference expression of genes in group *α*. Throughout this work, we assume that there is one single perturbation weight parameter λ across all cells and all perturbed genes to reduce the number of learnable parameters, i.e. 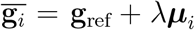. Doing so reduces the number of trainable parameters (*d*, the number of genes) and helps with convergence.

Our proposed neural network *F_α_* maps the steady-state expression 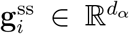, quenched expression 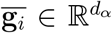, and reference expression 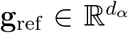 of genes in group *α* to a tuple of three real numbers 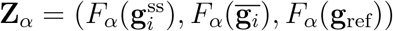 which we use to indicate the state of each group (Fig. 2**b**). We then stipulate causal relationships between **Z**_*α*_’s. In Fig. 2**c**, we have that both **Z**_1_ and **Z**_2_ cause **Z**_3_, namely, they form a v-structure (i.e. “collider”) directed acyclic graph (DAG). The intuition is that perturbations of genes in group 1 (2) should only affect the state of **Z**_1_ (**Z**_2_) and **Z**_3_ but not of **Z**_2_ (**Z**_1_). In addition, perturbations of genes in group 3 should only affect the state of **Z**_3_ since **Z**_3_ is the leaf node of the causal graph (i.e. it has not child). When genes in group 1 and 2 are perturbed, we require the effect to propagate linearly to the affect the state of **Z**_3_ (Fig. 2**d**). To enforce this mathematically, we first define the *quenched response* for node (i.e. gene group) *α* as the difference between the quenched and reference state,

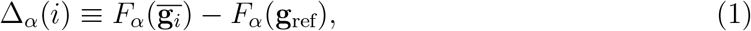

as well as the *observed response* as the difference between the steady-state and reference state

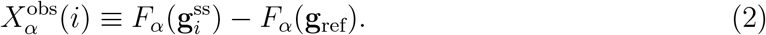

We then add up the quenched response of upstream nodes and propagate it to match the observed response of nodes downstream. Concretely, let **A** be the adjacency matrix of the low-dimensional causal graph, where **A**_*αβ*_ = 1 if and only if node *α* → node *β*. The *propagated predicted response* that node *α* receives is given by

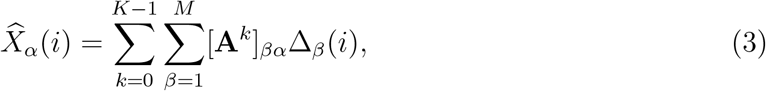

where *K* is the smallest positive integer for which **A**^*K*^ = 0, assuming **A** corresponds to a directed acyclic graph (DAG). Note that **A**^0^ = *I* (the identity matrix.) Neural networks *F_α_*’s are trained such that we can use the predicted response 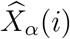 to accurately approximate the observed response 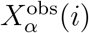 through their mean squared difference (see *Methods: Causal loss function* for details). A simple example is shown in Fig. 2**d** where genes in group 1 and 2 are perturbed. In this case, the quenched responses of these nodes are added up to match the observed response of 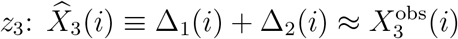. In Fig. S2, we enumerate all other possibilities of perturbation for the causal graph in Fig. 2**c** pictorially. We also list the mathematical expression of causal loss for all these conditions in Table S1. Note that additional loss terms are added to the causal loss to avoid trivial solutions and to improve training performance and interpretability, see *Methods* for details.

Overall, we propose a neural network scheme that takes in gene expression from different stages of a typical CRISPR perturbation event. The neural networks are trained by minimizing a causal loss function that enforces causal order between the low-dimensional representations of genes. This results in a model that maps high-dimensional gene expression to a low-dimensional space where the causal order is consistent with the biological response between genes subjected to perturbations.

### Extensive search in the space of causal graph and gene assignment schemes reveals models with high predictive success

Our modeling approach proceeds by fixing a choice of causal graph by assumption and then training arbitrary functions of gene expression represented by neural networks in order that their outputs obey the rules of that graph. This procedure of course raises the question: which causal graph should be chosen? Though more advanced techniques might be used in future, here we train separate models for a diverse range of small graphs, with genes assigned to each node of the graph by random assortment. The search is restricted to the space of directed acyclic graphs (DAGs) in order that the resulting models will admit a simple causal interpretation. Note that although this greatly reduces the space of graphs to perform search on, the number of DAGs with *m* nodes, *G*(*m*), is still super-exponential in *m*, e.g., *G*(2) = 3, *G*(3) = 25, *G*(4) = 543, *G*(5) = 29, 281, *G*(6) = 3, 781,503 etc. [22]. For this reason, we elect to consider 28 graphs consisting of all possible 2-node and 3-node weakly-connected DAGs (2 + 18 graphs), 2 (out of ~ 500) 4-node weakly-connected DAGs, 3 (out of ~3M) 6-node weakly-connected DAGs, and 3 (out of ~ 7.8 × 10^11^) 8-node DAGs, see Fig. S3 for their topology. For each graph, we partition genes randomly using 3-4 different random seeds, which yields 169 models trained and each optimized to select the combination of hyper-parameters (using Optuna [27])that yields the best validation node-average Pearson correlation between the predicted and observed response.

In Fig.3**a**, we show the node-averaged Pearson correlation for models we trained and optimized as described. For 3-node DAGs, we stratify the results according to graph topology: v-structure, cascade, common parent, and feedforward loop. The best performing model from this analysis is a three-node v-structure graph attaining 0.86 node-average validation Pearson correlation *ρ* between predicted and observed perturbation response of the graph nodes. We use its magnitude |*ρ*| as a metric to gauge model performance, as it quantifies the correlation between the response (in the low-dimensional space) expected by the model and what was measured in the experiments. In Fig. 3**b**, we plot the joint density of the observed response, Equation (2), and the predicted response, Equation (3), for this model and show the Pearson correlation between them for each node and across training, validation, and test data (80-10-10 split). Note that this model achieves 0.84 node-averaged Pearson correlation with test dataset. These results combined to show that our training procedure that aims to enforce causal order through a pre-specified causal graph can produce low-dimensional representation of genes in which the *measured* perturbation response in highly consistent with what the model predicts. Note that as a negative control, we train 400 models on graphs with zero connectivity (i.e., disconnected graphs). We find that models as such are hard to train (with averaged performance |*ρ*| ~ 0.5), see Fig. S19. We speculate that the diminished performance is due to the lack of constraint in the loss landscape from the connectivity of the imposed causal graph.

**Figure 3:**
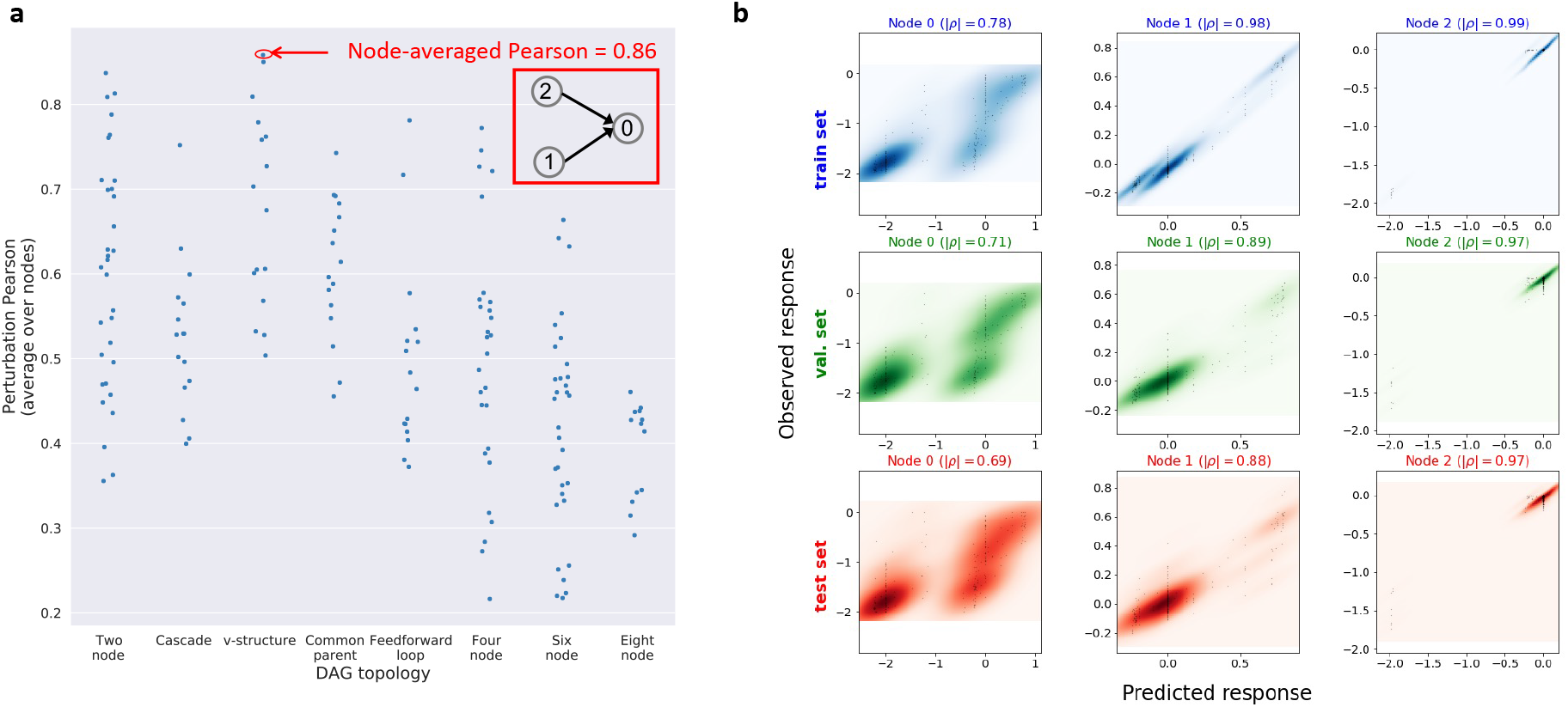
Searches in the space of causal graph and gene assignments selects a causal graph that is highly trainable and with good test performance. We train, in total, 169 models with different DAG topologies, each with 3-4 random gene bucketing/assignment schemes. The search in the DAG space consists of all possible 2-node and 3-node weakly-connected DAGs (2 + 18), 2 (out of 543) 4-node DAGs, 3 (out of 3,781,503) 6-node DAGs, and 3 (out of ~ 7.8 × 10^11^) 8-node DAGs, see Fig. S3 for their topology. All weakly-connected 3-node DAGs fall into one of the followings: Cascade, v-structure, common parent, and feedforward loop. **a**: Shown here are the node-averaged Pearson's correlations between the predicted response and observed response computed using the validation data stratified by graph topology. Members in each topology is given in Fig. S3. Graph structure of the best performing model (indicted by red circle) is shown in the inset. **b**: Observed response is plotted against the predicted response for the best performing model. Color indicates density estimated with Gaussian kernels. Columns are labeled by node number corresponding to the graph in the inset of **a**. Top row shows the result for training data, the middle row shows that for the validation data, and the bottom shows that for the test data. Node-wise Pearson’s correlation magnitude |*ρ*| is indicated for each node and data type (train, validation, or test).

It should be noted, however, that such a model should not be thought of as a uniquely predictable low-dimensionality in the data; it is very reasonable to suppose that there could be more than one way of projecting high-dimensional gene expression into a few quantities for which a certain causal graph relation is obeyed. Indeed, our results showed a number different causal graphs that could be satisfied with relatively high correlation, particularly with different random assignment of genes to nodes. Nonetheless, what the model training discovers in the data is a robust rule for predicting the downstream, low-dimensional effects of CRISPR activation across a range of different perturbations of gene singles and pairs.

### Group perturbation response accurately identifies related gene groups

In the above, we have trained a model to find functions that map high-dimensional gene expression to a small set of node variables that obey a simple, known causal structure. Such a model will only be useful, however, if the low-dimensional predictability it uncovers can be employed to reveal biological relationships that can be understood and acted upon. A large number of genes’ expression levels may contribute in principle to the value of a function *z_α_* = *F_α_*(**g**), but after model training, one might think that only specific combinations of expression changes from a subset of these genes will actually determine the value of node *α*. In Figure S14**b** we show the distribution of gradient magnitudes of the functions *F_α_* with respect to all of their gene arguments. Based on the gradient magnitude, we select the top-ranking genes and annotate them with KEGG pathways based on functional enrichment analysis through DAVID [29]. It turns out, except for nodes where we explicitly assign KEGG pathway genes to (e.g., node 0 in Figure S14**a** that consists entirely of genes from selected KEGG pathways), we do not get meaningful and significantly enriched annotations, see Figure S14**d**.

Since basic, single-gene measures of feature importance do not turn out to have obvious meaning in the model, we must consider a more novel approach. We note that the model is trained to detect predictable patterns in response to perturbation, and also that it does so by computing quantities that integrate information from the expression levels of many genes at once. We therefore now ask which *groups* of genes are predicted by the model to have a particularly strong response in the causally downstream node variable(s) when perturbed. Since the model is aimed at integrating information from the expression levels of many genes simultaneously, it stands to reason that part of the predictability in the data may be good at identifying may require examining collective, multi-gene properties. We stress that our attempt is to interrogate the potentially nonlinear interactions between genes that explain the high predictability in the projected low-dimensional space, rather than predicting the single-cell transcriptional state under multi-gene perturbation which is technical difficult in experiments. Therefore, the notion of “group perturbation response” presented here should be construed as a means by which meaningful interactions are revealed instead of an in silico tool to impute the cellular response to experimentally infeasible multi-gene perturbation [19].

To examine this possibility, we develop the concept of a group perturbation response. To simplify notation, we use [*d_α_*] = {1, 2,…, *d_α_*} to index all the genes in node *α*. Let *S* ⊂ [*d_α_*] be the index set for the genes of intereset. Similar to how we define one-hot encoding vectors, let ***μ**_S_* ∈ {0,1}^*d_α_*^ be the binary vector whose elements are zero except for those indexed by *S*. We define the **group perturbation response** of gene set *S* (whose members are in node *α*) under perturbation of strength λ as:

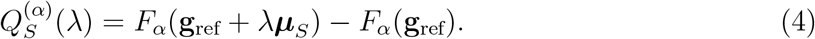

where **g**_ref_ is the reference expression of genes. Note that this definition mimics the finite difference version of directional derivative of *F_α_* in the direction of **v** = ***μ**_S_* evaluated at **g**_ref_ with step λ (without the normalization by λ). The motivation for this definition is that it is possible to ask the trained model for any group of chosen genes in the upstream nodes of the causal graph how strongly their simultaneous perturbation is predicted by the change in the node variable value at the perturbed node, such that this change is expected to be communicated to a corresponding change in the downstream node. In the data set employed here, perturbations were all single- and double-CRISPR activations, but only ~ 300 perturbations were actually performed. The trained model, however, can make mathematical predictions about any arbitrary set of combined perturbations

Due to the non-linearity of the functions of gene expression learned by our model, we note that cooperativity between genes may have the chance to play an important role in contributing to the predictive success in capturing the structure of biological pathways. We therefore hypothesize that the group perturbation response measure may correlate with different kinds of ground truth relating groups of genes to each other in the cell.

To investigate this question, we design a benchmark experiment that consists of a pre-selected sets of pathway genes and random genes (see *Methods* and Table S4 for details). Each gene is assigned to one of the two nodes in a simple causal graph *H*_Lowdeepredict_ (Fig. 4**a**), depending on their degree of upstreamness (defined in Equation (21)) in source of ground-truth for causal order [28] - genes with non-zero upstreamness (labeled *Pathway group* in Fig. 4**a**), meaning they are the causal parents of some other genes, are placed in the upstream node 0 while those with zero upstreamness (labeled *Dowstream (pathway) group*), meaning they are the leaf nodes in the ground-truth gene-gene interaction network, are placed in the downstream node 1. The upstreamness of a gene is determined based on a set of “groundtruth” derived from MetaBase (Clarivate) by filtering the overall transcriptional regulation for “causal” only interactions [28].

**Figure 4:**
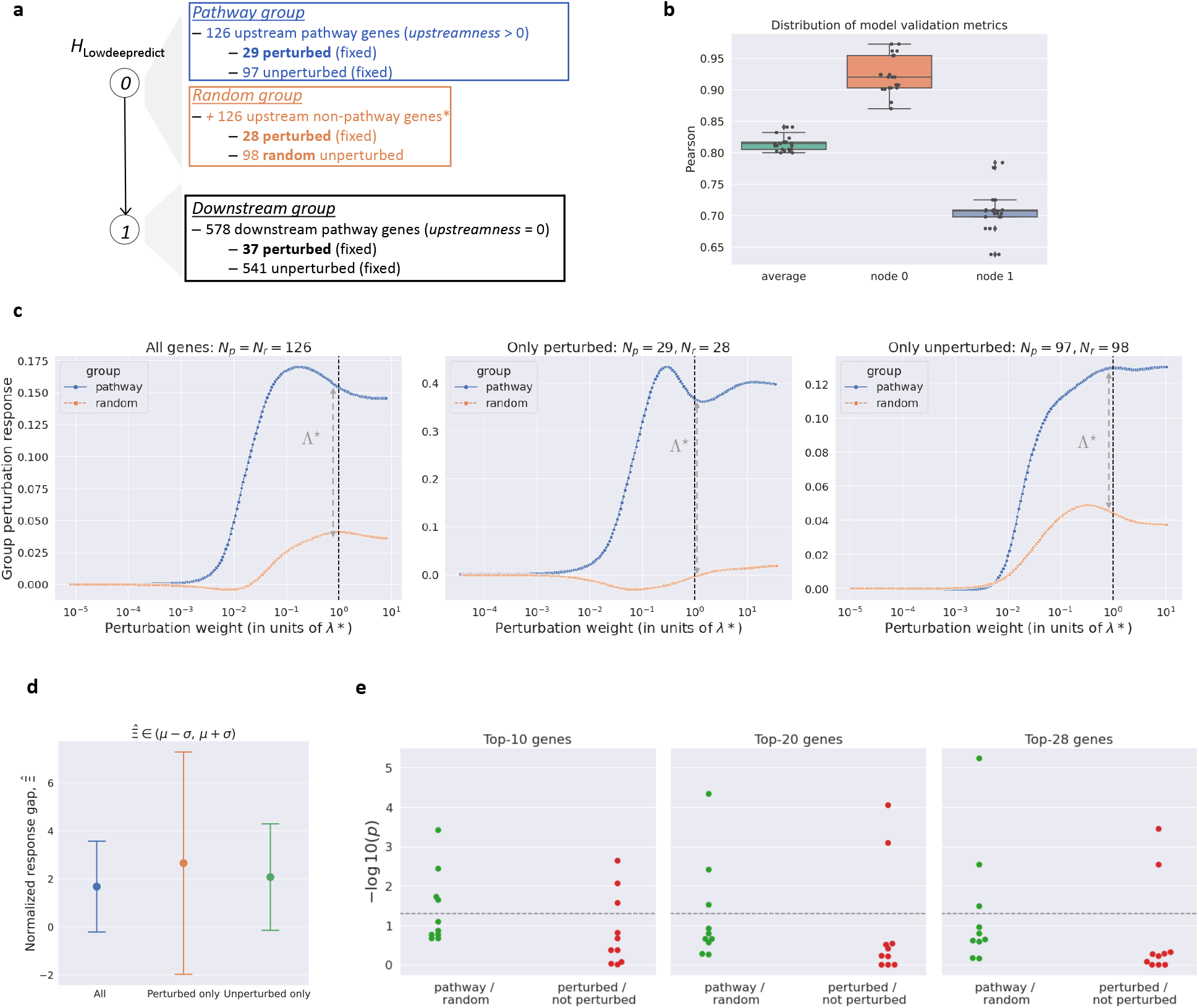
Group perturbation response accurately differentiates between the pathway from the non-pathway genes. **a**: Design of causal benchmark experiment. We select 704 genes from a set of KEGG pathways that are related to cell growth/death, signaling, and general transcriptional pathways (see Table S4). These genes are called *pathway* genes. As described in *Methods*, they are assigned to two groups depending on their upstreamness, defined in Equation (21): those with positive upstreamness are in the upstream node 0 *(Pathway group*) while those with zero upstreamness are assigned to the downstream node 1 *(Downstream (pathway) group*). To maximally utilize the data in [5], we assign all the remaining (i.e. after the pathway gene assignment) 28 genes that were experimentally perturbed to the upstream node. These upstream non-pathway perturbed genes are validated to not cause any genes in the downstream node and any of the upstream pathway genes (blue box) using causal ground-truth [28]. In this benchmark experiment, we also assign to the upstream node 98 randomly sampled genes that are validated to not cause any genes in the downstream node. These 98 genes and the 28 perturbed non-pathway gene constitute the *Random group* (orange box). A breakdown of gene composition for each node is summarized in Table 1. **b**: Model performance for the benchmark ensemble. **c**: Shown here are the response magnitude curves 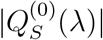 plotted against the total perturbation weight λ from a Lowdeepredict model trained according to **a**. Blue curves refer to *S* = pathway group and orange curves refer to *S* = random group. The left panel uses all genes in each group, the middle panel focuses only on the perturbed genes in each group, and the right panel considers only the unperturbed genes in each group. In all panels, 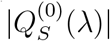 is plotted in units of learned perturbation weight λ* (≈ 1.6511 for this model). Dashed lines correspond to one unit of λ*. The gap between the pathway and random group, Λ*, as defined in Equation(7), is indicated. **d**: Distribution of the normalized response gap. The normalization amounts to rescaling the response gap, Λ*, by the geometric mean of gaps across the 10 experiments in the ensemble. The x-axis indicates the basis of comparison: compare between all genes in each group (blue), only the perturbed gene in each group (orange), or only the unperturbed genes (green) in each group. Note that all the gaps are positive. **e**: We perform the onesided Mann-Whitney U test with the null hypothesis that genes in the pathway group are stochastically equal in their gradient magnitude than those in the random group. A *p*-value < 0.05 favors the alternative hypothesis that pathway genes are stochastically higher in score than genes in the random group. Results from a similar test comparing the perturbed genes against the unperturbed genes are shown as well. In conducting the test, we focus only on the top-*k* (*k* = 10, 20, 28) genes in terms of single perturbation response magnitude. Dashed line indicates *p*-value = 0.05. Note that the *y*-axis is – log_10_(*p*).

**Table 1:**
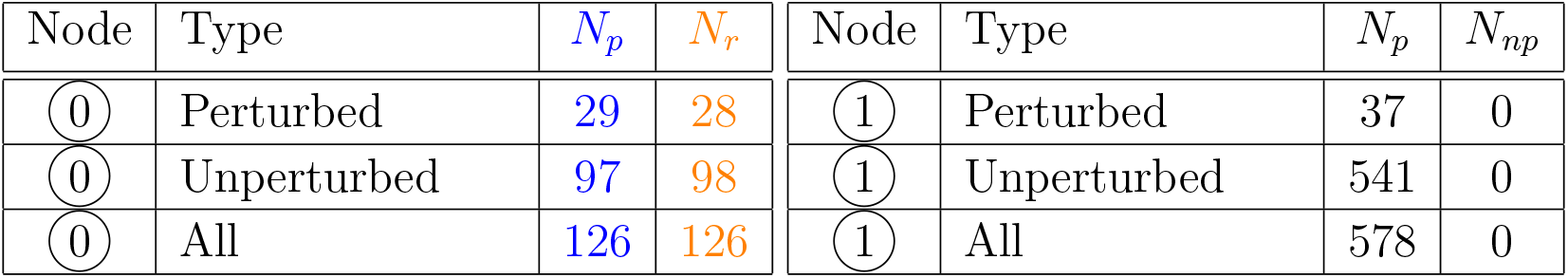
Gene composition of the benchmark experiment described in Fig. 4 and 5. Shown here are the number of genes in the pathway group (*N_p_*), random group (*N_r_*), and the non-pathway group (*N_np_*) for the benchmark experiment. Node refers to the graph in Fig. 4**a**.

In addition to these pathway genes, we randomly sample genes that are not in the selected pathways and assign them to the upstream node (labeled *Random group*). The number of random genes is chosen such that the perturbed and unperturbed populations (in the experiments conducted in [5]) are almost identical across the *pathway group* and *random group* (see Fig. 4**a** and Table 1). Ultimately, we want to assess if, conditioned on the perturbation status or not, genes in the *pathway group* tend to have high group response compared to those in the *random group*.

Since this benchmark experiment requires random sampling, we construct an ensemble of 10 experiments as described above, except that the random seed used for the sampling are different across experiments. For each experiment, we train a Lowdeepredict model with *H*_Lowdeepredict_ as the imposed causal graph and perform hyperparameter optimization [27]. Overall, the ensemble achieves around 0.8 Pearson correlation on average (Fig. 4**b**). For each trained model, we apply in silico perturbation of strength λ to genes in the pathway group (*S_p_* with *N_p_* genes) and the random group (*S_r_* with *N_r_* genes) in the upstream node 0 and measure their respective response magnitude (c.f. Equation (4)):

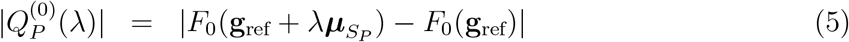

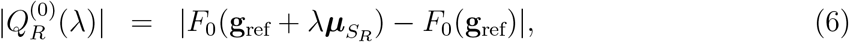

where the subscript *P* and *R* indicate the pathway and random group, respectively. We then vary the perturbation strength λ to trace out the *response curves*. As an example, in Fig. 4**c**, we compare the curve for the pathway group (blue) against that for the random group (orange), conditioned on gene perturbation status (middle and right) as well as unconditionally (i.e. using all genes in each group; left panel). In all cases, we observe that pathway group has higher response than the random group. To facilitate comparison across experiments in the ensemble, we define a scalar metric called the *response gap* to quantify the difference in the group response at the perturbation weight learned by the model, λ*:

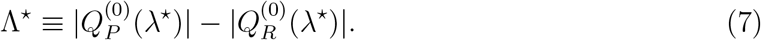

The response gap for each experiment 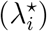 is then normalized by the geometric mean (GM) of all gaps across the ensemble: 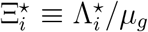, where 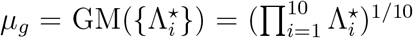. Such normalization ensures that gaps from different experiments (as scale factors) are compared on an equal footing. Note that 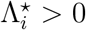 across the 10 experiments, which makes their GM well-defined. This also means that pathway genes consistently have higher response magnitude than the random genes. In Fig. 4**d**, we show the distribution of the normalized gap.

To assess if the pathway genes and random genes can be distinguished under simpler interaction score (as opposed to multi-gene perturbation described above), we compute the average gradient of each gene (as described previously). We then conduct one-sided Mann-Whitney *U* tests to compute a *p*-value under the null hypothesis that both groups are stochastically indistinguishable. This test stipulates an alternative hypothesis that the pathway group is stochastically higher in score than the random group. Since empirically the group distributions have fat tails, we opted to compute the *p*-values with only the top-*k* (*k* = 10, 20, 28, number of perturbed genes in the random group) genes within each group. For cases where the *p*-value is smaller than, e.g. 0.05, we say that the the null hypothesis is rejected and the alternative is favored. The results are summarized in Fig. 4**e** (colored green). We observe that only around 30% of the ensemble have pathway genes stochastically scored higher than random genes with *p* < 0.05 (in terms of their gradient magnitude). We also conduct the same *U* test to distinguish between perturbed genes and unperturbed genes (Fig. 4**e**; colored red). It turns out ~ 20% of the ensemble have the perturbed genes stochastically scored higher than the unperturbed genes with *p* < 0.05. This suggests that single-gene gradient magnitude does not reliably reproduce the distinguishablity between the pathway and random genes, nor does it consistently differentiate between genes with different perturbation state. Note that in the case of multi-gene group response (Fig. 4**c**,**d**), the distinguishability between the pathway and random genes is independent of the perturbation state. To recap, we design an ensemble of benchmark experiments using known causal ground-truth to show that group perturbation response accurately distinguishes between functionally related pathway gene groups from randomly assembled gene groups.

The discovery of particular examples where genes of related function show a highly cooperative response in the low-dimensional space learned by the model suggests a more general principle may be at play, whereby functional relatedness implies such cooperativity. To test this possibility, we now ask if groups of genes selected to maximize the predicted response are generally enriched for genes that are functionally related. Since the search space of groups of such genes is large, we devise a Monte Carlo (MC) approach to identify gene sets whose *group perturbation response*, i.e., the change in the magnitude of the low-dimensional variable under the simultaneous perturbation of *all* genes in the group, is maximized. In *Methods*, we describe the details of our MC algorithm that identifies a gene set *S* where the magnitude of Equation (4) is maximized (see also Algorithm 1).

#### Algorithm 1 Pathway identification with Metropolis algorithm from ground-up

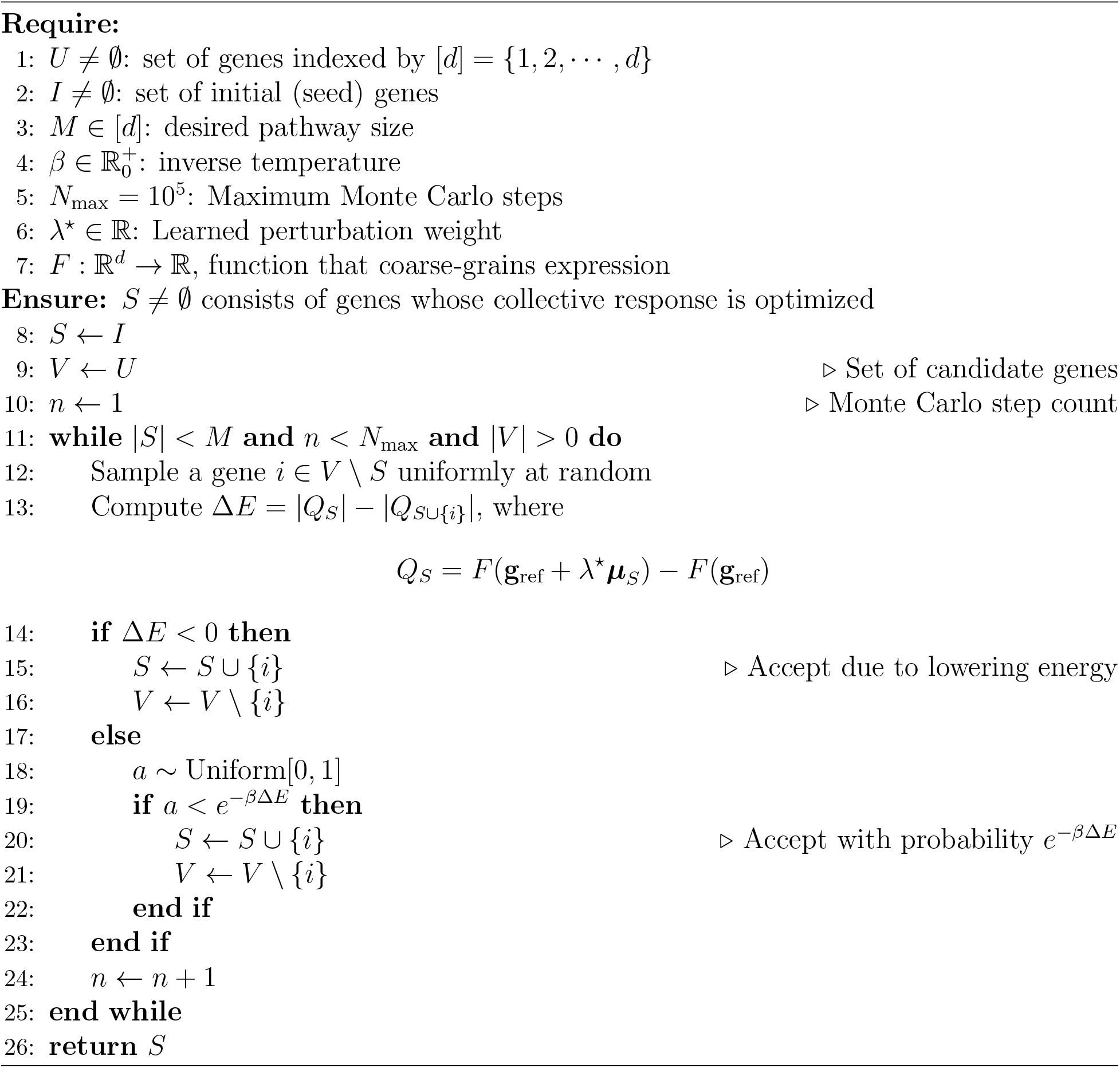

To start, we allow the Metropolis algorithm (a type of MC algorithms) to build a gene set from ground-up, meaning that it starts with a randomly sampled seed gene and builds up the whole set one at a time based on if the acceptance of a gene increases the group response (see Algorithm 1 for details). The algorithm stops if the gene set grows to a targeted size (set to be 100), or it reaches the maximum MC steps allowed (10^5^). The choice for the ground-up algorithm is due to faster convergence for smaller search space of the benchmark model (Fig. 4**a**).

We applied this algorithm to the ensemble study described in Fig. 4**a**. There are 10 experiments in this ensemble representing 10 unique realizations of random genes placement in upstream node 0. For each experiment, we apply Algorithm 1 with 10 different random seeds to identity gene sets with target size 100. Note that the size of identified sets varies, due to the stochasticity in the MC‥ In total, we have 10 (experiments) × 10 (random seeds in MC) = 100 identified gene sets, each of which is analyzed to uncover enriched functional pathways through DAVID [29]. These pathways (enrichment calls) are furthered examined based on the Benjamini adjusted *p*-value at threshold 0.05. A pathway with *p* < 0.05 is called significantly enriched and the number of gene sets calling this pathway as such is counted. This *count*, when normalized by the number of gene sets (= 100, called *maximum count*), is called the *frequency* (of significant enrichment). Note that any given pathway can be called significantly enriched for at most 100 times. In this case, every identified gene set would have called this pathway significantly enriched and the *frequency* of significant enrichment is 1 (or 100%).

In Fig. 5**a**, we show only pathways with *frequency* greater than 10%. We compare the MC results (blue) to those based on randomly assembled gene sets (with size fixed to that of the MC set; colored red) and gene expression clusters (with or without PCA prior to clustering; colored orange and green, respectively, see *Supplementary Methods* for details). We find that MC identifies gene sets with functional annotations consistent with the the biology of the genes we assign to node 0, e.g., the prevalence of significant enrichment calls for transcriptional pathways and signaling pathways (see Table S4). Number of gene sets (i.e. number of enrichment analyses conducted) that produces the result in **a** as well as the distribution of their sizes are given in Fig. 5**b**,**c**. To summarize, through the ensemble of benchmark experiments and gene set enrichment analysis, we show that compared to baseline methods such as expression clustering and random sampling, Monte Carlo algorithm that optimizes group perturbation response identifies genes whose functional annotations are more often aligned with the pathway biology present in the benchmark experiment.

**Figure 5:**
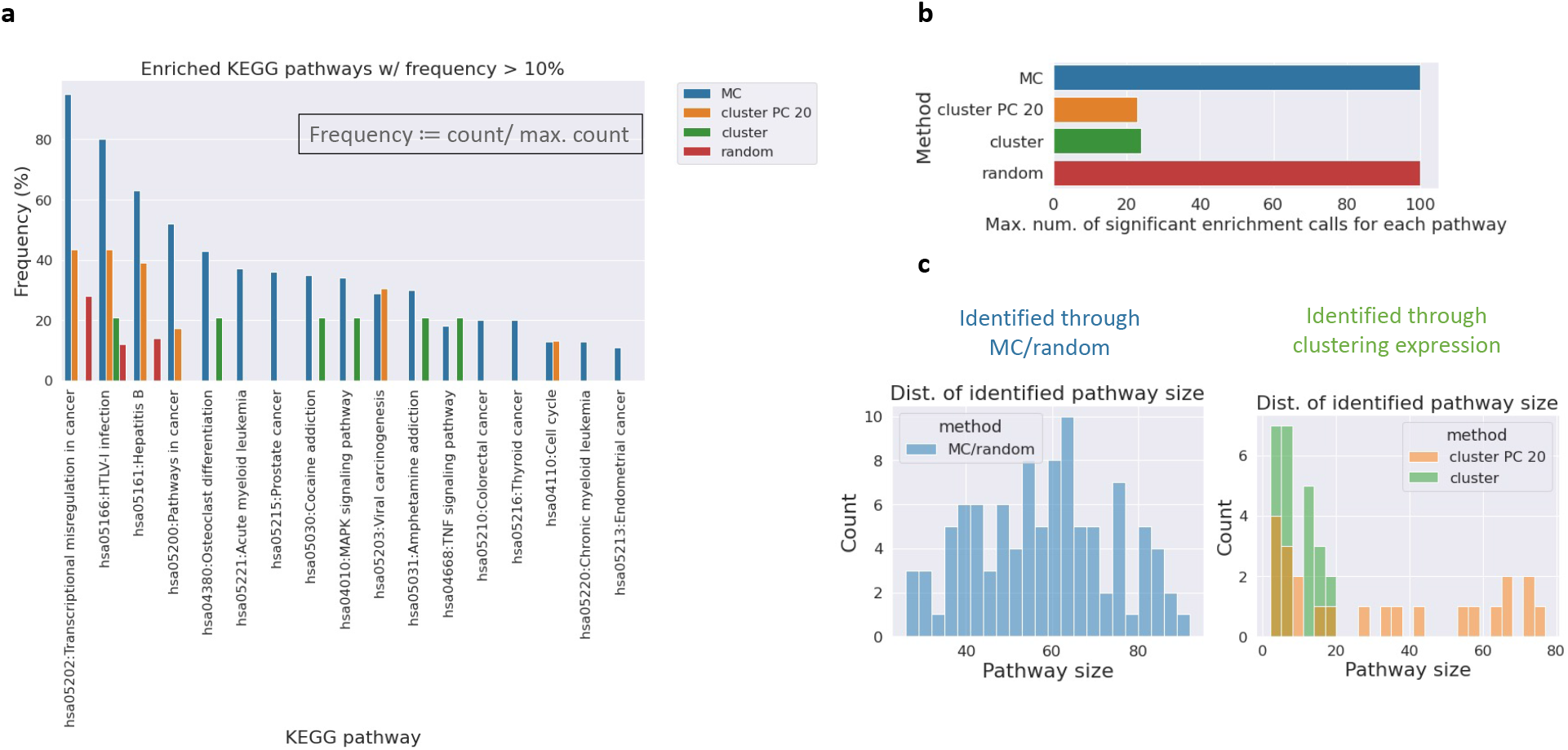
Monte Carlo approach that maximizes group response magnitude outperforms baseline methods in correctly identifying genes that are functionally related. As described in Fig. 4**a** and in the main text, the ensemble study consists of 10 realizations of random gene samplings. **a**: We perform 10 indepdent Monte Carlo runs, each with a unique random seed, to identify genes whose group response magnitude is maximized, see *Methods* and Algorithm 1 for details. Overall, the ensemble study produces 100 gene sets (10 realizations of random gene sampling × 10 MC runs). For each gene set, we then perform functional enrichment analysis through DAVID [29] to report the enriched KEGG pathways. A pathway with Benjamini *p*-value < 0.05 is called significantly enriched and the number of gene sets calling this pathway as such is recorded (i.e. *count*). Note that a KEGG pathway can have at most 100 significant enrichment calls (i.e. *max count*). In this case, the frequency is 1 (or 100 %). Here we show the frequency of (significant) enrichment calls of KEGG pathways based on gene sets identified by MC. These results are compared against baselines such as pure random sampling (i.e. randomly sample gene sets with size fixed to that of the MC results) and expression clusters using unperturbed cells (with and without applying PCA to the normalized expression matrix before clustering). In the legend: *PC 20* (keep only the top-20 principal components), *MC* (Monte Carlo), *random*: randomly sampled gene sets. Note that the missing of bars indicates zero frequency. **b**: Maximum count for various methods are shown. Note that for *cluster* the maximum count is equal to the number of clusters identified (based on expression correlation). **c**: Size distribution of gene sets identified by various methods is shown.

### Monte Carlo algorithm optimizing group perturbation response identifies gene sets with pairwise interaction consistent with the imposed causal order

In the previous section, we show that our Monte Carlo approach maximizing group perturbation response more often identifies genes with functional annotations consitent with the pathway biology present in a small benchmark experiment. Since this benchmark experiment is designed to be causal, meaning genes with non-zero upstreamness are *upstream* of genes with zero upstreamness, see Equation (21), we ask if this optimization scheme could identify genes with the right causal order in a setting where causality is not put in in the node assignment of genes. To do so, we select 6 KEGG pathways [26] related to cell growth and death and intersect the corresponding genes with those available in the CRISPRa dataset [5]. We then place the overlapping genes (326 in total) in the downstream node 0 of the v-structure DAG in Fig. 6**a**. The remaining genes in the CRISPRa dataset are then partitioned randomly and evenly into two groups (6, 846 genes each), which are then placed in the two upstream nodes (node 1 and node 2). Since the downstream biology is fixed, our model should be forced to learn functions, particularly for the upstream nodes, that, upon receiving perturbation, accurately predict the response of genes related to cell growth and death. In Fig. 6**a**, we show the model predictive performance in terms of node-wise Pearson correlation (averaged to *ρ* = 0.84 using validation dataset) after training and hyperparameter optimization as outlined in *Methods* (see also Fig. S5 for detailed performance breakdown and joint density between predicted and observed response.) Note that the predicted performance for the downstream node 0 (*ρ* = 0.77) is slightly worse than that of the upstream nodes (*ρ* = 0.88, 0.86). This is due to that the causal loss for the downstream node comes from two sources – the effect of self-perturbation (i.e. when genes assigned to downstream node is perturbed) and that propagated from the upstream nodes through perturbing genes in these nodes. The upstream nodes, on the other hand, have to deal with only the self-perturbation effect since they do not have causal parents.

**Figure 6:**
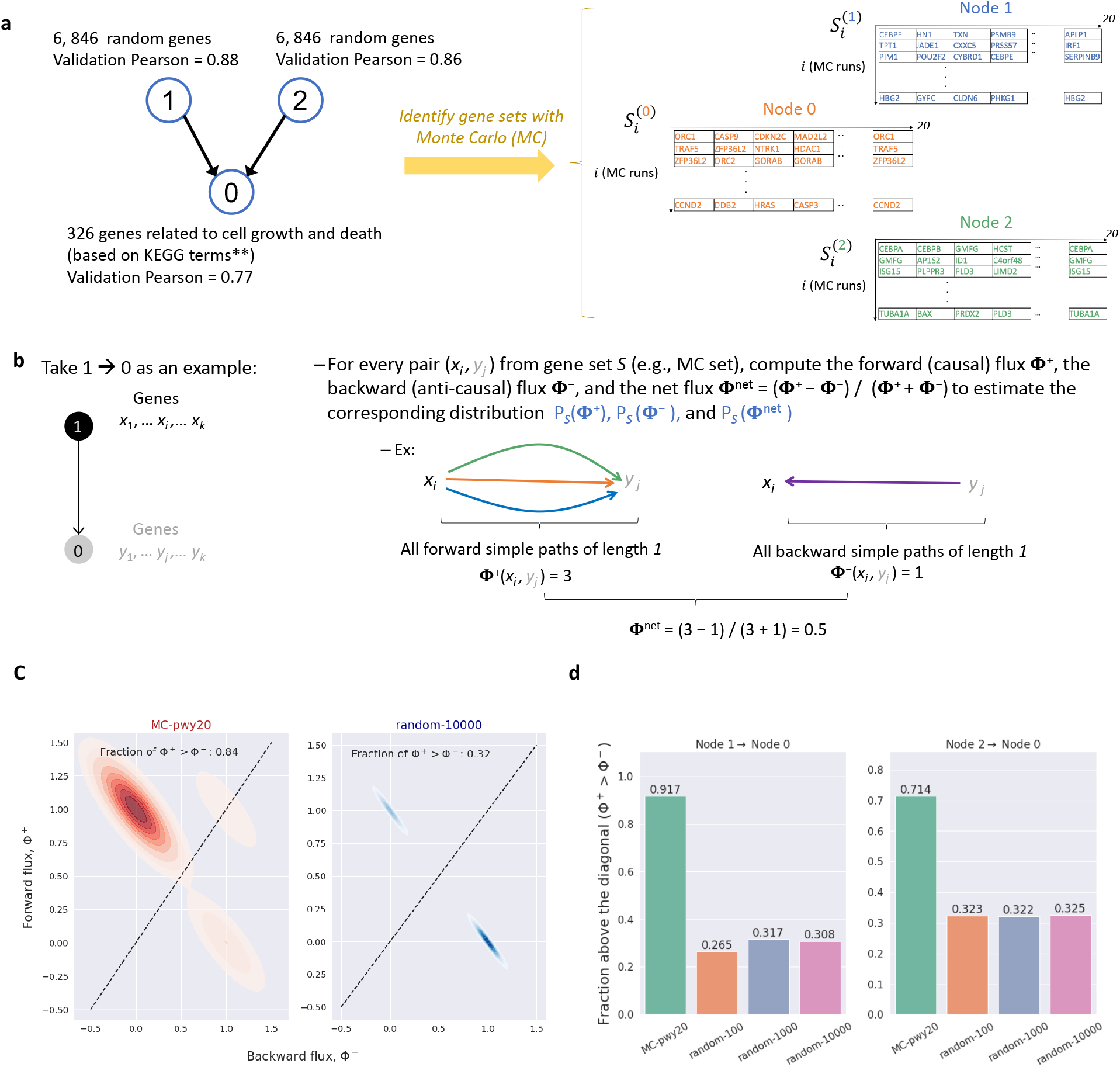
Optimizing group perturbation response with Monte Carlo identifies novel gene sets whose pairwise interactions among constituent genes respect the imposed causal order. **a**: Here we train Lowdeepredict on a 3-node v-structure DAG. Gene placement is such that the downstream node (0) consists purely of genes that are related to cell growth and death, see Table S3 for a list of associated pathways, while the remaining genes are equally and randomly partitioned into the upstream nodes (1 and 2). We use a Monte Carlo (MC) approach (c.f. Algorithm 2) to identify, for each node *α*, a set of genes 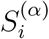, where *i* is the index for independent MC runs, whose group perturbation response magnitude is maximized. Shown on the right is an example of such gene sets. In this figure, we show results for 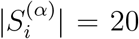 for 20 MC runs (i.e. *i* = 1, 2,…, 20), namely, MC identifies 20 sets of 20 genes for each node. **b**: Illustration on how causal (forward) Φ^+^, anti-causal (backward) Φ^−^, and (normalized) net flux Φ^net^ for any pair of (upstream, downstream) genes are computed. **c**: Joint density of forward and backward flux for both the MC gene set (red) and randomly sampled gene sets (blue). For every edge *e_αβ_* ≡ *α* → *β* in **a**, we first select the top-50 frequently identified genes for node *α,β* from 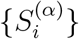 and 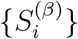, respectively. We then apply the procedure outline in **b** to compute the fluxes with simple path length *d* = 1. Results across all edges *e_αβ_* are aggregated to generate the density plot. The density for the random gene set is produced similarly, except that 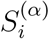 and 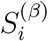 are sampled randomly and uniformly from genes in node *α,β*, respectively (with size 50). **d**: Shown here is the fraction of gene pairs with Φ^+^ > Φ^−^ for both the MC gene set and random gene sets of varying sample sizes (i.e. how many times we sample 50 genes randomly). *MC-pwy20*: MC set generated according to **a**, *random-X*: randomly sample 50 genes X number of times.

#### Algorithm 2 Pathway identification with Metropolis algorithm with replacement

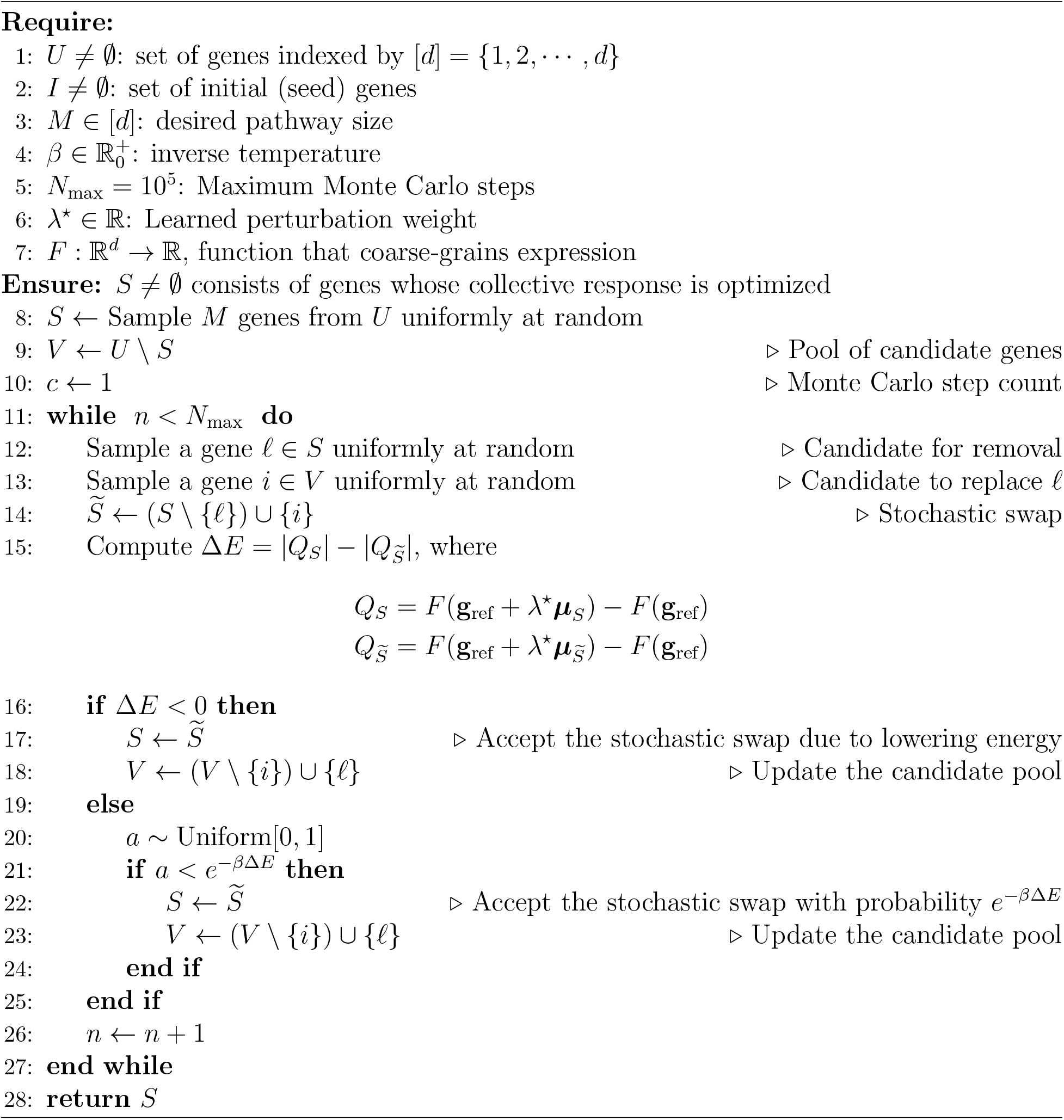

We then apply our MC scheme (Algorithm 2) to identify gene sets with large group perturbation response in each node. In Fig. 6**a**, we show example gene sets 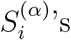, where *α* = 0,1,2 corresponds to nodes in the graph shown on the left and *i* is the index for different independent MC runs. We perform MC with 20 different random seeds (i.e. *i* = 1,2,…, 20) and ask the algorithm to identify, for each run, 20 genes (i.e. 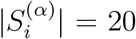) whose collective response in the low-dimensional space is maximized. Note that in this setting, one can have at most min(number of genes in node *α*, 20^2^) unique genes among all the Monte Carlo sets 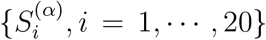. It turns out that gene sets discovered by MC have large overlaps, especially for node 1 and 2 (i.e. those of ~ 6.8k random genes, see Table S5). Analyzing the co-occurrence pattern of genes in these sets reveals that certain subsets tend to be selected by MC more often than others (see Fig. S6, S7, S8, S9.) It’s worth noting choosing genes whose expressions are highly correlated (i.e. co-expression clusters) does not necessitate high response upon their perturbation (see Fig. S13 for details.)

Armed with this discovery, we then ask if the interactions between the most-frequently identified genes contained in these sets are consistent with known literature. To answer that we employ a set of “ground-truth” transcriptional gene-gene interactions as described previously [28] (which we denote as *G*_gt_ and dub causal ground-truth from here on). We further develop an approach to compute, for every directed edge *e_αβ_* ≡ *α* → *β* in the model-imposed causal graph (e.g. Fig.6 **a**), the degree of causal consistency between genes in *α* and those in *β* with respect to known interactions in *G*_gt_. In Fig.6**b**, we illustrate this approach. In short, for every pair of genes (*x_i_,y_j_*), where *x_i_* ∈ *α* and *y_j_* ∈ *β*, we compute the *forward (causal) flux* Φ^+^ (*x_i_, y_j_*) as the number of simple paths (of length 1) from *x_i_* to *y_j_* based on the known causal interactions in *G*_gt_. Similarly, we can also compute the *backward (anti-causal) flux* Φ^−^(*x_i_,y_j_*) as the number of simple paths of length *d* from *y_j_* to *x_i_* with respect to *G*_gt_. Intuitively, Φ^+^(Φ^−^) quantifies the amount of *causal* (*anti-causal*) *flow* for this gene pair. In Fig.6**c**, we show the joint density for Φ^+^ and Φ^−^ for MC discovered gene sets (red) as well as randomly sampled gene sets (blue, sampled 10000 times). These densities come from the aggregating the fluxes for all edges *e_αβ_* in the graph shown in Fig.6**a**. Fraction of pairs (i.e. integrated density) that lie above the 45-degree line, namely, those with Φ^+^ > Φ^−^, are indicated (0.84 for MC gene sets and 0.32 for random gene sets). This result shows that ~ 84% of gene pairs identified by MC have interactions consistent with known ground-truth as opposed to ~ 31% for the random gene sets. In Fig.6**d**, we show such fraction for various sampling sizes (100, 1k, 10k). Consistently, the MC-optimized gene sets have the highest fraction of gene pairs whose causal interactions are compatible with those know in literature. These results combined to show that our MC approach optimizing the magnitude of group perturbation response identifies genes whose pairwise interactions strongly agree with the causal order of the deep-learned low-dimensional DAG model with known causal ground truth. Indeed, when examining gene-gene edges for the top few genes that recur in MC-optimization, we even observe perfect, edge-by-edge agreement between the ground truth and the causal order of our DAG model (see Fig. S10).

### Gene-gene interaction networks constructed with MC gene sets contain high proportion of known causal interactions

In the previous section, we demonstrate that our MC-based approach optimizing group perturbation response identifies genes whose interactions are highly causal. Here we ask if the directional gene-gene interaction networks constructed with these genes are consistent with the known ground-truth. First, we use the top-*k* frequently identified genes from the MC gene set to build an interaction network out of the causal ground-truth *G*_gt_ (i.e. a directional gene-gene interaction network based on known interactions, see *Methods* for details). This network, denoted as *G*_known_, is then examined for its consistency with the model imposed causal graph *H* (Fig. 6**a**). This can be done by, for example, counting the interactions between genes (or edges *i* → *j*, where *i*, *j* are genes) in *G*_known_ not consistent with *H*. An edge *i* → *j*, where *i* ∈ *α* and *j* ∈ *β* (i.e., gene *i/j* is assigned to node *α/β* in *H*) is said to be not consistent with *H* if *α* → *β* is not in *H*. In Fig. 7**b**, we show these graphs for the top-50 genes from the MC set. In this case, *G*_known_ contains 3 edges (colored red) not consistent with *H* (Fig. 7**a**). These are the edges that correspond to node 0 → node 1, which is not present in *H*. After pruning these edges from Gknown one arrives at a graph *G_m_* that fully respects the model mandated causal order. To quantify the degree of causal inconsistency, we use the fraction of edges in *G*_known_ not aligned with *H* (3/20 = 0.15, labeled in Fig. 7**b**). Note that this fraction becomes zero when we used only the top-30 genes to construct *G*_known_ (Fig. 7**c**). In Fig. S10, we vary the gene set size *k* (as in top-*k*) and compute the corresponding fraction of inconsistency. These results are compared against that based on random sampling (1,000 times, each of size *k*). Across the board, the MC gene sets have at most 0.18% inconsistency (or at least 0.82 consistent, when *k* = 60) compared to ~ 0.66 inconsistent (or 0.34 consistent) for the random gene sets. In Fig. S12, we use a variety of causal discovery metrics (e.g. directed/undirected precision and recall, structural hamming distance, graph *p*-values etc., see *Supplementary Methods* for details) to compare the agreement/disagreement between *G*_known_ and *G_m_*. Again, all these metrics show that the directional interactions among the MC causal graph are highly consistent with those known in literature.

**Figure 7:**
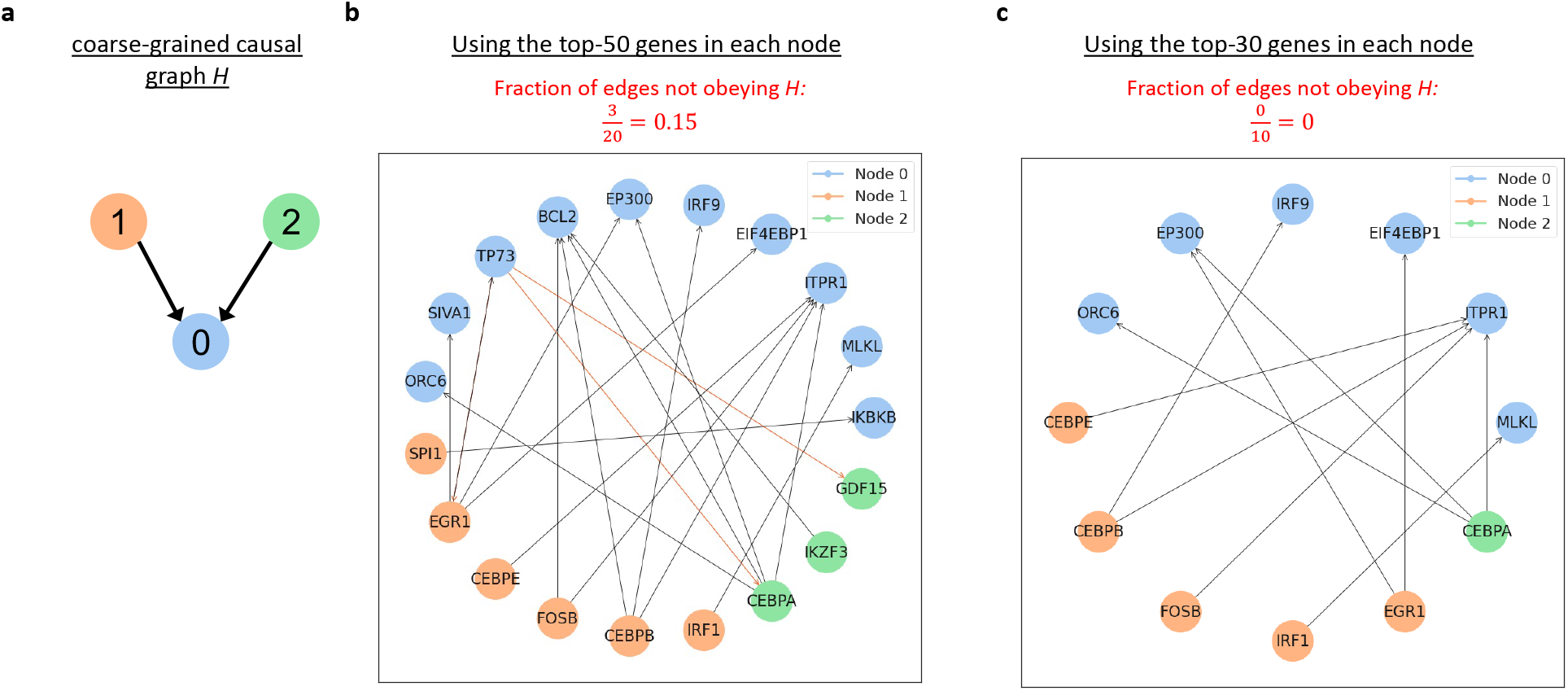
Causal graph constructed from MC genes is highly consistent with known gene-gene directed interaction network. **a**: Imposed causal order for the setup discussed in Fig.6**a**. Here we name it the *coarse-grained causal graph H*. **b**: We use the top-50 frequently identified genes for each node in **a** to construct a subgraph based on the groundtruth causal network *G*_gt_. Details of graph construction is given in *Methods*. The resulting graph contains genes (as nodes) from the top-50 MC set and edges that indicate direct transcriptional interactions found in *G*_gt_. Genes are colored according to their assignment in *H*. In this graph, there are 3 edges not consistent with *H* (colored red): TP73 (node 0) → GDF15 (node 1), TP73 (node 0) → CEBPA (node 1), and TP73 (node 0) → EGR1 (node 1). The fraction of edges not obeying *H* is indicated in the middle. **c**: Results for using top-30 genes. In Fig.S10, we vary top-*k* and compare these results with randomly generated graphs.

### Model imputed synergy is consistent with previously reported fitness measure

In the above, the most basic version of Monte Carlo optimization for generation of strongly cooperative gene groups has been shown to agree significantly with ground truth pertaining both to functional annotation and to causal order. However, the same procedure for nucleating gene-specific groups can also be modified to produce a predicted measure of synergy between two different genes being perturbed simultaneously (Algorithm S1). By comparing the predicted downstream response for scenario in which two genes are perturbed, versus one where only one member of the pair is perturbed, one may construct a quantity, Equation (S.27) in *Supplementary Information*, which measures the degree to which one gene strongly impacts the ability of the other gene's perturbation to cause a large downstream impact. Fortunately, in the same experimental data set with which our deep, low-dimensional causal model is trained [5], there is also an associated cell fitness score, called Genetic Interaction (GI) score in [5], that was measured at the time the perturbation experiment was performed in the same cell population, see Fig. 8**a**, S16. We therefore now devise a comparison of the model-predicted gene pair synergy score (whose derivation only has access to information from experimental gene expression changes) to the corresponding GI measured for the same pair in the same Perturb-seq experiments. To do so, we make use of the v-graph model mentioned above (Fig. 6**a**), which was trained with a selected set of genes involved in cell growth and death in the downstream node. GI scores for the experiment quantify the degree to which the impact of double CRISPRa perturbations deviates from the linear combination of fitness scores measured for the individual single-gene perturbations making up the pair. It therefore is reasonable to ask whether the model’s predicted measure of gene-pair synergy tracks with this experimentally observed non-linearity in impact of the double-perturbation on fitness.

**Figure 8:**
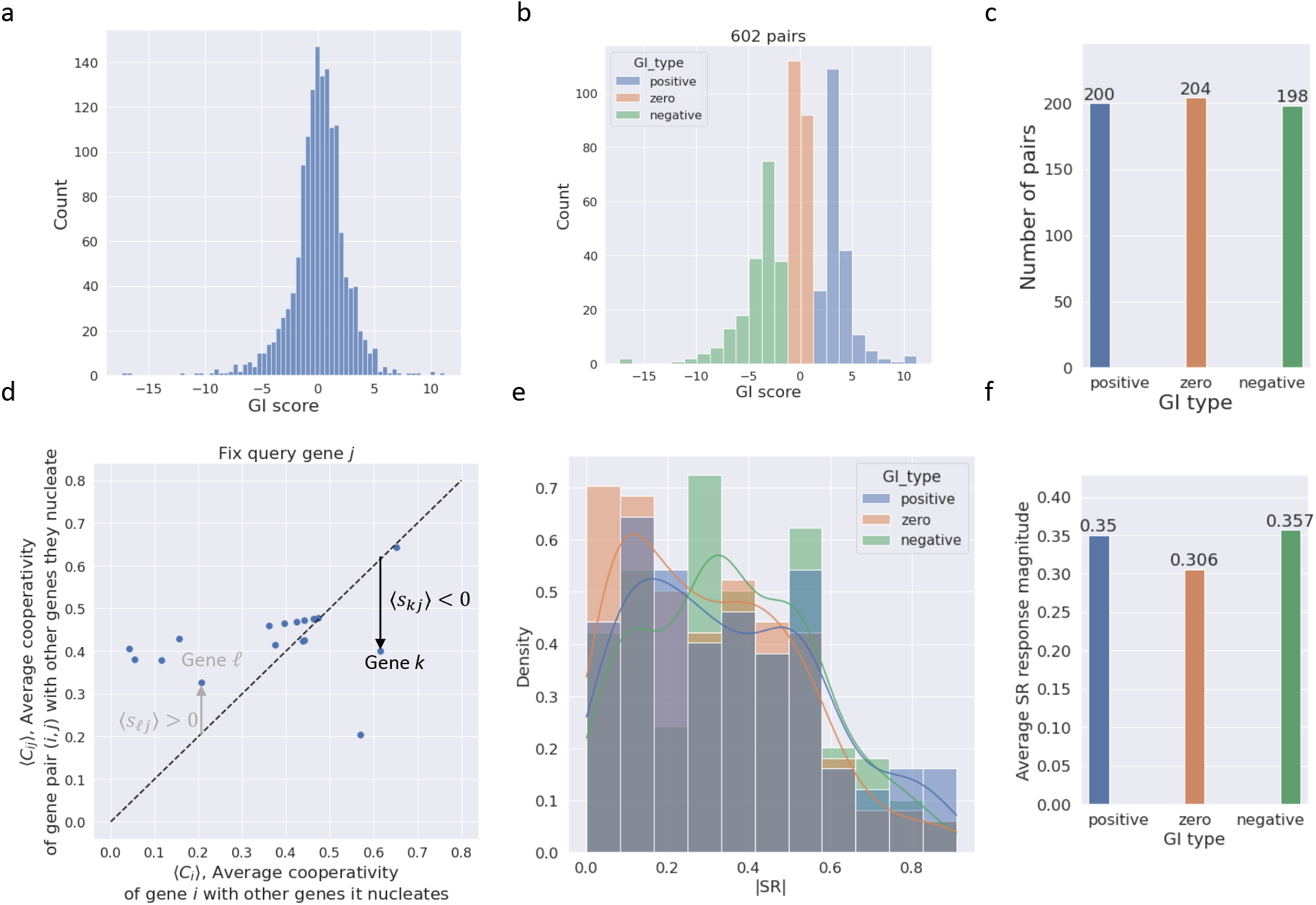
Distribution of GI score and the computation of synergistic response. **a**: Distribution of GI scores. This histogram is compiled from 1483 gene pairs from the GI map shown in Fig. S16 after filtering out pairs whose members belong to different nodes and pairs belong to downstream node 0 in the Lowdeepredict causal graph shown in Fig. 6**a**. **b**: 602 GI pairs are first stratified based on their GI type then sampled from the histogram in **a** for each stratum. Colors indicate GI types (i.e. strata): *positive*: those with GI > 2.25, *negative*: those with GI < −2.0, and *zero*: those with |GI| < 0.3. Number of pairs for each type is shown in **c**. **d**: An example where gene *j* is treated as the query gene in the computation of synergistic response (SR). ‘Sample genes’ *k*, *ℓ* ≠ *j* are highlighted. For gene *ℓ*, since the average cooperativity between the pair (*ℓ*,*j*) and the genes they nucleate is greater than that between *ℓ* and the genes it nucleates, their average synergistic response 〈*s_ℓj_*〉 > 0. For sample gene *k*, we have 〈*s_kj_*〉 < 0. **e**: Distribution of the synergistic response magnitude defined in Equation (S.28). Color indicates the GI type. **f**: Synergistic response magnitude averaged across all pairs of the same GI type (i.e. mean of histograms in **d**).

In order to make this comparison, we divide the number of gene pairs assayed in the experiments in strongly negative, neutral, and strongly positive categories of GI score (Fig. 8**b**,**c**). For these three groups, we then generate histograms of the model-predicted synergy scores. As can be observed in Fig. 8**e**, a statistically significant difference in the high-synergy tails of these distributions is observed, whereby the neutral or non-interacting group of gene pairs has a less predicted synergy. Of course, the large variability in the synergy score predicted from the model in all three groups indicates that these predictions cannot be used to predict GI scores for individual pairs with high accuracy. It should be remembered, however, that the model itself only is being used to produce a prediction about the impact on the node function value for of a large number of genes involved with growth and death, and has not been trained on any labels containing direct experimental measures of cell fitness. Thus, the general trend of being able to detect bias towards higher synergy in the strong GI pairs is an encouraging indication that the causal model has detected some signal of relevant biology *ab initio*.

## Discussion

In this study, we have demonstrated it is possible to extract predictable patterns systematically from single-cell Perturb seq data. By training a neural network to map transcriptomes into a low-dimensional, directed acyclic graph, we demonstrate that the impact of a range of perturbations can be represented predictably in terms of a simple set of causal, linear-response relations. By analyzing the strongly cooperative gene groups identified by the trained model, we have moreover demonstrated agreement with sources of biological ground truth for functional annotation, causal ordering, and synergistic impacts on cell growth and death. In all of these cases, the level of agreement with ground truth was highly statistically significant, but not consonant enough to serve as a replacement for the ground truth. However, what makes this level of agreement striking is that the model was never being trained to discover gene groups of common function or to promote impact on cell death: the gene groups identified through Monte Carlo optimization here have been selected only for the cooperativity and downstream predictability of their combined perturbation. The model training did include an explicit sense of causal order in the architecture, and this is perhaps the reason that the agreement with causal ground truth was the strongest; nonetheless, the sense of causal order discovered by the model had to be learned entirely from data without any causal attribution in the labels.

It has been more common in past approaches to causal discovery in functional genomics data to seek to map relationships between individual genes. Traditional approaches to such a challenge in the full transcriptome case have run aground on the computational complexity of handling so many variables simultaneously, although recent work employing low-rank and sparsity assumptions has demonstrated an efficient means to generate such gene-level graphs [30–32]. Though useful, such highly granular graphs with many interconnected nodes still pose the risk of exhibiting complex and unpredictable behavior at the gene level. We therefore have tried here to articulate a complementary approach that focuses by construction on finding that which is most predictable in the collective behavior of many genes. Our model begins with the stipulation of a low-dimensional causal order and the specification of gene group assignment before training. Guided by the causal loss function, neural networks are optimized towards learning a low-dimensional representation in which gene groups respect the imposed causal order. Due to the high-dimensional nature of gene expression, one should not expect the learned representation to be unique, meaning that there are conceivably many representations of gene expression that achieve the same predictive performance (e.g. the Pearson correlation between the predicted and observed response). Regardless of the nonuniqueness, gene sets identified by our Monte Carlo scheme seem to be highly consistent among models trained with different random seeds that are used to initialize and optimize neural networks but under the same architecture and gene set assignment (see Fig. S15). However, since the MC search space is constrained to genes that are assigned to the same node, a new gene-to-node assignment might reveal hidden associations that are absent in we presented in the main text.

Despite being assembled out of highly diverse and constantly fluctuating types and numbers of molecular components, living things exhibit many predictable behaviors. Traditionally, the characterization of the molecular basis for this predictability has begun by taking a behavior of interest (like cell division) and gradually uncovering groups of genes (such as cyclins) that play key roles in the instantiation of that behavior. High-dimensional biological data present the temptation to run things in the opposite direction by beginning with a panoramic view of many molecular components as they covary under the same conditions and then identifying the groups of genes whose coordinated action is predictable. Such an approach is difficult because it is less anchored in what is already known, but has the potential to discover many new kinds of biological predictability that were previously not identified.

Much still remains to be understood about what the cooperative gene groups identified in this low-dimensional causal model mean, and how knowledge of them might be used to design better future experiments or develop therapeutic strategies. A follow-up set of perturbation experiments that empirically tests whether the multi-gene cooperativity predicted by the model bears out in reality would serve as a natural starting point, as well as a more general attempt to establish with the co-nucleated gene groups can be demonstrated to be involved in some of the same functional tasks. Lastly, a wider range of models can be developed as larger panoramic perturbation-response data sets arrive, and as specific research objectives guide the assignment of genes to nodes in the causal graph in a way that could help to reveal relationships between selected gene groups of interest.

## Methods

### Causal loss function

The idea is to causally propagate perturbation response without using the measured steadystate expression. Such information is only used in the subsequent step to construct the loss function. Algorithmically, this procedure consists of two steps:

For (sample/cell index) *i* = 1, 2,…, *N*

1. Compute the **quenched response**: Compute the quenched response 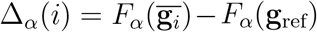, c.f. Eq.(1). Note that this quantity is zero if no genes in node *α* were directly pertutbed in experiment for sample *i*.
2. Compute the **predicted response** by summing up the contributions from the propagated the quenched response:

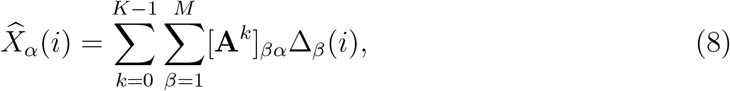

where *K* is the smallest positive integer for which **A**^*K*^ = 0, assuming DAGness. Note that **A**^0^ = *I*, the identity matrix. In matrix notation, this translates into

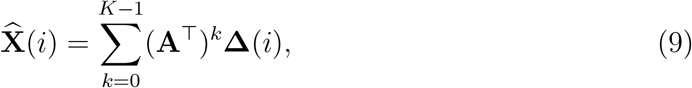

where both 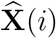 and **Δ**(*i*) are in 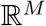 (*M* is the number of nodes in the imposed DAG.)
3. Compute the loss for this sample, defined as the square difference between the **observed response** (c.f. Equation (2))

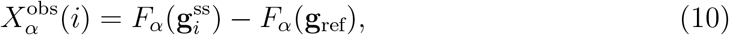

and the predicted response Eq.(3)(9):

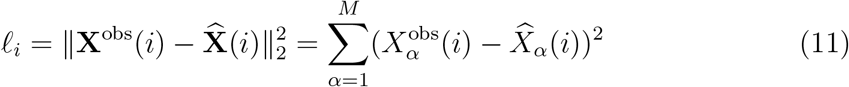

The overall causal graph loss is computed by summing over all (mini-batch) samples

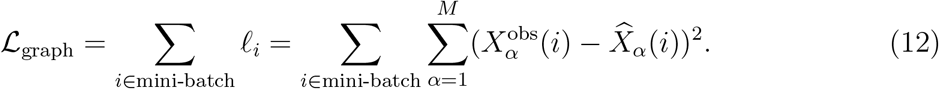

Empirically, we observed that samples receiving no perturbation (neither directly from experiment nor indirectly through parental influence) are given less emphasis throughout training. To mitigate that, we introduce a hyper-parameter *∊* > 0 that places a positive weight when evaluating the causal graph loss on such samples. Concretely, for sample *i*, let **NI**(*i*) ⊂ {1, 2,…,*M*} be the set of nodes not receiving any kind of input perturbation (direct or indirect). The modified causal graph loss becomes:

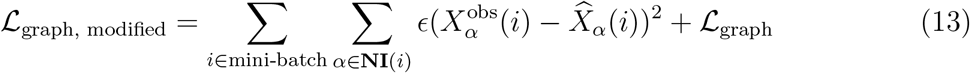

### Variance loss and perturbation strength loss

Based on the causal graph loss, one can easily identify trivial solutions where *F_α_*’s are constant functions. In this case, the causal graph loss ***ℒ***_graph_ achieves the global minimum (zero!) To avoid trivial solutions as such, we impose the following variance loss:

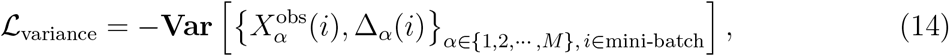

which is the negative variance computed based on the collection of the observed response and quenched response across all the DAG nodes and mini-batch samples, as well as the perturbation strength loss:

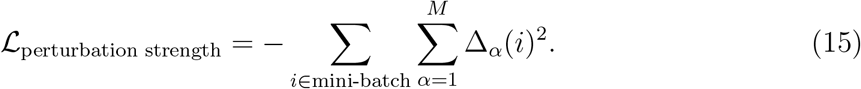

Intuitively, Eq.(14) encourages *F_α_* to be non-constant while Eq.(15) favors distinguishability between the quenched state and reference state (which also implies non-constancy of *F_α_*.)

### Regularization

L2 regularization is imposed:

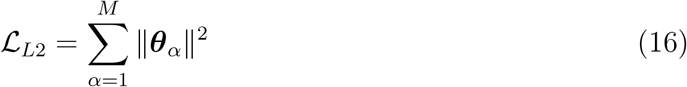

along with dropout.

### Putting all loss terms together

The loss function (to be minimized) is given by, c.f. Eq.(13)(14)(15)(16)

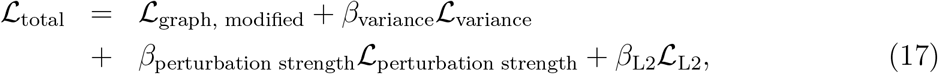

where *β*_variance_, *β*_perturbation strength_ and *β*_L2_ are positive real-valued regularization constants (treated as hyper-parameters).

### Projecting gene expression down to a group subspace

In Table S2, we defined 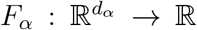 as the neural network that maps the expression of genes in group *α* to a 1-dimensional real number, see Fig. S18 for an illustration. The reduction of dimensionality from *d* to *d_α_* is done by projecting the expression data onto the gene group subspace.

Let **V**_*α*_ ∈ {0,1}^*d*^×^*d_α_*^ be a matrix that projects **g**_*i*_ to a subspace spanned by genes in group *α* through

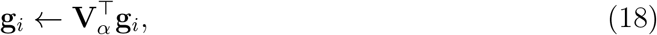

where **g**_*i*_ is a *d* × 1 expression vector for sample *i*. To construct such matrix, first notice that randomly partitioning *d* genes into *M* blocks of size *d*_1_, *d*_2_,…, *d_M_*, respectively, is the same as randomly permuting the index vector **u** through

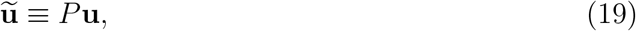

where

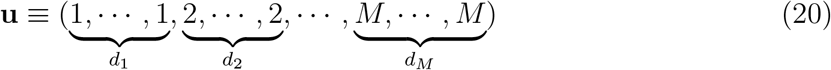

and *P* denotes random permutation. Armed with this observation, we define the projection matrix as **V**_*α*_[*k*, *j*] = 1 only when *j*, *k* are such that the *j*-th element of *M_α_* is *k*, where *M_α_* is the solution set defined as *M_α_* ≡ arg{**ũ** = *α*}.

### Monte Carlo approach to identify gene sets with optimized group response

Monte Carlo (MC) methods are a class of computational algorithms that involves doing some kind of random walk in the space of “configurations” (e.g., of spins or of some microscopic states) [33]. They are particularly useful for problems where analytic solutions are notoriously hard to find. A canonical example in statistical physics is concerned with finding the spin configurations that yield the lowest energy for Ising model. In spatial dimension *d* ≥ 3, such problem is known to be analytically intractable, and one standard approach is to resort to computational methods like Monte Carlo. Here we rephrase the problem of finding a set of genes *S* that maximizes the group response magnitude 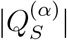 as finding the spin configuration that minimizes energy. In particular, we implement the Metropolis algorithm which is a special case of a more general Metropolis–Hastings algorithm by assuming symmetric random walk proposal density and Boltzmann distribution for the marginal density. For a detailed discussion, we refer the readers to [34].

We implement two versions of Metropolis algorithm: with replacement (Algorithm 2) and from ground-up (Algorithm 1). For the former (used in Fig. 6, 7), one starts by assuming a set of random genes (of the target size) and incrementally replaces genes in this set (one at a time) with candidates that, upon their replacement, increases the magnitude of group perturbation response. The gene in the set to be replaced, called candidate for removal, is chosen at random. The gene to replace the candidate for revmoal is drawn at random and accepted if its replacement increases the response magnitude (c.f. lowering the energy). However, in cases where the solution landscape is glassy (i.e. rugged and full of metastable local minima), one would allow for the acceptance of a gene swap (with probability *e*^−*β*Δ*E*^, where *β* is the inverse temperature and Δ*E* is the energy difference, see Algorithm 2 for its definition) even if it does not lower the energy (or increase the response magnitude). In this case, this algorithm is dubbed *finite temperature* Metropolis. In the zero temperature limit (i.e. *β* → ∞), a gene swap is accepted only if it lowers the energy (or increases the magnitude).

The ground-up version (Algorithm 1, used in Fig.5) follows the same logic– the only difference is that instead of starting with a gene set of the target size, one attempts to build a gene set starting from a smaller seed set and incrementally include new genes (one at a time) into the set if such inclusion increases the group perturbation response. At finite temperature, inclusion is accepted (with probability *e*^−*β*Δ*E*^) even if not increasing the response.

### Causal benchmarking scheme for Fig. 4 and 5

Here we describe the design of the benchmark experiment used in Fig. 4 and 5. The design consists of the following steps:

#### Step 1 Constructing the pathway causal graph (*G*_pathway_)

First, we collect genes from KEGG pathways that are related to cell cycle/growth, apoptosis, signaling and generic transcription (see Table S4 for details.) We then filter them to keep only those in the CRISPRa dataset [5]. These genes, called *pathway genes* in Fig. 4 and 5, are then taken as nodes to construct a directed gene-gene interaction network using the ground-truth causal graph [28]. This graph, denoted as *G*_pathway_, contains 704 nodes and 3,068 edges. Note that *G*_pathway_ covers 66 (out of a total of 94) perturbed genes in the CRISPRa dataset.

#### Step 2 Computing upstreamness

To quantify how upstream a given gene/node is, we define the upstreamness *u*(*s*) for a node *s* as the average of the longest simple path length from *s* to every other nodes in the graph. Formally, let 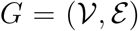 be the directed graph of interest with node set 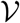 and edge set 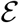. Recall that a *simple path* is a path with no repeated nodes. For every pair of nodes (*s*,*t*), where 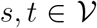, denote the set of *longest simple paths* from *s* to *t* by 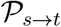. Let *ℓ_s→t_* be the length of such paths. Note that 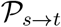 could be 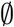. For each (source) node 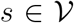, we define 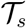 as the set of (target) nodes 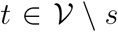 such that 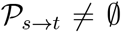 (i.e., descendants of *s*). The **upstreamness** of *s* is defined as:

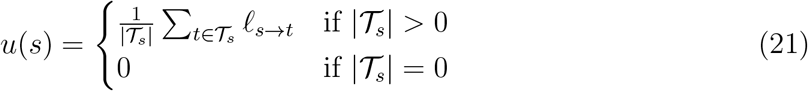

#### Step 3 Constructing causal graph for Lowdeepredict (*H*_Lowdeepredict_)

Based on Equation (21), we can break the nodes in *G*_pathway_ into two groups: 126 with nonzero upstreamness (29 of which were perturbed) and 578 with zero upstreamness (28 of which were perturbed). Genes in the first group are then placed in the upstream node 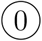 while those in the second group are placed in the downstream node ➀ of Lowdeepredict causal graph *H*_Lowdeepredict_ depicted in Fig. 4**a**.

#### Step 4 Placing the non-pathway perturbed genes in (*H*_Lowdeepredict_)

As we previously mentioned, at this stage there are still 94 − 66 = 28 perturbed genes in the CRISPRa dataset [5] that are not in *G*_pathway_ (called non-pathway perturbed genes.) We eventually assign these genes to 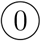 of *H*_Lowdeepredict_ after validating that none of them are

- causal ancestors of pathway genes in 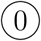 of *H*_Lowdeepredict_
- causal ancestors of pathway genes in ➀ of *H*_Lowdeepredict_,

using the causal graph constructed with *all* entries in the causal ground-truth [28] (which gave rise to a graph *G*_all_ of 9,945 nodes and 39,553 edges). This is to ensure that the causal order *among the pathway group* uncovered by our model comes entirely from this group (to the best knowledge of the causal ground-truth in MetaBase) rather than from these additional genes. Note that the presence of the additional genes is necessary to make the full use of the entire CRISPRa dataset. Note also that *G*_pathway_ is just a subgraph of *G*_all_.

#### Step 5 Randomly sample genes to place in the upstream node of *H*_Lowdeepredict_

At the end of step 4, we placed 28 non-pathway perturbed genes in 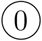. To ensure the equality of group size between the pathway and non-pathway, we randomly sample 126 − 28 = 98 genes from those remaining in the CRISPRa dataset to place in 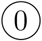 which we call non-pathway random (unperturbed) genes. Gene composition of this benchmark experiment is summarized in Table 1.

## Data availability

Upon publication, our code will be made available at https://github.com/GSK-AI/lowdeepredict. Data used for training is accessible at NCBI Gene Expression Omnibus (GSE133344).

## Supplementary Material

### Supplementary Methods

#### Co-occurrence frequency of genes

Here we define the co-occurrence matrix 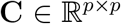 of genes discussed in the main text (see also FIG.S7, S8, S9). At the end of *N_S_* independent Monte Carlo runs, each uncovering, say, *q* genes, we record only the unique genes. Let *p* ≤ *N_S_* × *q* be the number of unique genes. We then construct a matrix 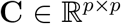 with index *i,j* running over genes in the unique set. The off-diagonal element *C_ij_* is defined as the fraction of gene sets containing both gene *i* and gene *j* while the diagonal *C_ii_* is defined as the fraction of gene sets containing gene *i*. *C_ij_* = 1 means that both gene *i* and *j* appear together in all *N_S_* sets.

#### Constructing the causal graphs in Fig. 7

Here we describe how we construct causal graphs from the top-*k* frequently identified genes from the MC set. Following the notation in Fig. 7**a**, for every edge *e_αβ_* in the coarse-grained graph *H* (i.e. the causal order in the low-dimensional space imposed by the model), we build a subgraph out of *G*_gt_ (i.e. causal graph of directional gene-gene interactions based on MetaBase) by keeping only genes and interactions pertaining to the top-*k* genes in node 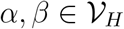, where 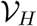 is the set of nodes in *H*. Call this *edge subgraph G_α→β_*. Next we prune the edges of intra-node interaction in *G_α→β_*, namely, removing *i* → *j* whenever *i,j* ∈ *➀* (both in node *α*) or *i,j* ∈ *β* (both in node *β*). The removal of edges as such is necessary since the imposed causal graph *H* only considers inter-node interactions. This procedure is repeated for every edge *e_αβ_* in *H*, and the resulting edge subgraphs are composed to form *G*_known_ (shown in Fig. 7**b,c**). In Fig. 7**b,c**, we color edges *i* → *j* in *G*_known_ that are not consistent with *H* red, meaning those for which 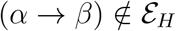 (edges of *H*), for *i* ∈ *α* and *j* ∈ *β*.

#### *p*-value for the causal subgraph constructed by Monte Carlo

Let *S_α_* be the set of unique genes in node *α* identified by Monte Carlo (MC). We first construct a causal subgraph 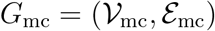, where 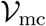 is the set of genes that are in both the MC set, ∪*_α_S_α_*, and the causal ground-truth [28] while 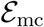 is the set of directed edges concerning 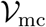 that are found in the causal ground-truth [28], see Table S5 for a summary. We then compute the size of the largest connected component of *G*_mc_. Let *s*_mc_ be its size. To estimate the distribution of the size of the largest connected component, we sample 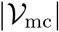 genes from all genes in the dataset, build the corresponding causal subgraph [28], and compute the size of its largest connected component *ŝ*. We repeat this procedure 10^6^ times and record *ŝ* to construct a histogram as a proxy to the distribution *P*(*ŝ*). The *p*-value for *G*_mc_ is defined as the survival function *P*(*ŝ* ≥ *s_mc_*), namely, the probability of observing a graph with connectivity at least as large as *s*_mc_. The results are given in FIG. S11.

#### Clustering the unperturbed expression

We utilize the control cells (i.e. unperturbed cells) from the same CRISPRa dataset [5]. The raw expression matrix of the control population was filtered to keep only cells with at least 200 genes expressed and only genes that are expressed in at least 3 cells. We then rescaled this matrix so that each cell has a total count equal to the median of total counts for cells before normalization. Finally, we “logarithmize” it by adding pseudo-count 1 then taking the natural logarithm. This yields a matrix of 7644 cells × 17538 genes. For a given subset of genes, their expression profiles were clustered using the HDBSCAN python package [35] with the following parameters: correlation distance as the distance metric between two expression profiles, minimum cluster size: 4, minimum samples to call a cluster: 1, cluster selection method: Excess of Mass (EOM).

#### Comparing the correlatedness of expression

Let 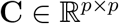 be the correlation matrix (or variance-covariance matrix) of variables *X*_1_,…, *X_p_* with elements

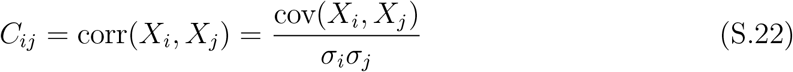

Note that the correlation matrix is related to the covariance matrix Σ by

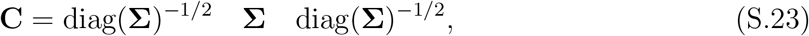

where

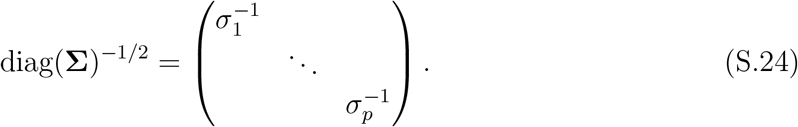

Equivalently, the correlation matrix can be seen as the covariance matrix of standardized variables *X_i_*/*σ_i_*. To summarize the correlatedness of *X*_1_,…, *X_p_*, we defined the **collective correlated coefficient** *η* as

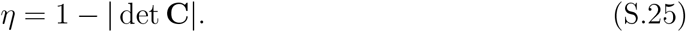

Note that for any correlation matrix **C**_d×d_, 0 ≤ det **C** ≤ 1, and that all its eigenvalues lie in [0,*d*]. Geometrically, the determinant of **C** is the “volume” of the space occupied by the swarm of data points in the space of *X_i_*/*σ_i_*. When these variables are uncorrelated, this space is a hyper-sphere with a volume of 1. When they are correlated, the space occupied becomes a hyper-ellipsoid with volume less than 1. In defining *η*, we compute the difference between 1 and this volume so that higher correlation implies larger *η*. In general, one can take *p*-th root of *η* to compare between correlation matrices of different dimension, but here we simply use *η* since all matrices we're comparing are of the same dimension.

In what situation would *η* attain value 1? One can understand this in terms of PCA. Note that there are two ways to perform PCA: doing eigen-decomposition on the covariance matrix **Σ** or on the correlation matrix **C** (for an extensive discussion, see [36, 37].) Here we stick to the latter. First note that the eigenvalue of the correlation matrix λ_*i*_ is related to the singular value of the data design matrix, *s_i_* via 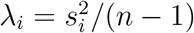, where *n* is the sample size. Therefore, choosing the importance of principal components using the singular value *s_i_* is the same as using the eigenvalues of the correlation matrix λ_*i*_. Since the determinant of a matrix can be expressed as a product of its eigenvalues: 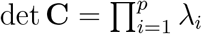, one finds *η* = 1 whenever there’s a zero eigenvalue *λ_i_*. The physical intuition is that perfect collective correlatedness is achieved whenever there’s at least one redundant feature (i.e. one that can be expressed as a linear combination of other features).

#### Definitions of causal graph identification metrics used in Fig. S12

In the following, we use *G_p_* and *G_r_* to indicate the predicted and reference graphs, respectively. Given the adjacency matrix of a directed graph, say, **A**_*d*_, its undirected counterpart is computed by removing its orientations through 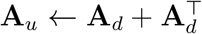 then setting [**A**_*u*_]_*αβ*_ = 1, for [**A**_*u*_]_*αβ*_ > 1.

- **SHD**: Structural Hamming Distance. It counts the number of differences between the adjacency matrix of *G_p_* and of *G_r_*; additional edges, missing edges, and misdirected edges are added up.
- **Binary neighborhood**: The binary neighborhood of two nodes *α,β* in a graph *G* indicates the number of paths of length 2 between *α* and *β* in *G*.
- **Undirected** *p***-value**: This concerns the problem of guessing where edges should be, regardless of orientation. It is given a *p*-value by Fisher’s exact test over the edge-noedge confusion matrix, where the preserved marginals mean that the comparison is against a random selection of the same number of edges.
- **Directed** *p***-value**: This concerns the problem of correctly guessing which orientation edges are, having been provided the informatoin that they are indeed edges. It is given a *p*-value by inspection of the CDF of the relevant binomial distribution, with success corresponding to correctly-oriented edges as a subset of the correct undirected edges chosen previously

#### A Monte Carlo scheme to identify genes that are highly cooperative with user-provided genes of interest

In the main text, we discussed a Monte Carlo scheme to identify genes (collected in a set *S*) whose group response in the low-dimensional space learned by Lowdeepredict, namely,

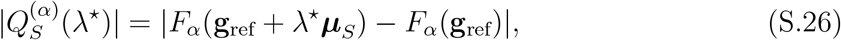

is maximized. Here we assume that genes in *S* are assigned to node *α* in the Lowdeepredict causal graph, e.g., Fig. 6**d**. We also use λ* to denote the perturbation weight learned by the model. Intuitively, λ* is the “amount of perturbation” that is applied to the genes in the dataset that best explains the observed changes in gene expression under the constraint of low-dimensional causal graph. In Fig. 5, we showed that genes in *S* tend to be functionally related. Here we explore another use case for identification scheme of this sort.

One interesting application lies in finding genes that are associated with particular genes of interest where the association is defined according to the phenotypic effect (e.g. cell death) of their collective intervention (e.g. knockout or overexpression). Since the notion of association concerns the interaction between genes, here we extend the original search algorithm in the main text to allow for the identification of genes that (i) highly cooperates with genes of interest and (ii) produce a strong response in affecting genes related to a given downstream biology upon their perturbation, see Algorithm S1 for details. This new scheme starts with a user specified set of genes *I* and stochastically “nucleates” other genes, say, set *O*, such that *O* → *O*\* is the set that maximally cooperates with *I*. Let *S* = *I* ∪ *O*. The cooperativity of the *fixed* initial gene set *I* with *O* is defined as (c.f. Equation (S.26))

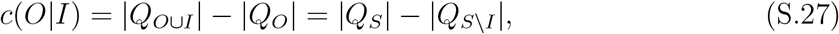

namely, the importance of *I* in producing a response when coupled with *O*. Note that *c*(*O*|*I*) > 0 ⇒ |*Q_O∪I_*| > |*Q_O_*|, meaning that *I* cooperates with *O* since they couple to produce a larger response than *O* along.

#### Cooperativity-based synergistic response

Here we describe how we compute the synergistic response (SR). Following the definition in Equation (S.27), let 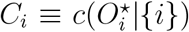 be the optimized cooperativity of gene *i* with other genes it nucleates (which we call 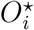). Consider gene *j* as the *query gene*. For every gene *i* ≠ *j* (treated as *sample genes*), we first compute the corresponding optimized cooperativity *C_i_*. We then compute their *pair* cooperativity 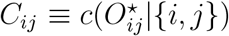, where 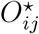 is the gene set nucleated by gene pair (*i*, *j*). The synergistic response for the (sample, query) pair (*i*, *j*) is defined as *s_ij_* = *C_ij_* − *C_j_*, i.e., the difference between the pair and singleton cooperativity. Finally, these quantities are averaged across MC realizations and the (sample, query) and (query, sample) synergistic response for each pair is averaged to obtain a symmetric synergistic response (SR) matrix with elements

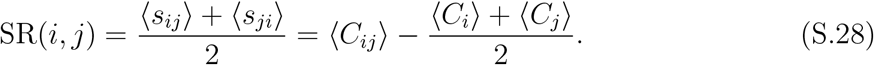

Note that 〈*C_ij_*〉 = 〈*C_ji_*〉 since they both correspond to nucleating genes with the same pair. In Fig. 8**c**, we show an example where gene *j* is treated as the query gene.

#### Assessing the significance of increased synergistic response for non-zero GI pairs

In Fig. 8**f**, we show that non-zero GI pairs tend to have high synergistic response. Here we develop a statistical procedure to assess its significance. Since we’re dealing with finite observations (i.e. histograms in Fig. 8**e**), this procedure begins by estimating the true density of synergistic response from empirical observations. It then performs Monte Carlo sampling to compute the probability of observing an synergistic response magnitude greater than a given threshold, which defines a *p*-value.

To set the notation straight, let 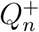 and 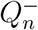 be the empirical distribution of SR magnitude for the positive and negative GI pairs, respectively (i.e. blue and green histograms in Fig. 8**e**). For a given threshold on the SR magnitude *∊*, define 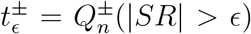. Let *P_n_* be the empirical distribution of SR magnitude for the zero GI pairs. With a slight abuse of notation, we denote the observed SR magnitude for these pairs (i.e. orange histograms in Fig. 8**e**) as *X*_1_.*X*_2_,…, *X_n_* ~ *P*, where *P* is the unknown true distribution. Define a statistic 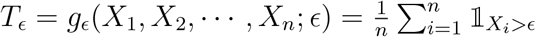 (i.e., fraction of zero GI pairs with SR magnitude greater than *∊*). Note that *T_∊_* = *T_∊_*(*P*) is a statistical functional. With these quantities defined, we compute the *p*-values as follows:

1. Estimate *P* with *P_n_* through resampling (e.g. bootstrapping or uniform sampling pushed through the inverse cumulative distribution of *P_n_*). Denote the estimated distribution as 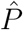.
2. From 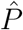, estimate the distribution of *T_∊_* through Monte Carlo. Call this distribution *P_T_∊__*.
3. Compute the *p*-value for the positive and negative GI pairs: *p*^±^ ≡ *P_T_∊__*(*T_∊_* ≥ *t*^±^). This is the probability of observing positive/negative GI pairs with SR magnitude at least as large as *∊*.

Fig. S17 summarizes the findings based on this procedure.

##### Algorithm S1 Pathway identification optimizing cooperativity with seed genes

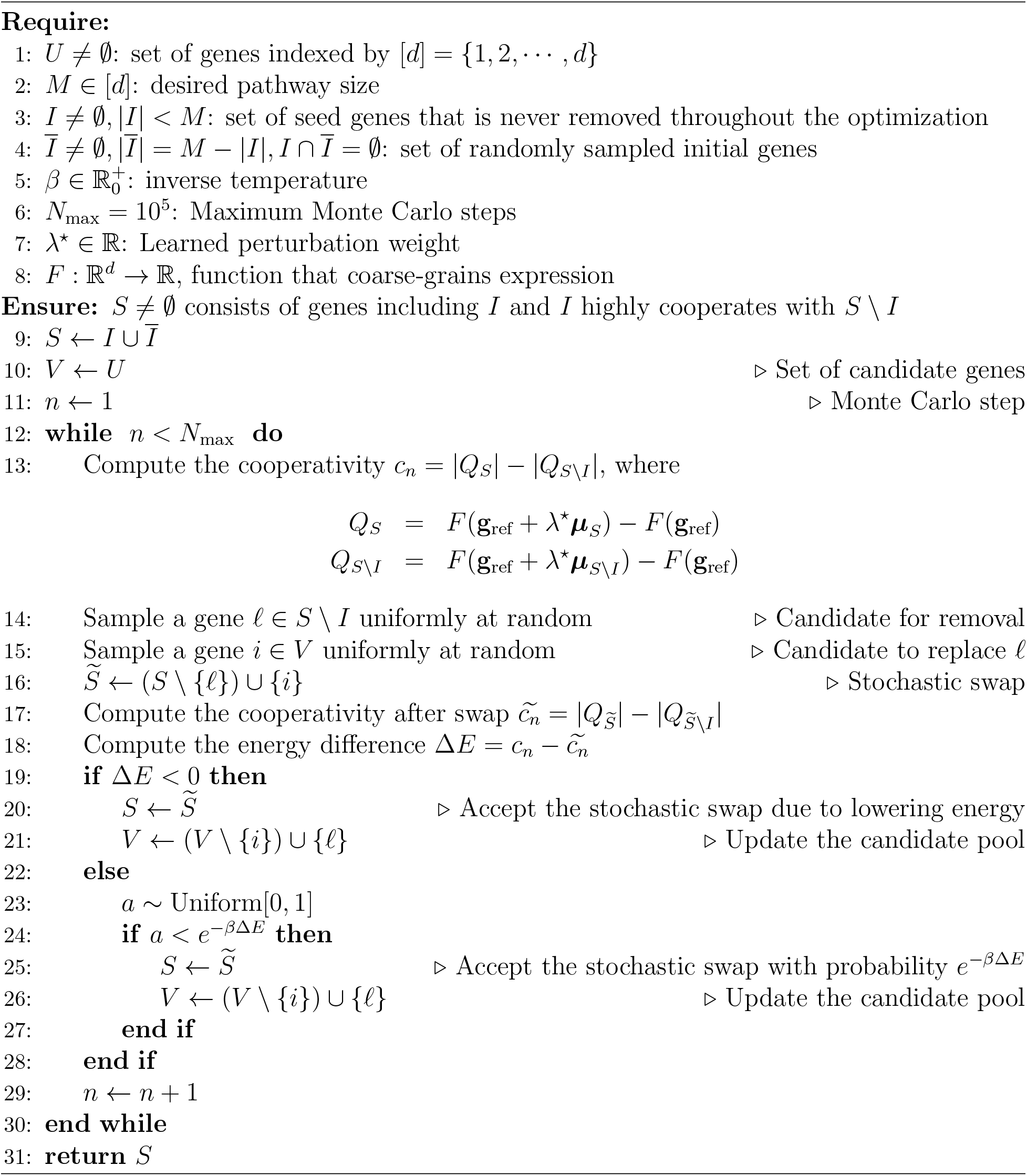

## Supplementary Figures and Tables

**Figure S1:**
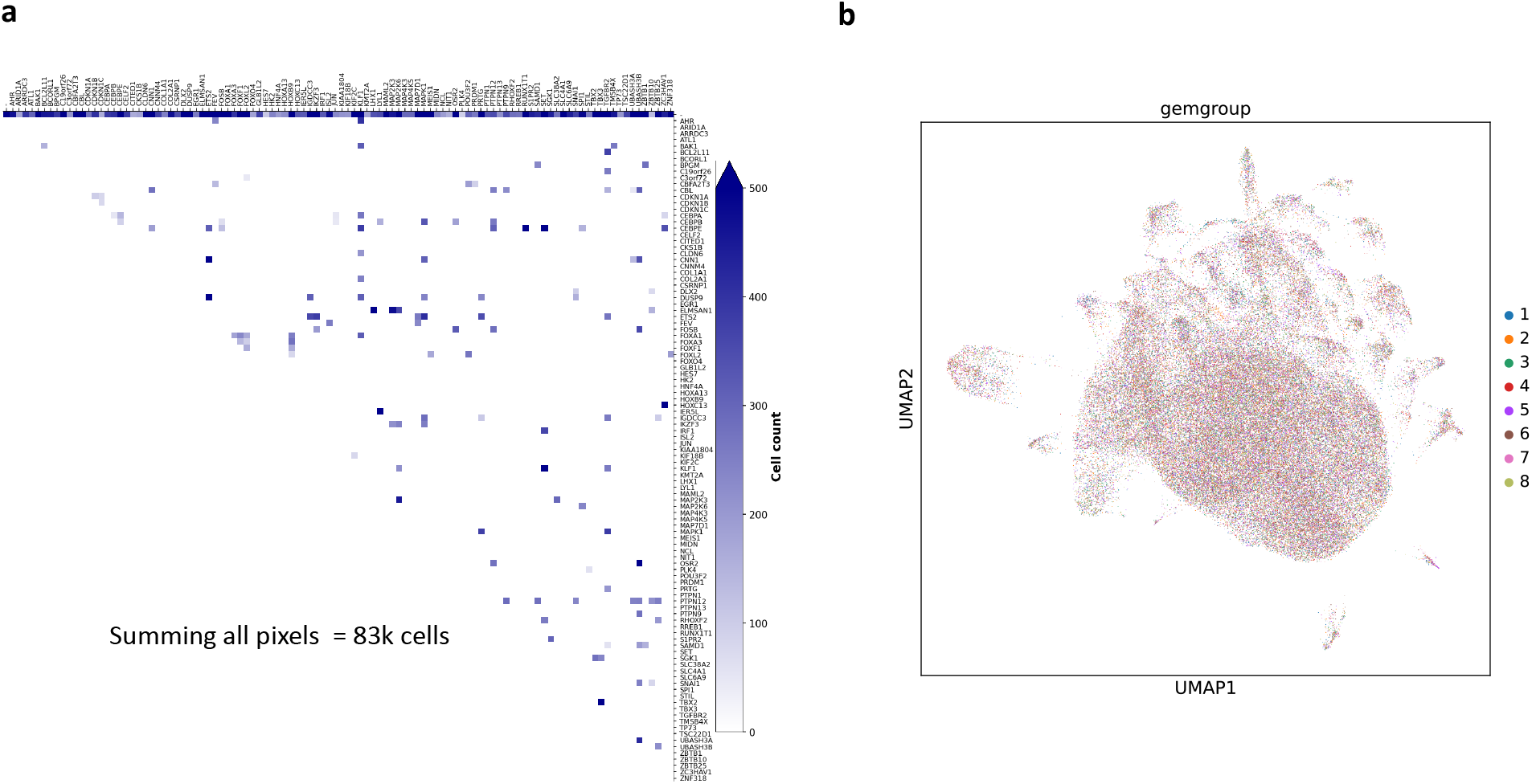
CRISPRa perturbation screen data used in this paper. Dataset is taken from [5] where single-cell RNA-sequencing pooled CRISPR activation screens were used to interrogate the combinatorial expression of genes. Due to technical limitations [5], only 155 single- and 132 double-CRISPRa transcriptional readouts were obtained across ~ 83, 000 cells. **a**: Combinatorial perturbation indicator matrix. Color indicates cell count (i.e., number of cells under specific perturbation condition). **b**: UMAP of single-cell expression profiles colored by *gemgroup* (i.e. corresponding to batches in the 10x experiment). Mixing of the colors indicates minimal batch effect in the dataset.

**Figure S2:**
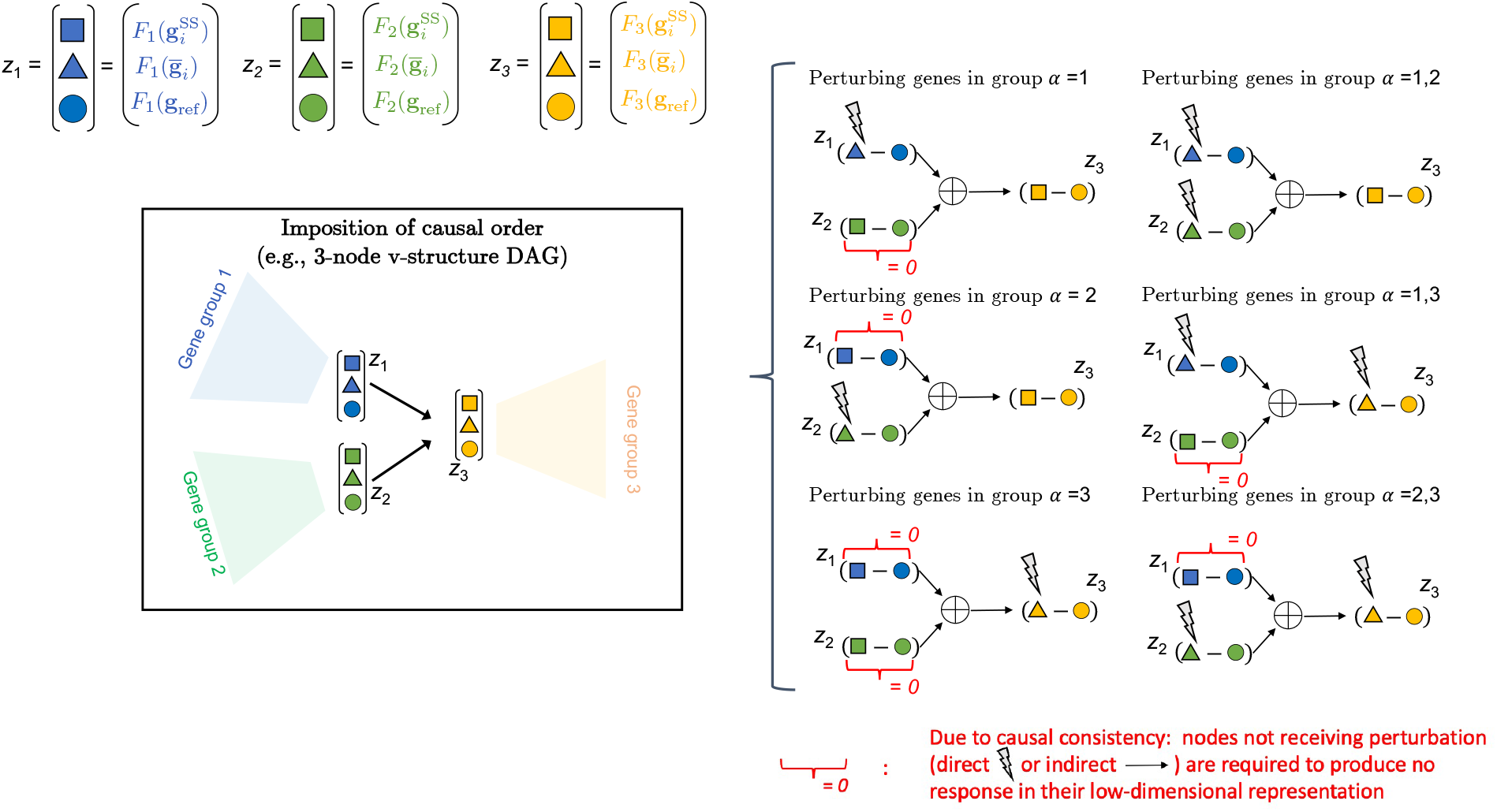
Imposition of causal order: perturbation propagation. Based on Fig.2, here we enumerate all possible cases of perturbation to illustrate the the imposition of causal order. When upstream nodes are perturbed, their quenched responses, defined as the difference between their quenched and reference low-dimensional values of expression (i.e. 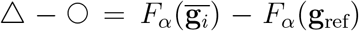, for *α* = 1,2) are summed together and propagated to match the observed response of the downstream node (i.e. 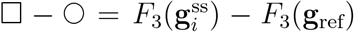). For nodes receiving neither direct perturbation in experiments (indicated by the lightening marker) nor through propagated response from their upstream nodes (indicated by arrow), we require their observed response to be zero. Nodes that are subjected to this requirement are indicated by red curly brackets.

**Figure S3:**
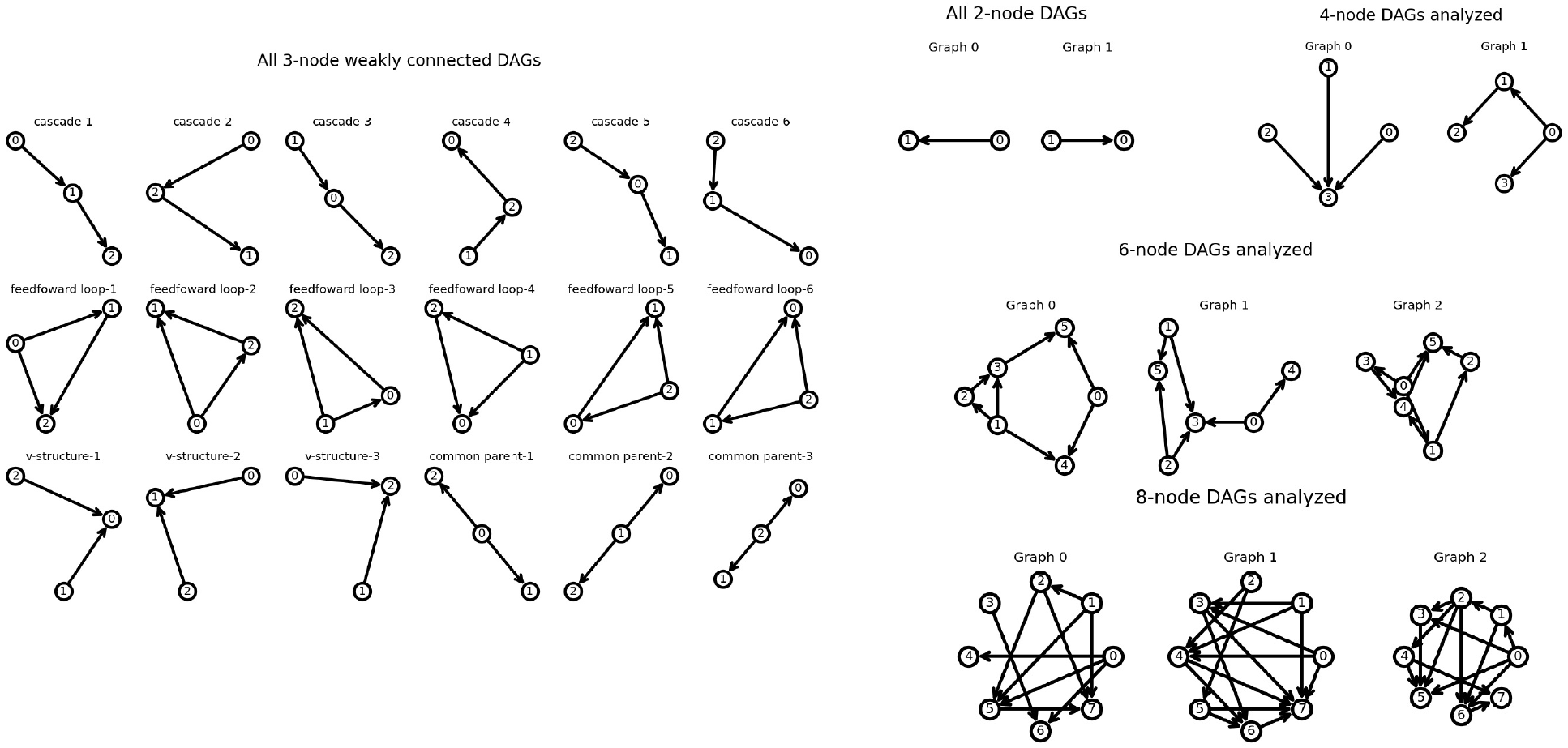
DAGs analyzed in this paper. We enumerate all possible weakly connected 2-node and 3-node DAGs and randomly survey a few representative 4-node, 6-node, and 8-node DAGs. These graphs are used to train multiple Lowdeepredict models with various random gene placement schemes. Results are summarized in Fig.3. Note that 4-node and 6-node graphs shown here are chosen at random.

**Figure S4:**
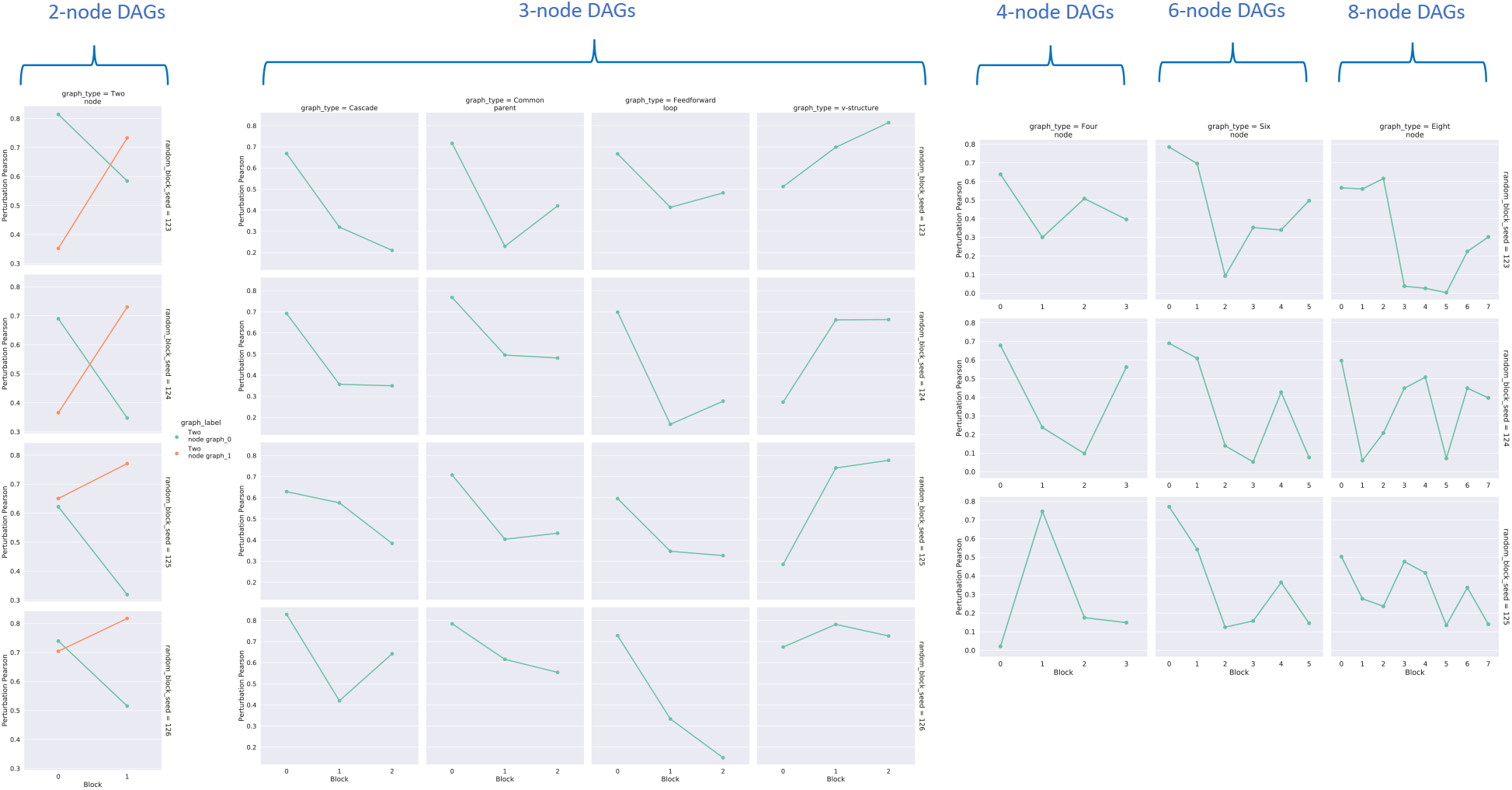
Node-wise validation predictive performance for the best model in each category. Shown here is the node-wise performance breakdown of the best model in each graph category in Fig.3. For all panels, columns indicate graph topology while rows represent random gene bucketing/placement.

**Figure S5:**
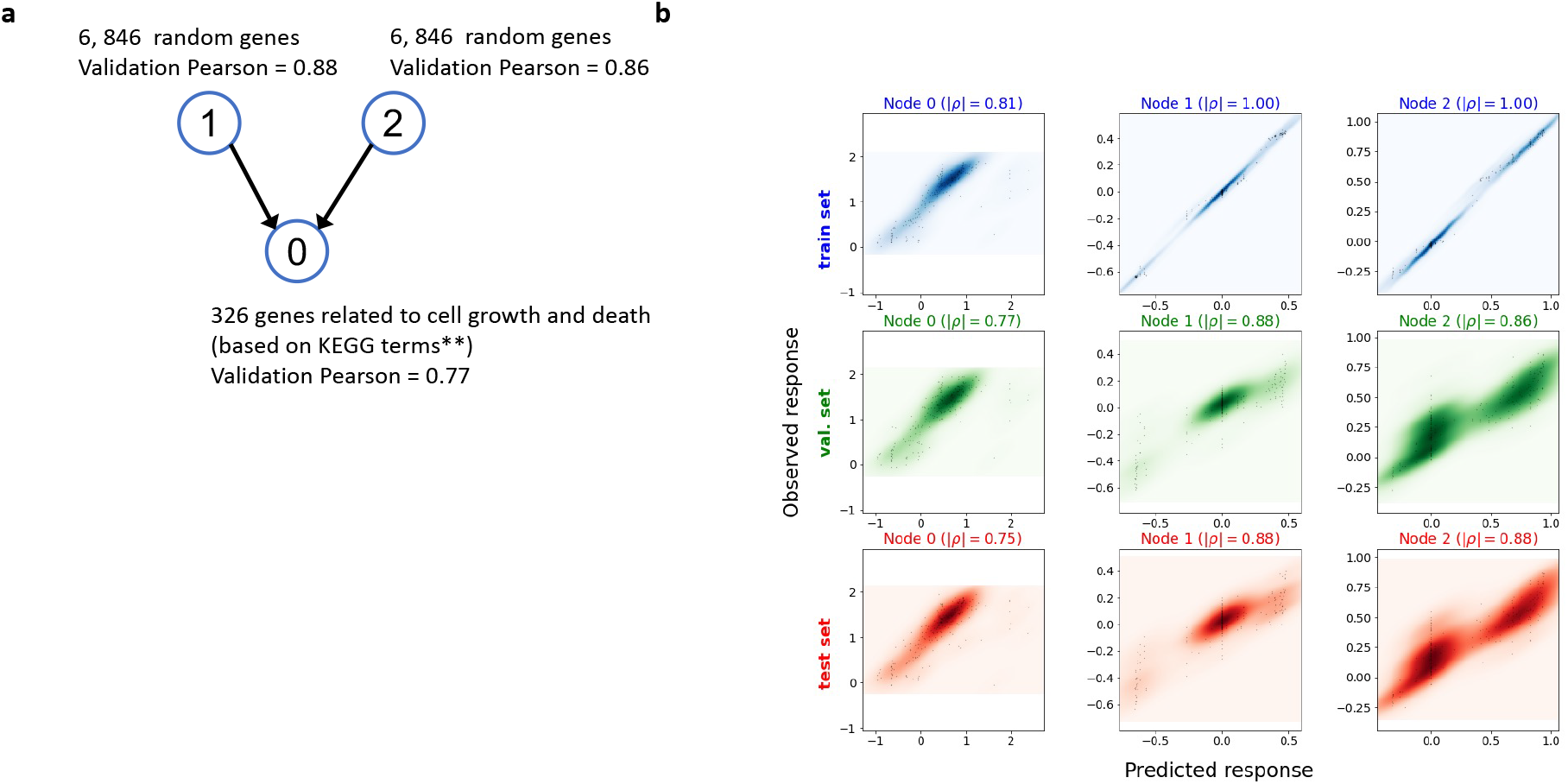
Predicted performance of the model analyzed in 6. **a**: Causal graph of the model analyzed in FIG. 6. **b**: Observed response is plotted against the predicted response for this model. Color indicates density estimated with Gaussian kernels. Columns are labeled by block/node number corresponding to the graph in the inset of **a**. Top row shows the result for training data, the middle row shows that for the validation data, and the bottom shows that for the test data. Node-wise Pearson’s correlation magnitude |*ρ*| is indicated for each node and data type (train or validation).

**Figure S6:**
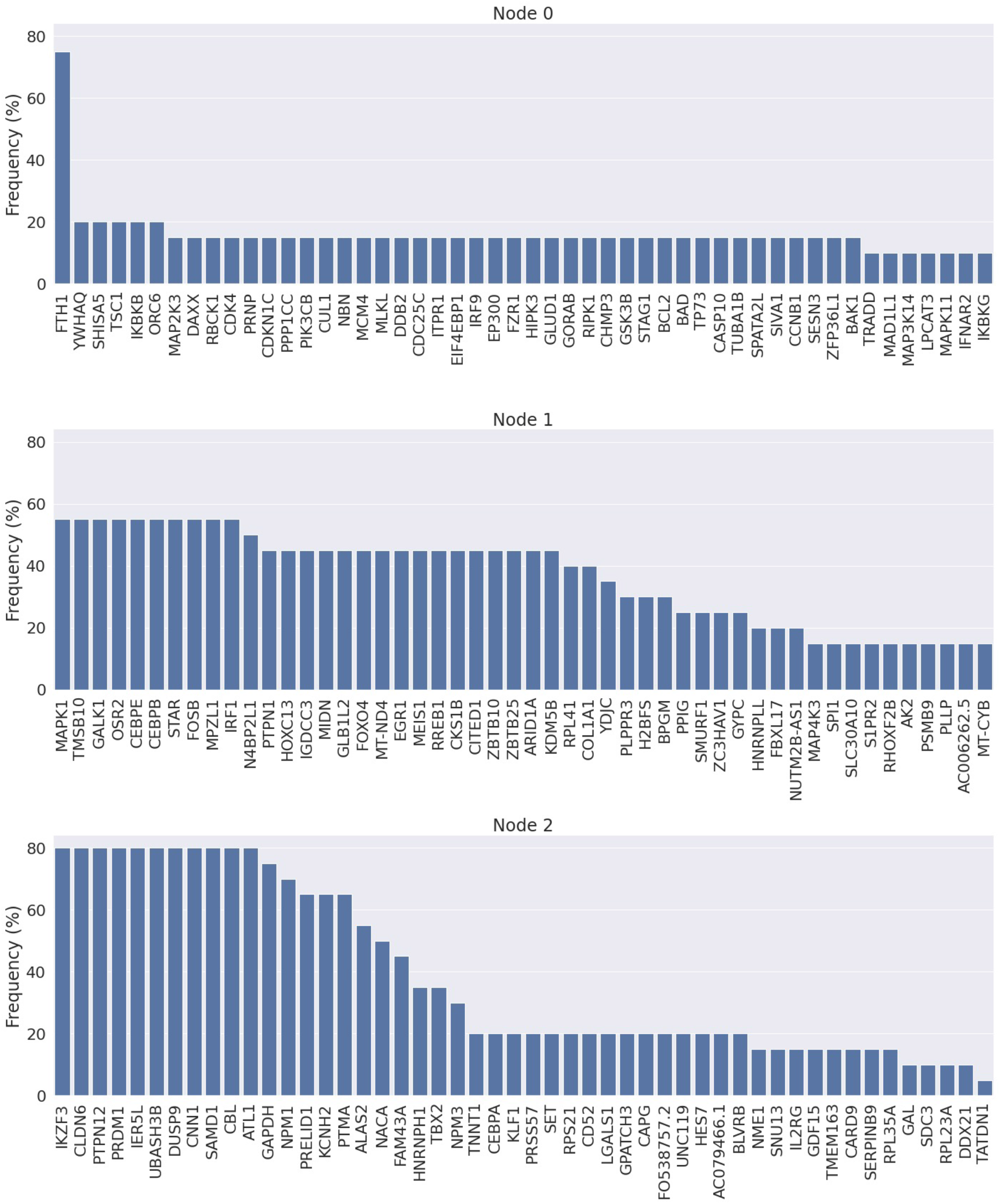
Frequently identified genes by Monte Carlo. As described in FIG. 6, here we list the top-50 frequently identified genes for each node using Monte Carlo (MC). Note that each gene can at most be identified 20 times (since there are 20 independent MC runs), so a gene attaining frequency 100% means it is identified in all 20 MC runs. Genes of higher frequency are those imputed to be important by our model to affect cell growth and death, see Fig. 6 **a**.

**Figure S7:**
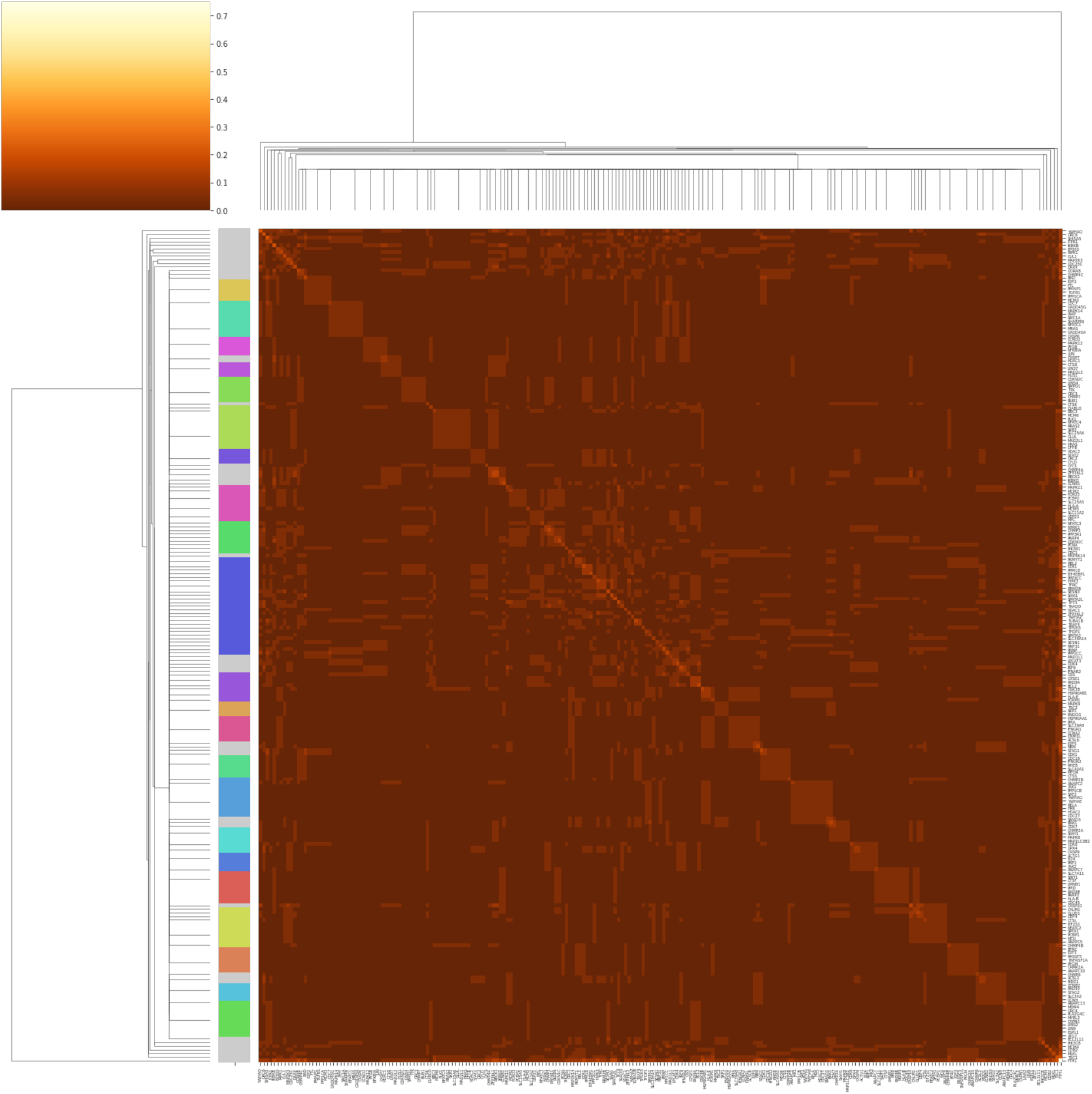
Co-occurrence frequency matrix of gene sets identified by Monte Carlo for node 0. The 20 sets of 20 genes identified by Monte Carlo, see Fig. 6, are pooled together to keep only the unique genes. The unperturbed expression of genes in each node are used to compute the frequency of pair-wise co-occurrence (see *Methods* for details). A pair of gene attaining frequency 1 means both genes are present in all of the 20 gene sets. Based on the frequency of co-occurrence matrix, genes are clustered using correlation as distance metric. Shown here is the cluster heat map for node 0. Color indicates correlation.

**Figure S8:**
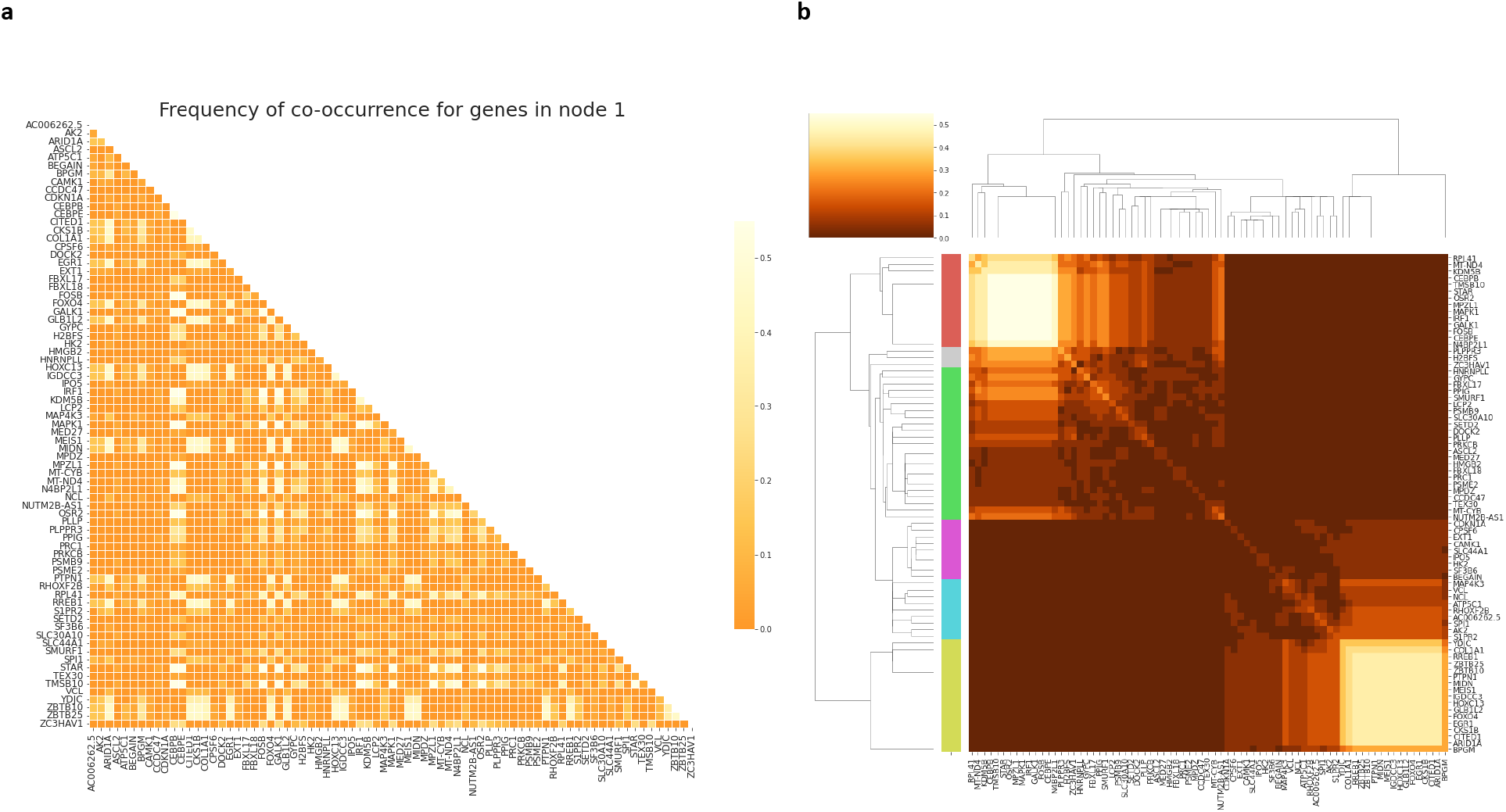
Co-occurrence frequency matrix of gene sets identified by Monte Carlo for node 1. Similar to FIG. S7, shown here are the co-occurrence frequency matrix and cluster heat map for node 1.

**Figure S9:**
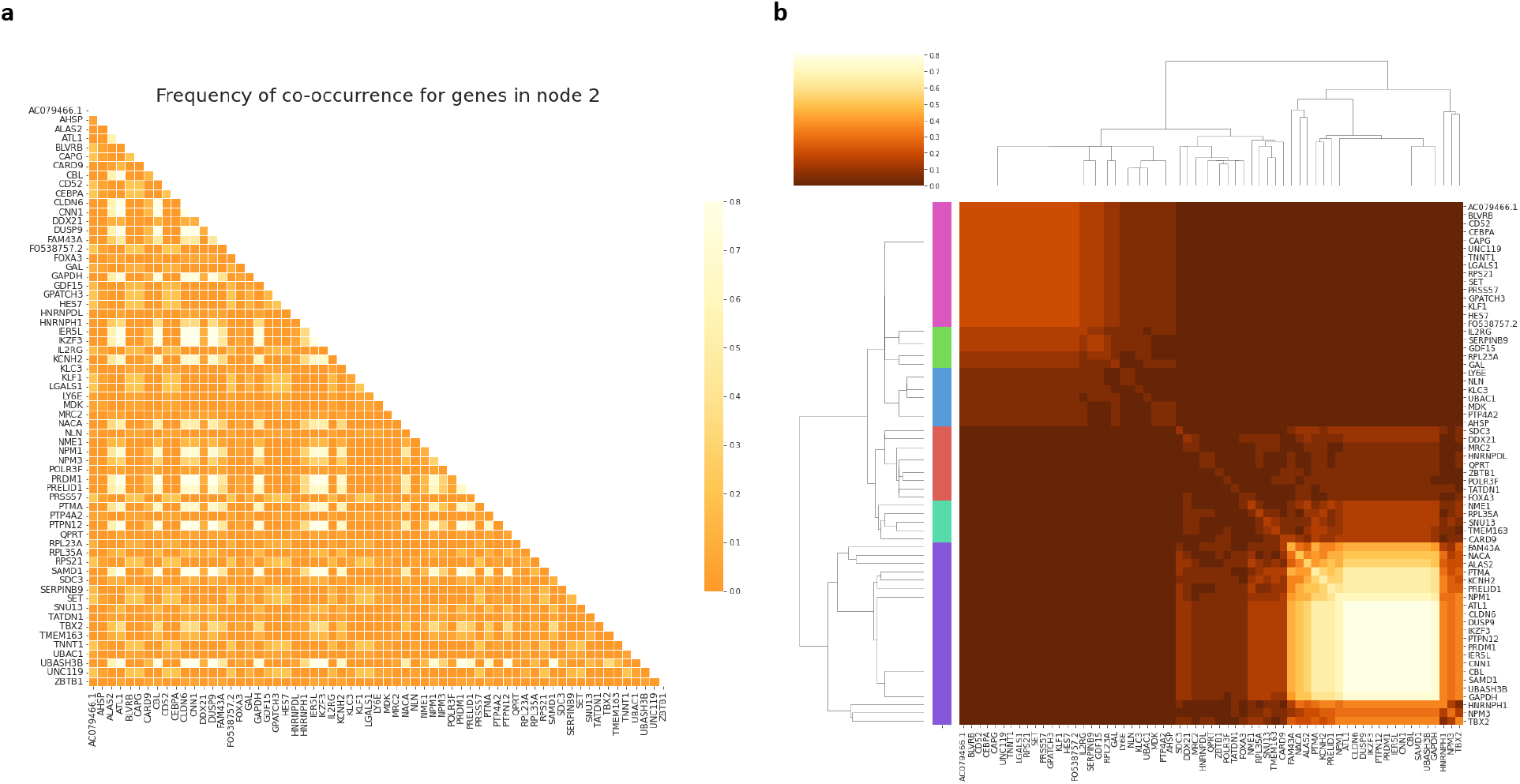
Co-occurrence frequency matrix of gene sets identified by Monte Carlo for node 2. Similar to FIG. S7, shown here are the co-occurrence frequency matrix and cluster heat map for node 2.

**Figure S10:**
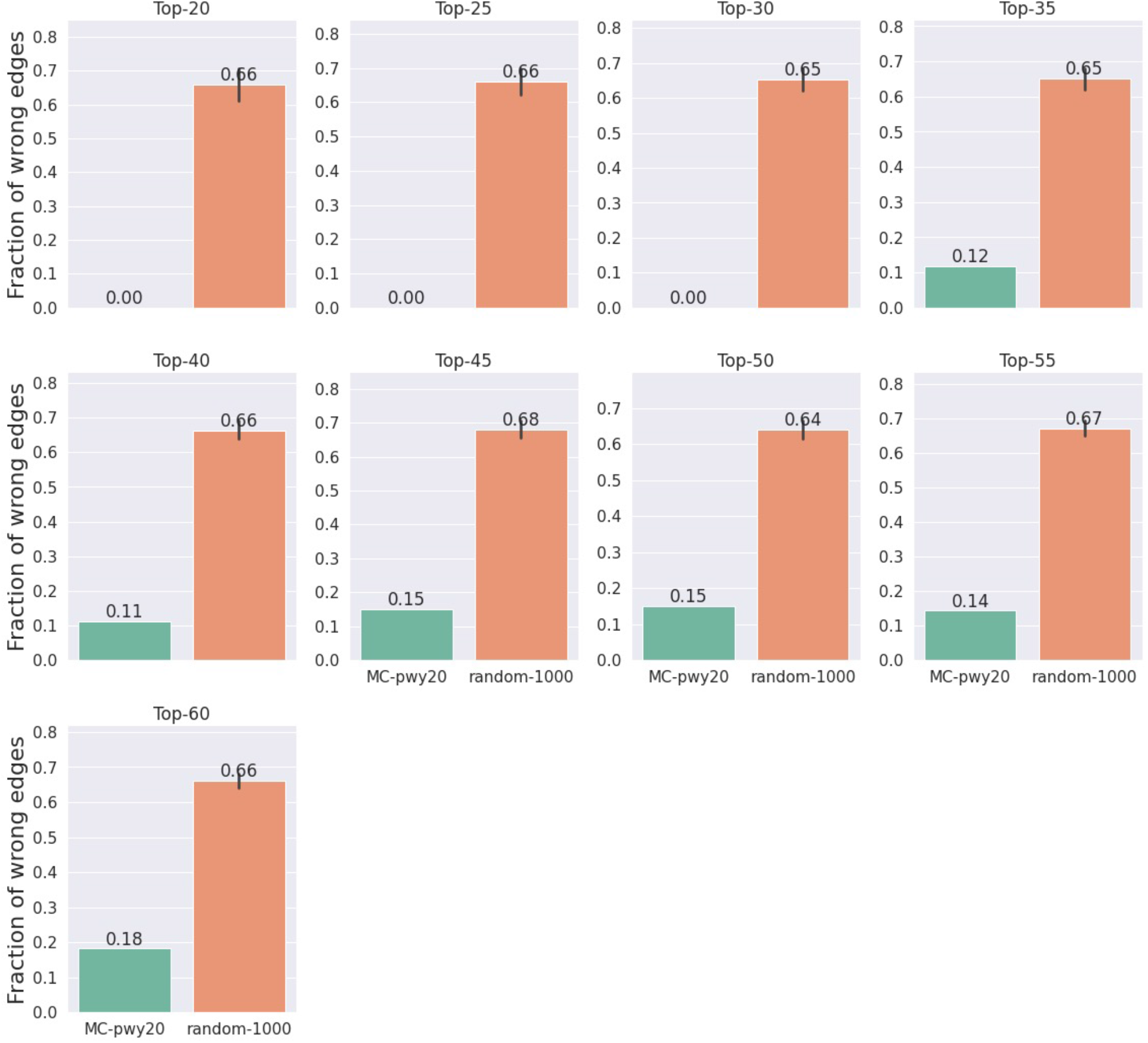
Fraction of anti-causal edges. Similar to Fig. 7, here we show the fraction anti-causal edges in (i) *G*_known_ (i.e. gene-gene interaction network constructed using the top-*k* MC genes, labeled *MC-pwy20*) and (ii) random graphs (*random-1000*). For each panel (i.e. *k* as in top-*k*), we generate an ensemble of 1,000 graphs, each with *k* nodes sampled from all the genes in the CRISPRa dataset. The construction of random graphs from these genes is as described in *Methods*.

**Figure S11:**
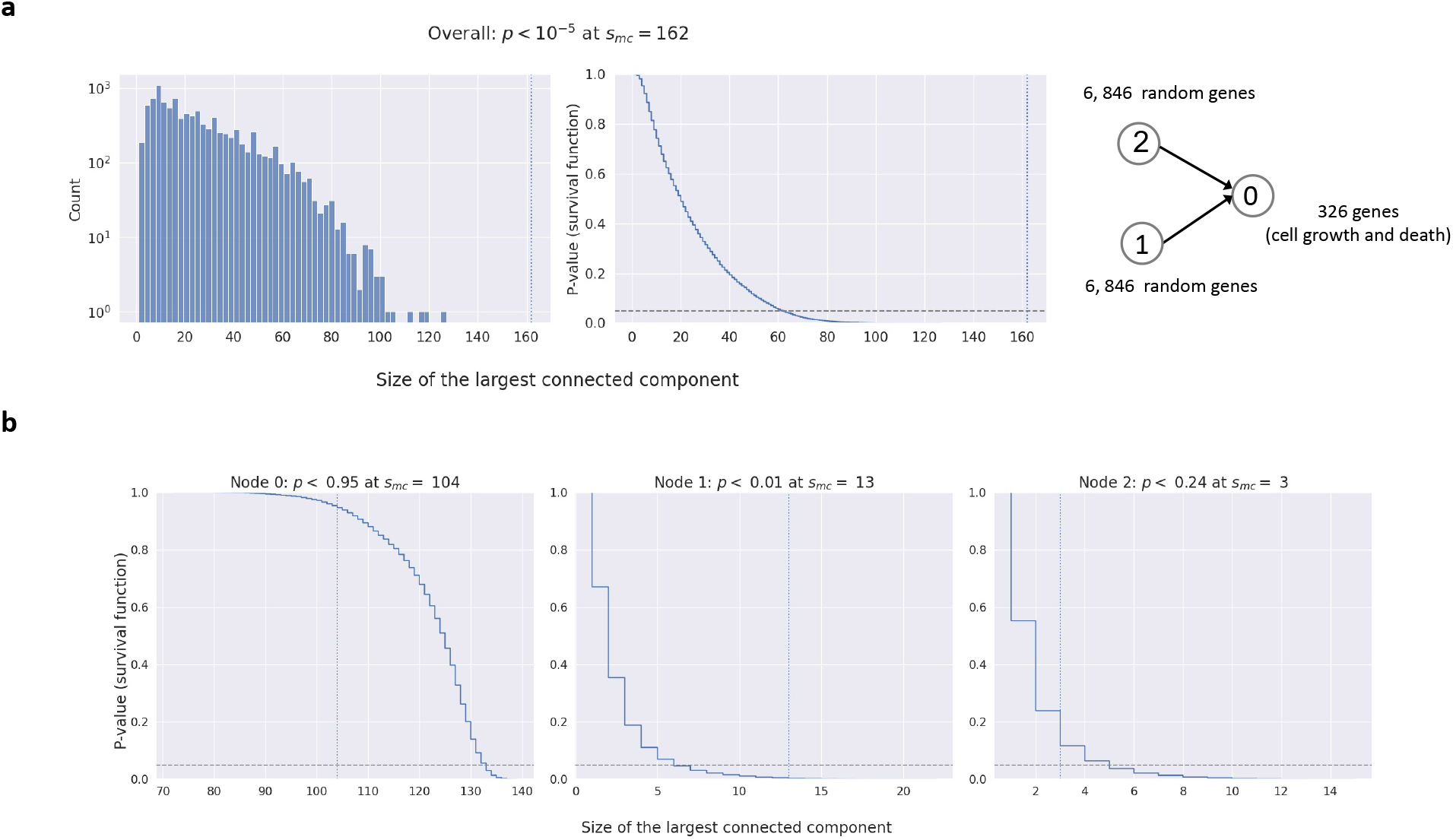
Computing the *p*-value for causal subgraphs to gauge their connectivity compared to random graphs. **a**: As described in *Methods*, here we show the distribution of the largest connected component size *ŝ* (left) and the *p*-value for having a graph with largest connected component larger than *ŝ* (i.e. *Pr*(*s* ≥ *ŝ*), or the survival function, shown on the right), estimated based on randomly sampling 100k graphs of 267 genes (i.e., number of nodes in *G*_mc_ for pathway size 20, see *Methods* and Table S5) from the those available in the CRISPRa dataset [5]. In the right panel, *p*-value threshold of 0.05 is indicated by the horizontal dashed line. The vertical dotted line shows the largest connected component size for *G*_mc_, *s*_mc_. **b**: Similar to **a**, except that the sampling is done node-wise. *s*_mc_ for each node is indicated by the vertical dashed lines and labeled.

**Figure S12:**
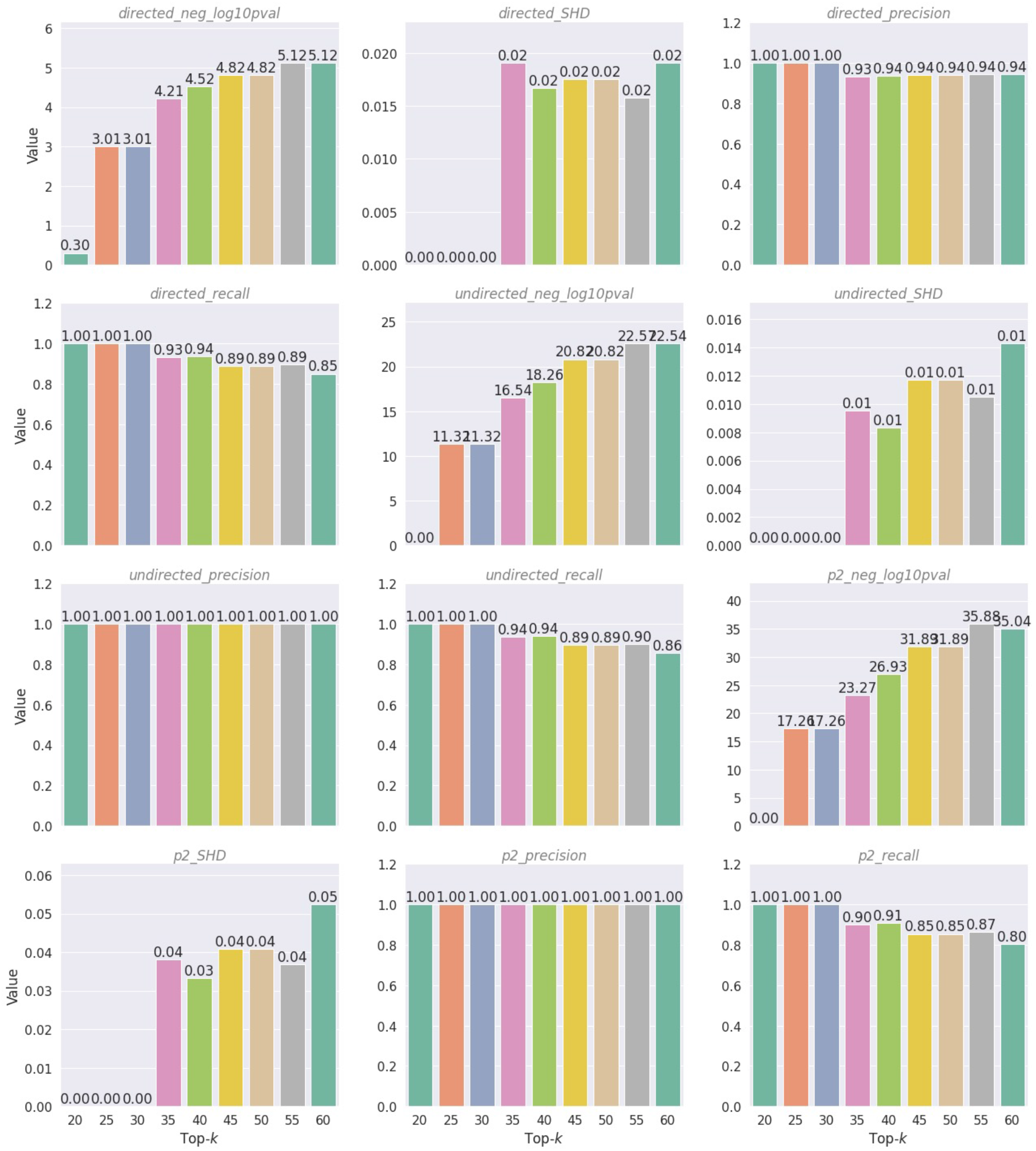
Causal graph identification performance using the top-*k* MC genes described in Fig. 7. These performance metrics are evaluated by taking the reconstructed graph (e.g. those in Fig. 7) as the “predicted graph” and the pruned graph (i.e. those in Fig. 7 with the red edges removed) as the “reference graph”. SHD: structurally hamming distance, P2: binary neighborhood. For the definition of directed and undirected *p*-value, see *Supplementary Methods* for details.

**Table S1:** Evaluation of causal loss for the example in Fig. 2c,d and S2. Queched response **Δ**, predicted response 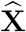, and the causal loss function are defined in the main text and in Table S2.

| Node(s) perturbed | $\Delta$ | $\hat{\mathbf{X}}$ | Causal loss $\ell_i$ |
| --- | --- | --- | --- |
| 1 | $\begin{pmatrix} \Delta_1 \\ 0 \\ 0 \end{pmatrix}$ | $\begin{pmatrix} \Delta_1 \\ 0 \\ \Delta_1 \end{pmatrix}$ | $(F_1(\mathbf{g}_i^{\text{ss}}) - F_1(\bar{\mathbf{g}}_i))^2$<br>$+(F_2(\mathbf{g}_i^{\text{ss}}) - F_2(\mathbf{g}_{\text{ref}}))^2$<br>$+(F_3(\mathbf{g}_i^{\text{ss}}) - F_3(\mathbf{g}_{\text{ref}}) - \Delta_1)^2$ |
| 2 | $\begin{pmatrix} 0 \\ \Delta_2 \\ 0 \end{pmatrix}$ | $\begin{pmatrix} 0 \\ \Delta_2 \\ \Delta_2 \end{pmatrix}$ | $(F_1(\mathbf{g}_i^{\text{ss}}) - F_1(\mathbf{g}_{\text{ref}}))^2$<br>$+(F_2(\mathbf{g}_i^{\text{ss}}) - F_2(\bar{\mathbf{g}}_i))^2$<br>$+(F_3(\mathbf{g}_i^{\text{ss}}) - F_3(\mathbf{g}_{\text{ref}}) - \Delta_2)^2$ |
| 3 | $\begin{pmatrix} 0 \\ 0 \\ \Delta_3 \end{pmatrix}$ | $\begin{pmatrix} 0 \\ 0 \\ \Delta_3 \end{pmatrix}$ | $(F_1(\mathbf{g}_i^{\text{ss}}) - F_1(\mathbf{g}_{\text{ref}}))^2$<br>$+(F_2(\mathbf{g}_i^{\text{ss}}) - F_2(\mathbf{g}_{\text{ref}}))^2$<br>$+(F_3(\mathbf{g}_i^{\text{ss}}) - F_3(\bar{\mathbf{g}}_i))^2$ |
| 1, 2 | $\begin{pmatrix} \Delta_1 \\ \Delta_2 \\ 0 \end{pmatrix}$ | $\begin{pmatrix} \Delta_1 \\ \Delta_2 \\ \Delta_1 + \Delta_2 \end{pmatrix}$ | $(F_1(\mathbf{g}_i^{\text{ss}}) - F_1(\bar{\mathbf{g}}_i))^2$<br>$+(F_2(\mathbf{g}_i^{\text{ss}}) - F_2(\bar{\mathbf{g}}_i))^2$<br>$+(F_3(\mathbf{g}_i^{\text{ss}}) - F_3(\mathbf{g}_{\text{ref}}) - \Delta_1 - \Delta_2)^2$ |
| 1, 3 | $\begin{pmatrix} \Delta_1 \\ 0 \\ \Delta_3 \end{pmatrix}$ | $\begin{pmatrix} \Delta_1 \\ 0 \\ \Delta_1 + \Delta_3 \end{pmatrix}$ | $(F_1(\mathbf{g}_i^{\text{ss}}) - F_1(\bar{\mathbf{g}}_i))^2$<br>$+(F_2(\mathbf{g}_i^{\text{ss}}) - F_2(\mathbf{g}_{\text{ref}}))^2$<br>$+(F_3(\mathbf{g}_i^{\text{ss}}) - F_3(\bar{\mathbf{g}}_i) - \Delta_1)^2$ |
| 2, 3 | $\begin{pmatrix} 0 \\ \Delta_2 \\ \Delta_3 \end{pmatrix}$ | $\begin{pmatrix} 0 \\ \Delta_2 \\ \Delta_2 + \Delta_3 \end{pmatrix}$ | $(F_1(\mathbf{g}_i^{\text{ss}}) - F_1(\mathbf{g}_{\text{ref}}))^2$<br>$+(F_2(\mathbf{g}_i^{\text{ss}}) - F_2(\bar{\mathbf{g}}_i))^2$<br>$+(F_3(\mathbf{g}_i^{\text{ss}}) - F_3(\bar{\mathbf{g}}_i) - \Delta_2)^2$ |

**Table S2:**
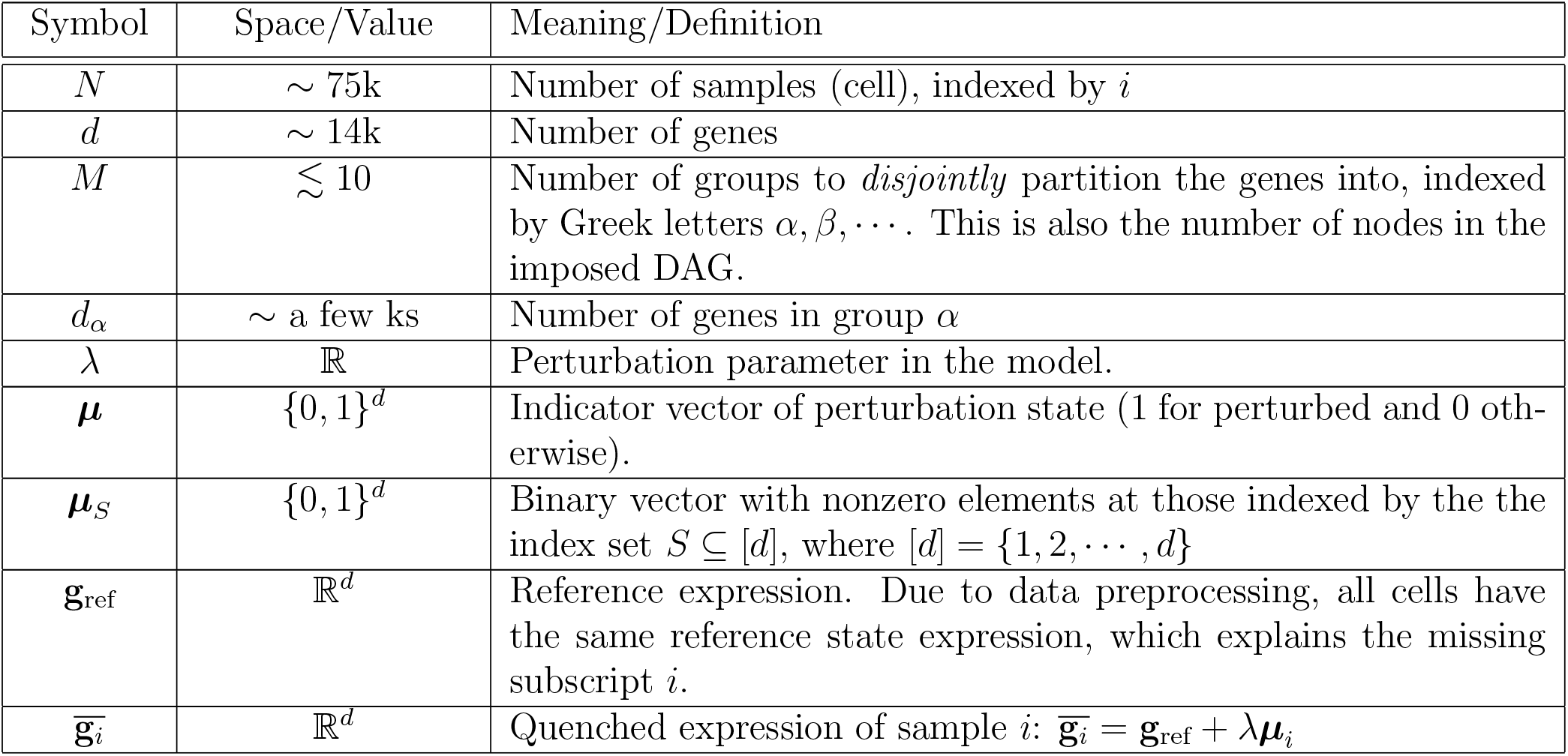
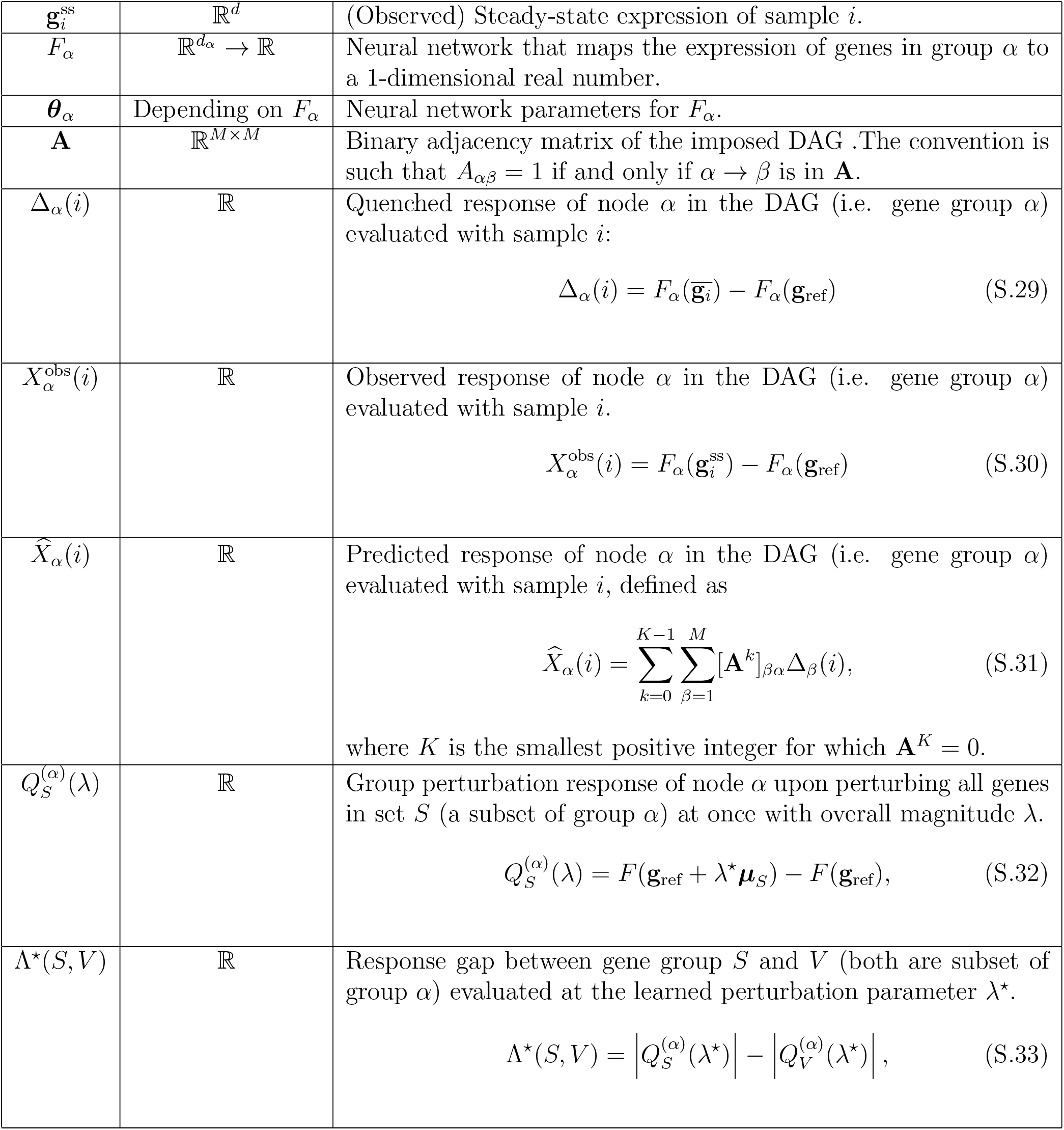
Table summarizing the notation used in the main text.

**Figure S13:**
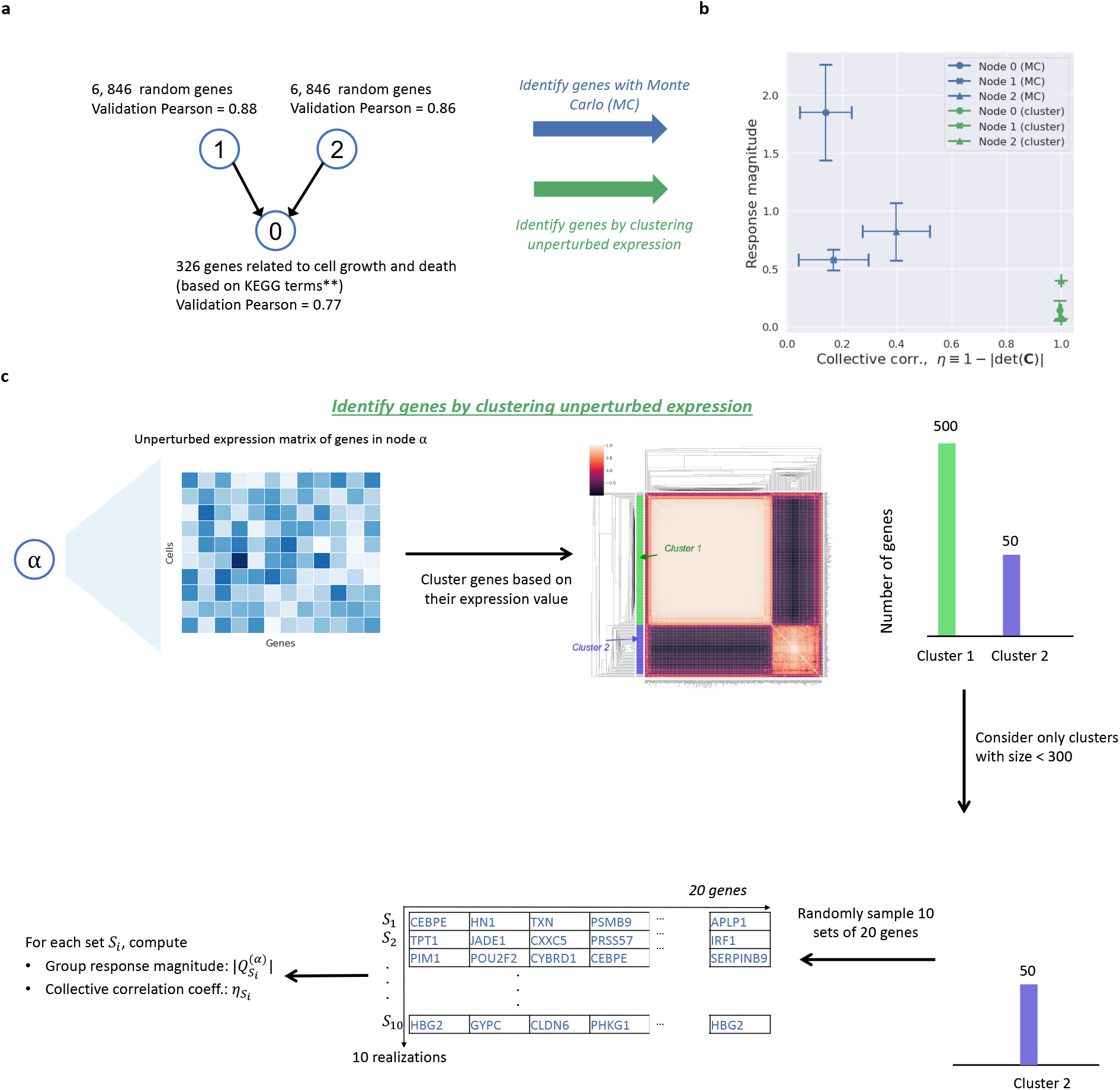
MC identifies gene sets that are distinct from co-expression clusters. **a**: Causal graph and gene assignment based on the discussion in Fig. 6. **b**: Group response magnitude and collecitve correlation of gene sets identified by MC or co-expression clustering. Markers indicate node in the causal graph in **a** while colors correspond to identification method (blue for MC and green for co-expression clustering). **c**: Procedure to cluster genes based on their expression. Co-expression clusters identified are then sampled to compute their collective correlation coefficient, *η* = 1 − |det(**C**_*i*_)| and the group response magnitude. See *Supplementary Methods* for a discssion on *η*.

**Figure S14:**
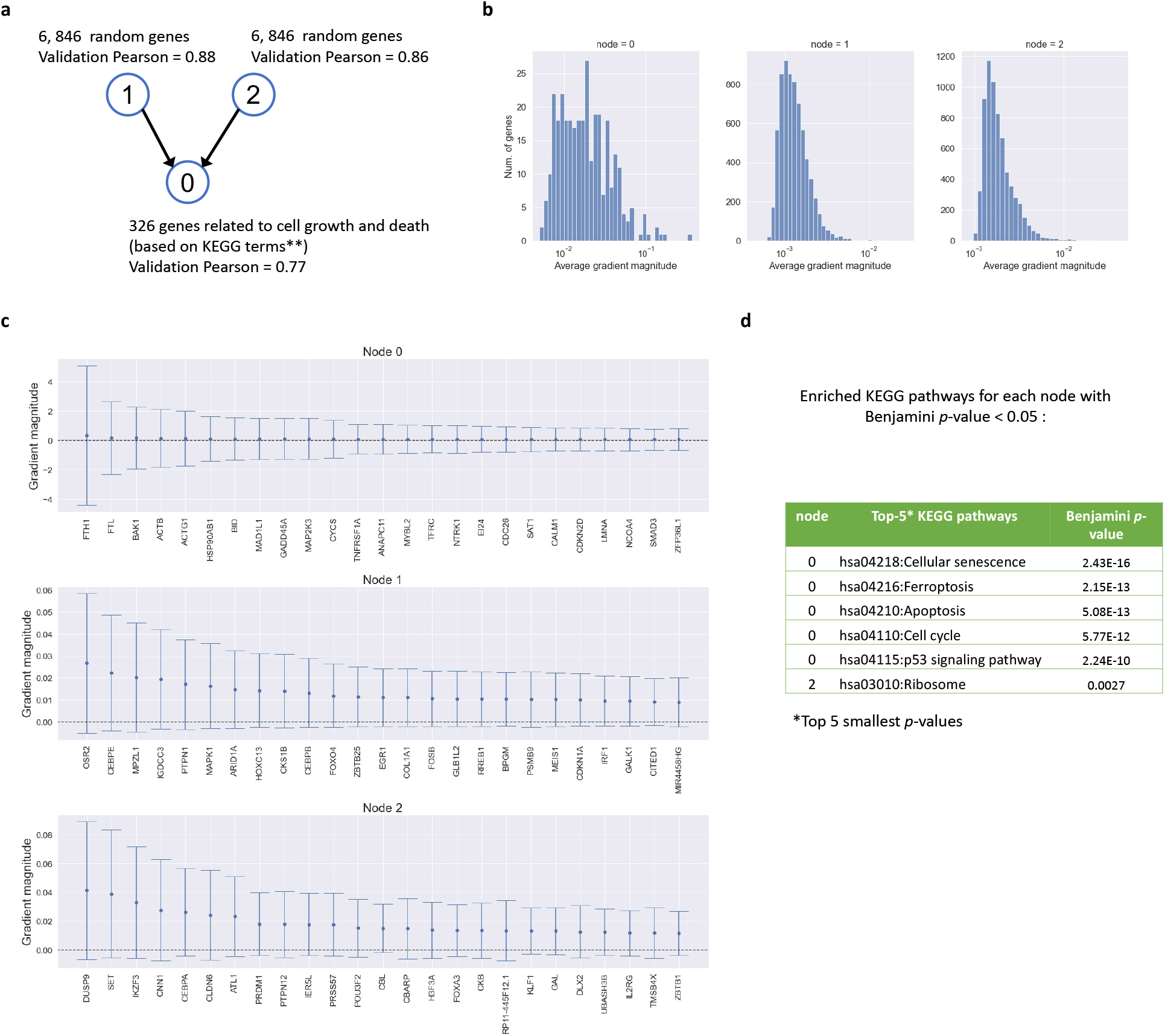
Gradients of the learned function *F_α_* and the KEGG functional enrichment analysis of genes selected based on high gradients. **a**: Causal graph of the model analyzed in Figure 6. **b**: Distribution of the average gradient magnitude for each node. The gradient of function *F_α_* for gene *j* in node *α* evaluated on the steady-state expression of cell *i* is defined as 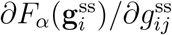. The average is taken over the validation data (*i* ∈ validation). **c**: Top-25 genes in each node based on the average gradient magnitude 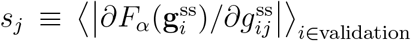 **d:** The top-25 genes for each node shown in **c** are used to perform functional enrichment ananlysis. Significantly enriched KEGG pathways (with Benjamini *p*-value < 0.05) are shown for each node. For simplicity, we only show results with 5 smallest *p*-values (all below the 0.05 threshold). Note the absence of significantly enriched pathways for node 1. For node 2, only 1 pathway has *p* < 0.05.

**Figure S15:**
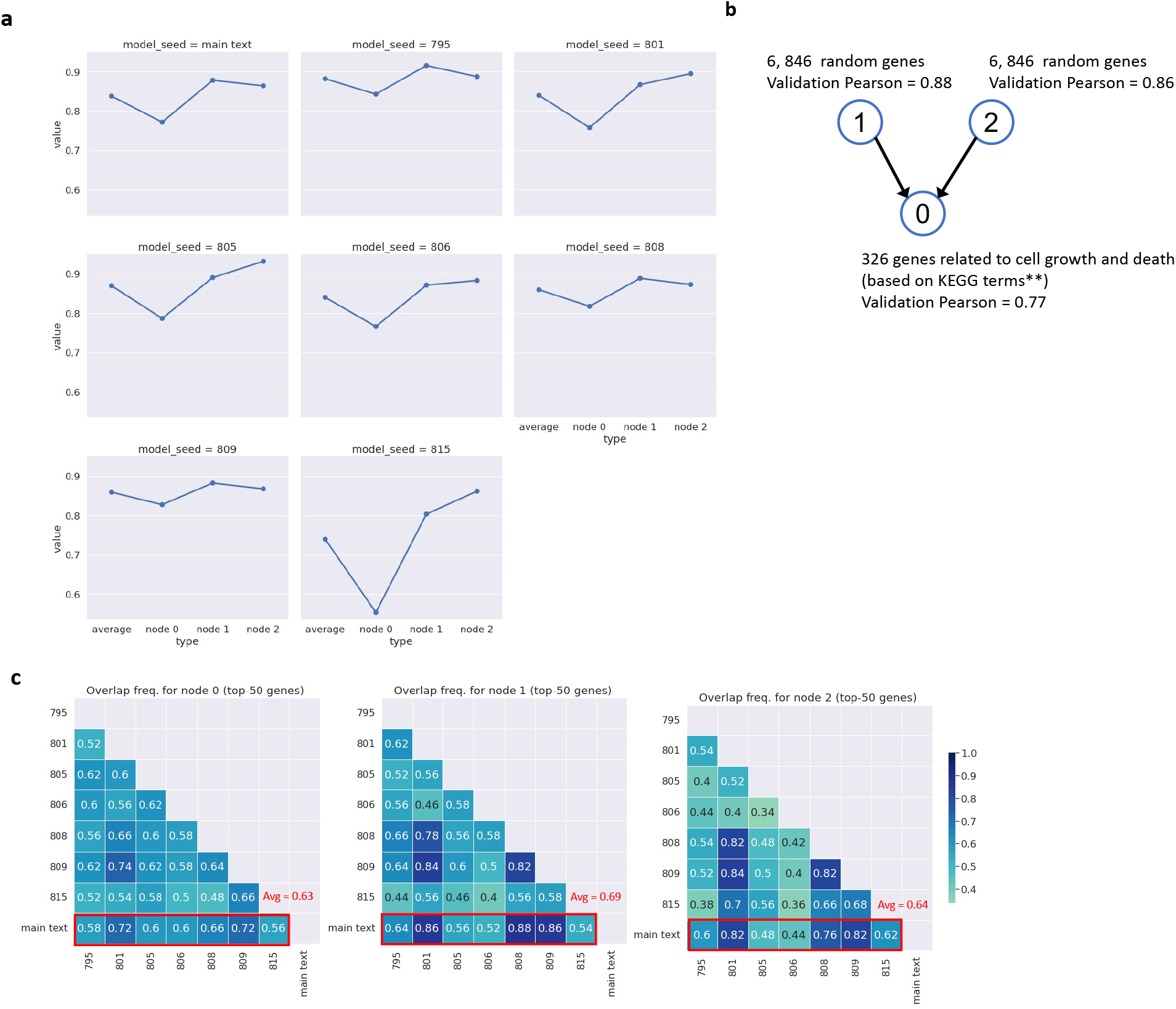
Gene sets identified by Monte Carlo using models with similar performance under the same neural network architecture and gene group assignment schemes shows good consistency. **a**: Pearson correlation between the predicted and observed response for each node in the causal graph shown in **b** as well as their average. Model seed indicates the random seed used to initialize the optimize neural networks. *Main text*: model presented in Fig. 6 and 7. **c**: Overlap frequency between the top-50 genes identified by different models.

**Figure S16:**
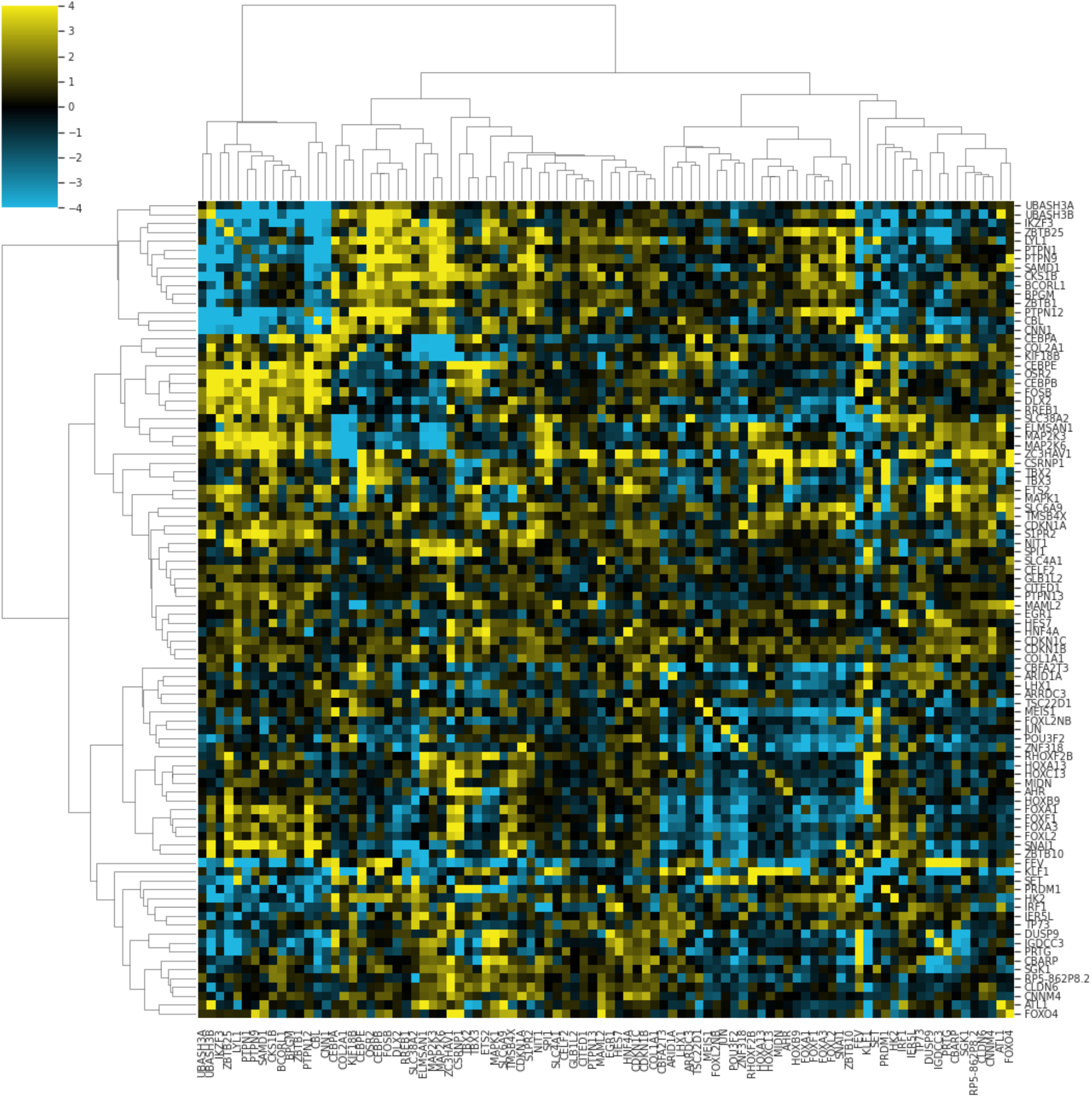
GI map from [5]. Shown here is the genetic interaction (GI) map from the CRISPRa Perturb-seq screen reported in [5]

**Figure S17:**
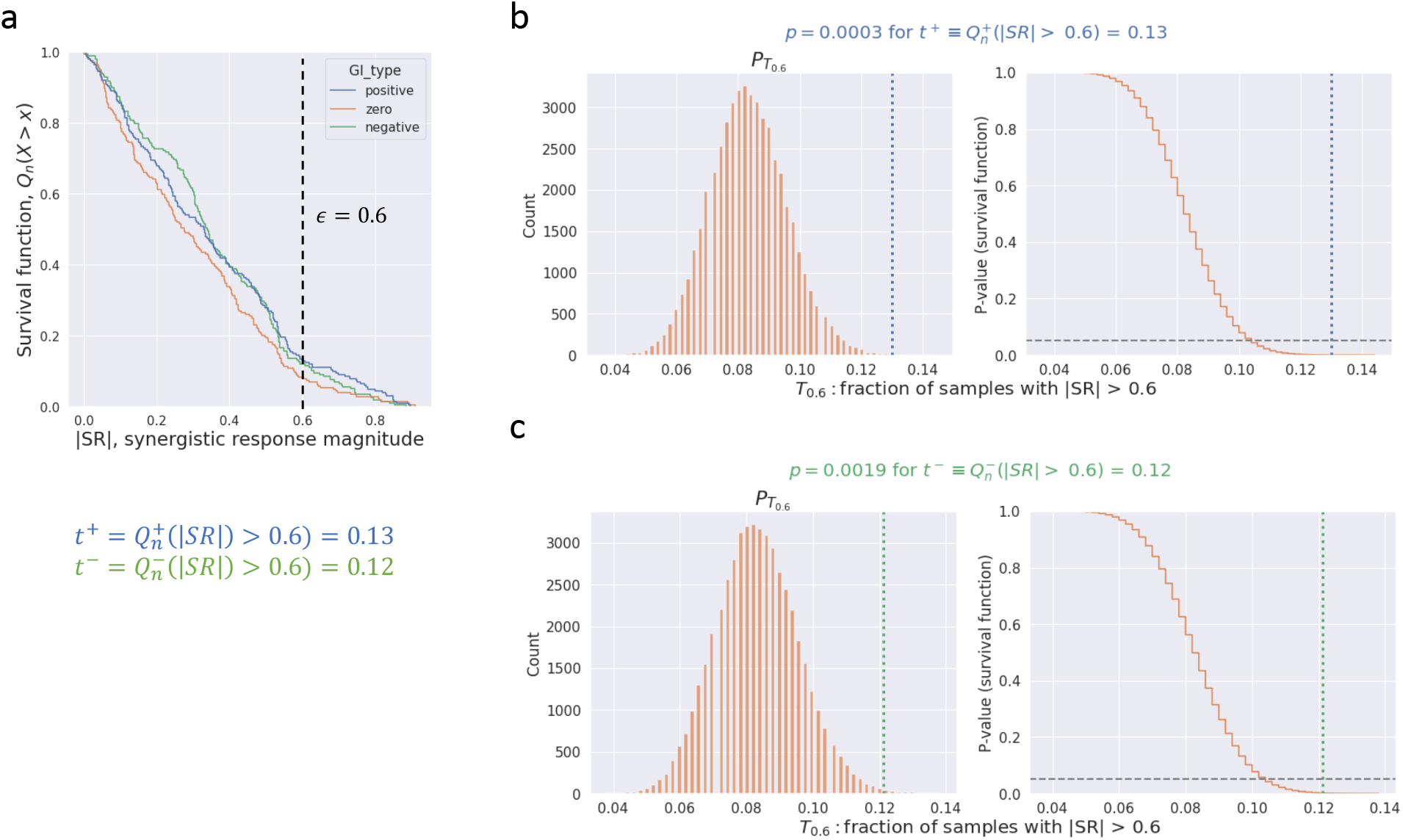
Gene sets nucleated with high GI pairs tend to have high synergisitc response. **a** Empirical survival function of synergistic response magnitude, 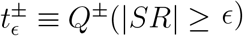 (blue and green). For convenience, we also show such distribution for zero GI pairs (orange). Note that at *∊* = 0.6, *t*^+^ = 0.13 and *t*^−^ = 0.12. **b**: On the left shows the distribution of the statistic *T_∊_* for the zero GI pairs estimated with Monte Carlo sampling. Blue dashed line indicates *t*^+^ at *e* = 0.6 for positive GI pairs. On the right shows the survival function for *T_∊_*: *P*(*T_∊_* ≥ *t*). At *t* = *t*^+^ = 0.13, we have a *p*-value of *p*^+^ = *P_T_∊__(*T_∊_* ≥ *t*^+^) = 0.0003. **c**: Similar to **b** except that the green dashed line on the left indicates *t*^−^* = 0.12 at *∊* = 0.6 for negative GI pairs. The corresponding *p*-value is *p*^−^ = *P_T_∊__* (*T_∊_* ≥ *t*^−^) = 0.0019. In **b,c**, we resample from the SR distribution for zero GI pairs (orange histogram in Fig. 8**e**) to generate 500 samples. These samples are used to compute *T*_0.6_ (i.e. fraction exceeding 0.6). We repeat this procedure 50,000 times to estimate *P*_*T*_0.6__.

**Figure S18:**
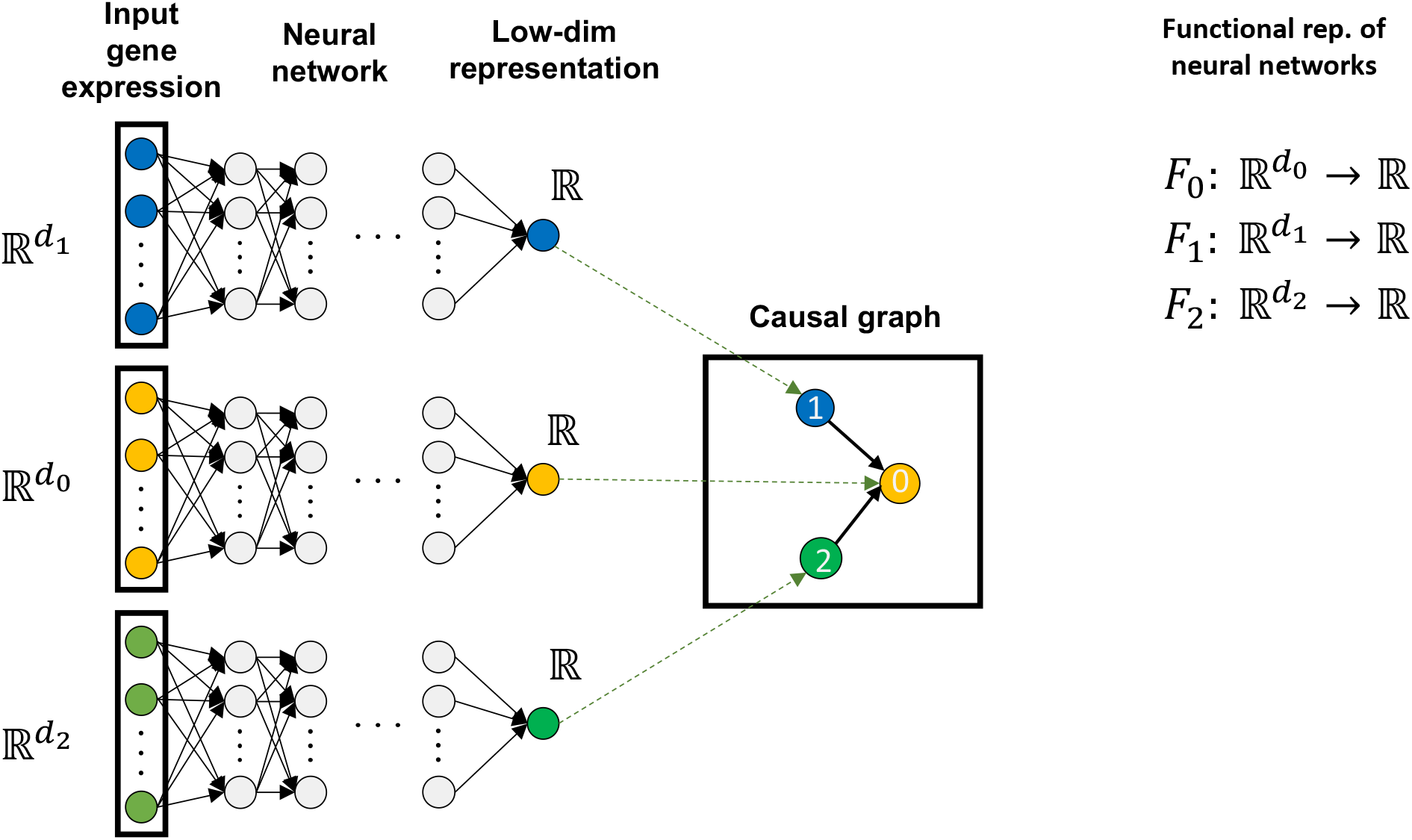
Explicit neural network architecture for models trained with the v-structed graph. Notations are defined in the main text and summarized in Table S2.

**Table S3:** KEGG pathways associated with downstream node 0 in Fig. 6a. All genes in this node are from the pathways listed in this table.

| KEGG identifier | Pathway |
| --- | --- |
| hsa04110 | Cell cycle |
| hsa04210 | Apoptosis |
| hsa04216 | ferroptosis |
| hsa04217 | necroptosis |
| hsa04115 | p53 signaling pathway |
| hsa04218 | Cellular senescence |

**Figure S19:**
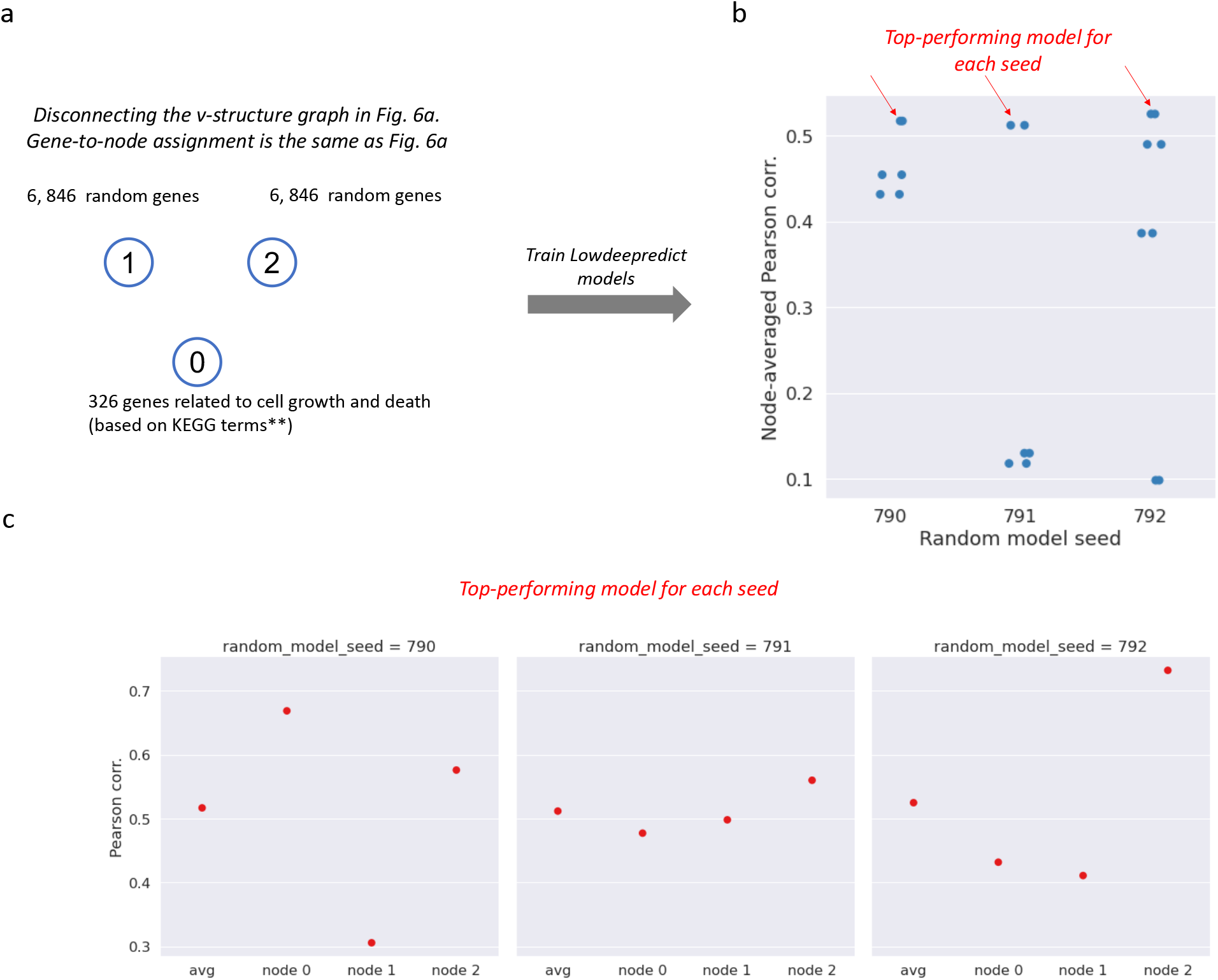
Predicted performance for models trained on disconnected v-structure graph. **a**: Causal graph is obtained by removing edges in the graph shown in Fig. 6**a**. Gene-to-node assignment is the same as in Fig. 6**a**. **b**: We train 400 Lowdeepredict models with the causal graph and gene assignment scheme described in **a**. We conduct 20 Optuna studies [27] (each containing 20 trials/models) for hyper-parameter optimization, and the node-averaged Pearson’s correlation for the best performing model for each study is shown. These results are stratified by the random seed we used to initialize neural networks (i.e. random model seed). The top-performing model under each random model seed is highlighted by a red arrow. **c**: The node-wise performance for the top-performing models highlighted in **b** is shown.

**Table S4:** KEGG pathways associated with pathway genes in Fig. 4a and 5. All *pathway genes* referenced in these figures are from the pathways listed in this table. These pathways are selected by performing functional enrichement analysis with DAVID [29] using all the perturbed genes in [5].

| KEGG identifier | Pathway |
| --- | --- |
| hsa04110 | Cell cycle |
| hsa04210 | Apoptosis |
| hsa04216 | ferroptosis |
| hsa04217 | necroptosis |
| hsa04115 | p53 signaling pathway |
| hsa04218 | Cellular senescence |
| hsa04330 | Notch signaling pathway |
| hsa05202 | Transcriptional misregulation |
| hsa04012 | ErbB signaling pathways |
| hsa04071 | Sphingolipid signaling |
| hsa05165 | Human papillomavirus infection |
| hsa04514 | Cell adhesion molecules |
| hsa04966 | Collecting duct acid secretion |
| hsa04714 | Thermogenesis |
| hsa00010 | Glycolysis / Gluconeogenesis |

**Table S5:** Size of causal subgraph based on gene sets identified by Monte Carlo. As described in FIG. 6, we apply Monte Carlo algorithm to identify 20 sets of genes (with size indicated by the *MC pathway size* column. These gene sets are first filtered to keep only the unique elements. In the *Num. unique genes* column, we list the number of unique genes for each node in the graph (Fig. 6**a**) along with the maximum number of unique genes. Note that for MC pathways of size s, the maximum number of unique genes is given by min(num. genes in each node, 20 × *s*). We then apply the causal ground-truth [28] to construct a causal subgraph *G*_mc_ using only the unique genes (*Causal subgraph* column). The size of the largest connected component of the causal subgraph is listed in the last column.

| MC pathway size, $s$ | Num. unique genes<br>(Max unique genes) | Causal subgraph $G_{mc}$ | Size of largest c.c., $s_{mc}$ |
| --- | --- | --- | --- |
| 20 | node 0: 231 (326)<br>node 1: 75 (400)<br>node 2: 63 (400) | 273 genes<br>358 edges | 162 |

## Notes

### Competing Interest Statement

C.-H.W, K.L, F.F, G.B, L.G, and J.L.E are employees of GSK. C.L contributed to this
work during his employment with GSK.

